# ProtSyntax: a protein large language model for decoding post-translational modification syntax and function

**DOI:** 10.64898/2026.07.18.739331

**Authors:** Yiyu Lin, Jiahui Wu, You Zhou, Xinye Ni, Shan Chang, Jun Ding, Yan Wang, Xin Gao, Sen Yang

## Abstract

Post-translational modifications (PTMs) expand protein function by encoding context-dependent regulatory states, and their dysregulation contributes to cancer, neurodegeneration and metabolic disease. However, existing methods treat PTMs as independent residue labels, limiting their ability to distinguish contextually permissible sites, model crosstalk and infer functional consequences. Here we introduce ProtSyntax, a PTM-aware foundation protein language model combining protein-aware positional encoding, bidirectional state-space propagation, geometry-constrained attention and adaptive multi-objective learning. This design integrates residue chemistry, motif order, long-range context and three-dimensional microenvironments while coupling PTM recognition to enzyme function. Across 40 PTM-site benchmarks, ProtSyntax exceeded the strongest baselines in mean MCC and AP by 12.66% and 10.67%. ProtSyntax also recovered masked PTM types and sites, rejected structural decoys, generalized to data-scarce modifications, reconstructed crosstalk and linked PTM perturbations to enzyme kinetics. Applications to pathogenic variants, biomolecular condensates and disease-associated PTM landscapes demonstrate its potential to decode the regulatory language of the modified proteome.

## 1. Introduction

A central challenge in modern protein science is to understand how a genetically encoded amino acid sequence is converted into a dynamic regulatory molecule inside the cell^1^. Post-translational modifications (PTMs) provide one of the most powerful mechanisms for this conversion^2^. By adding, removing or interconverting chemical groups on selected residues, PTMs expand the functional vocabulary of proteins far beyond sequence alone^3^, controlling enzymatic activity, molecular recognition^4^, localization, degradation, phase behavior and signal propagation^5^. Their importance spans nearly every major domain of biology, including cell signalling, chromatin and epigenetic regulation, metabolism, immunity, neurobiology, cancer, infection, ageing, drug discovery and protein engineering^6^. Yet PTMs are uniquely difficult to model because they are neither simple residue labels nor static sequence features^7^. Whether a residue can be modified depends on a hierarchy of constraints, including amino-acid chemistry, local motif order, enzyme-specific recognition, structural accessibility, long-range domain organization, cellular context and the modification state of neighbouring or distal residues^8^. PTM regulation can therefore be viewed as a biochemical language of the proteome: residues form the alphabet, motifs and three-dimensional microenvironments define grammar, and combinatorial modification states encode context-dependent regulatory meaning^9^. However, this language remains incompletely observed. Experimental PTM maps are sparse, condition-specific, biased toward well-studied modification types and insufficient for many rare or transient regulatory events. Thus, the key computational problem is not only to predict where PTMs occur, but to learn the rules by which proteins write, read and reinterpret modification states^10^. Even advanced proteomic assays remain constrained by detection sensitivity, incomplete site localization and limited temporal and cellular coverage, making exhaustive experimental reconstruction of PTM regulation impractical and motivating computational models that can infer unobserved events from learned biochemical principles.

Existing computational approaches have significantly advanced PTM-site annotation^11^, progressing from feature-based classifiers to deep neural networks and protein language models^12^. MusiteDeep, for example, established a general deep-learning framework that uses ensemble convolutional and capsule networks to predict and visualize multiple PTM types from local sequence windows^13^. More recently, PTMGPT2 introduced prompt-based fine-tuning of GPT-2 to exploit transferable sequence representations for PTM prediction^14^, whereas AstraPTM2 combined ESM-2 embeddings, AlphaFold2-derived structural features and a context-aware transformer to predict 39 PTM types across full-length proteins^15^. A more detailed discussion of previous PTM-prediction studies is provided in Supplementary Section 1. Collectively, these models have broadened PTM coverage, reduced dependence on handcrafted features and improved the representation of sequence and structural context. Nevertheless, their principal learning objective remains the assignment of modification labels to individual residues. Even when multiple PTM types, full-length sequences or structural features are incorporated, PTMs are generally treated as parallel prediction targets rather than as interacting elements of a shared regulatory system. This site-centric formulation leaves several fundamental biological questions unresolved. Local sequence motifs alone cannot explain why a residue is modified while an identical motif elsewhere remains inert, because modification competence can depend on solvent accessibility and the spatial convergence of sequence-distant residues into a permissive three-dimensional microenvironment^16^. Likewise, treating PTM classes as independent labels does not explicitly capture the cooperative or antagonistic crosstalk through which one modification alters the establishment, recognition or functional effect of another^17^. Most critically, predicting the mere occurrence of a PTM provides no mechanistic insight into how that specific event rewires protein function ^18^, alters enzyme kinetics, or drives disease pathogenesis^19^. Consequently, there is an urgent need for a paradigm shift—from task-specific, pattern-matching site predictors to a foundational protein language model inherently designed to learn the transferable regulatory grammar linking sequence syntax, structural geometry, and emergent biophysical consequences.

To address this conceptual gap, we introduce ProtSyntax, a foundation protein large language model specifically architected to comprehensively decode the syntax and function of PTMs. Recognizing that language models require robust, unified training corpora to learn generalized rules, we curated a comprehensive, four-million-scale PTM dataset spanning 40 modification classes, seamlessly augmented with kinase-specific phosphorylation, modification crosstalk, and enzyme kinetic supervision. Fundamentally, ProtSyntax is built upon an advanced hybrid attention and sparse Mixture-of-Experts (MoE) architecture, enabling it to efficiently scale massive model capacity while maintaining computational tractability. Embedded within this foundational macro-architecture, the core innovation of ProtSyntax lies in its specialized multi-scale design, integrating four biologically motivated modules to capture distinct hierarchical levels of PTM grammar: 1) Bio-RoPE adapts positional encoding to capture both structural periodicity and residue-dependent physicochemical modulation, embedding essential motif order and amino acid chemistry directly into the representation space; 2) Bi-Gated DeltaNet achieves highly efficient bidirectional long-context propagation, filtering out incidental local motifs by seamlessly integrating global N-terminal and C-terminal evidence around candidate residues; 3) Geometric Gated Attention (GGA) injects residue-frame-based three-dimensional constraints directly into the attention mechanism, allowing the model to strictly differentiate between sequence-compatible residues and genuinely permissive 3D structural microenvironments; and 4) PACE-Nash, an adaptive multi-objective optimization strategy, dynamically coordinates PTM classification with uncertainty-aware enzyme kinetic regression, forcing local residue-level syntax and global protein-level function to shape a shared latent representation.

By reformulating PTM prediction as a unified language-modeling problem, ProtSyntax advances beyond task-specific site annotation towards multiscale reasoning over PTM syntax and function. Across 40 PTM-site prediction benchmarks, ProtSyntax outperformed the strongest competing models, improving mean MCC and AP by 12.66% and 10.67%, respectively. More importantly, its capabilities extended beyond conventional pattern recognition: ProtSyntax recovered masked PTM types and sites in cloze-style evaluations, distinguished authentic modification sites from sequence-matched but structurally incompatible microenvironment decoys, transferred learned biochemical rules to sparsely annotated PTM classes and reconstructed cooperative and antagonistic PTM crosstalk. Through a shared multi-task encoder, the model further linked in silico PTM perturbations to directional changes in *k*_cat_, *K_m_*and *K_i_*, thereby connecting local modification syntax with protein-level functional consequences. This sequence–structure–function reasoning generalized to disease and condensate biology, enabling discrimination of pathogenic missense variants from benign polymorphisms through predicted disruption of local PTM microenvironments and capturing the nonlinear relationship between multivalent PTM patterns and liquid–liquid phase separation within intrinsically disordered regions. At pathway and therapeutic scales, ProtSyntax generated a residue-resolved PTM atlas for 383 proteins in the KEGG Alzheimer’s disease pathway, recovering established tau-associated modification landmarks while expanding the candidate PTM landscape, and integrated proteomic, phosphoproteomic and network evidence from temozolomide-resistant glioblastoma to prioritize candidate kinase–site–substrate intervention axes. Collectively, these results establish ProtSyntax as an interpretable foundation language model for decoding the regulatory syntax of the modified proteome and for quantitatively prioritizing the functional and disease-relevant consequences of PTM events.

## 2. Results

ProtSyntax processes residue-centered PTM inputs and full-length enzyme sequences through a shared sequence–structure modeling pipeline. Frozen ESM-C and SaProt embeddings are integrated with amino acid representations and AlphaFold2-derived residue geometry, then encoded by a sparse mixture-of-experts backbone combining Bio-RoPE, Bi-Gated DeltaNet and Geometric Gated Attention. Task-specific heads predict general and kinase-specific PTM sites, PTM crosstalk and enzyme kinetic parameters, while PACE-Nash coordinates these residue- and protein-level objectives within a unified representation space.

### 2.1 Establishing ProtSyntax as a PTM-aware protein language model

We first asked whether ProtSyntax learns transferable PTM syntax rather than merely fitting site-level labels. We define PTM syntax as contextual rules linking residue chemistry, motif organization, three-dimensional microenvironment and long-range context to modification compatibility, crosstalk and function. We therefore selected five complementary tests. Cloze-style recovery assesses contextual reconstruction of missing PTM types or sites; structure-aware decoys distinguish three-dimensional permissiveness from linear motif recognition; low-resource transfer measures reuse of biochemical rules across PTM classes; crosstalk reconstruction evaluates conditional dependencies between modification events; and PTM-to-function coupling tests whether local syntax is linked to protein-level functional consequences. Together, these experiments span residue-level completion, structural discrimination, cross-PTM transfer, combinatorial regulation and functional semantics, providing a multiscale evaluation of PTM-aware language learning. Finally, due to space constraints in the main text, we provide comprehensive implementation details of the experiments and a unified mechanistic explanation in Supplementary Section 4.7.

#### 2.1.1 ProtSyntax recovers residue-level PTM syntax by cloze-style inference

To examine whether ProtSyntax can infer missing modification information from residue-level context, we formulated PTM prediction as two cloze-style language tasks. In the PTM-type recovery task^20^, the model was provided with a candidate residue, its flanking sequence context and available structural information, and was asked to recover the correct PTM type among 40 modification classes. In the PTM-site recovery task, the model was given a PTM type and a protein segment containing multiple chemically compatible residues and was required to rank the experimentally modified residue above unmodified candidates. These tasks mimic masked-token recovery in language modeling but replace words with biochemical modification events^21^. The comprehensive implementation details of the experiments are provided in Section 4.7.2 of the supplementary material.

ProtSyntax achieved the strongest performance across both evaluation settings (Figure 2.A). In PTM-type recovery, ProtSyntax reached an accuracy of 0.842, outperforming ESM-C+MLP (fine-tuned), SaProt+MLP (fine-tuned) and AstraPTM2, which achieved accuracies of 0.741, 0.766 and 0.719, respectively. Ablation of Bio-RoPE reduced the accuracy to 0.801, indicating that PTM-aware positional encoding contributes to the recovery of modification-type syntax. In PTM-site recovery, ProtSyntax achieved a site-ranking AUPRC of 0.812, exceeding ESM-C+MLP, SaProt+MLP and AstraPTM2 by 12.3, 9.6 and 8.2 percentage points, respectively. This improvement was most pronounced in fragments containing multiple chemically compatible residues, suggesting that ProtSyntax identifies the contextually favored regulatory residue rather than simply detecting residue-level permissibility.

**Figure 1.**
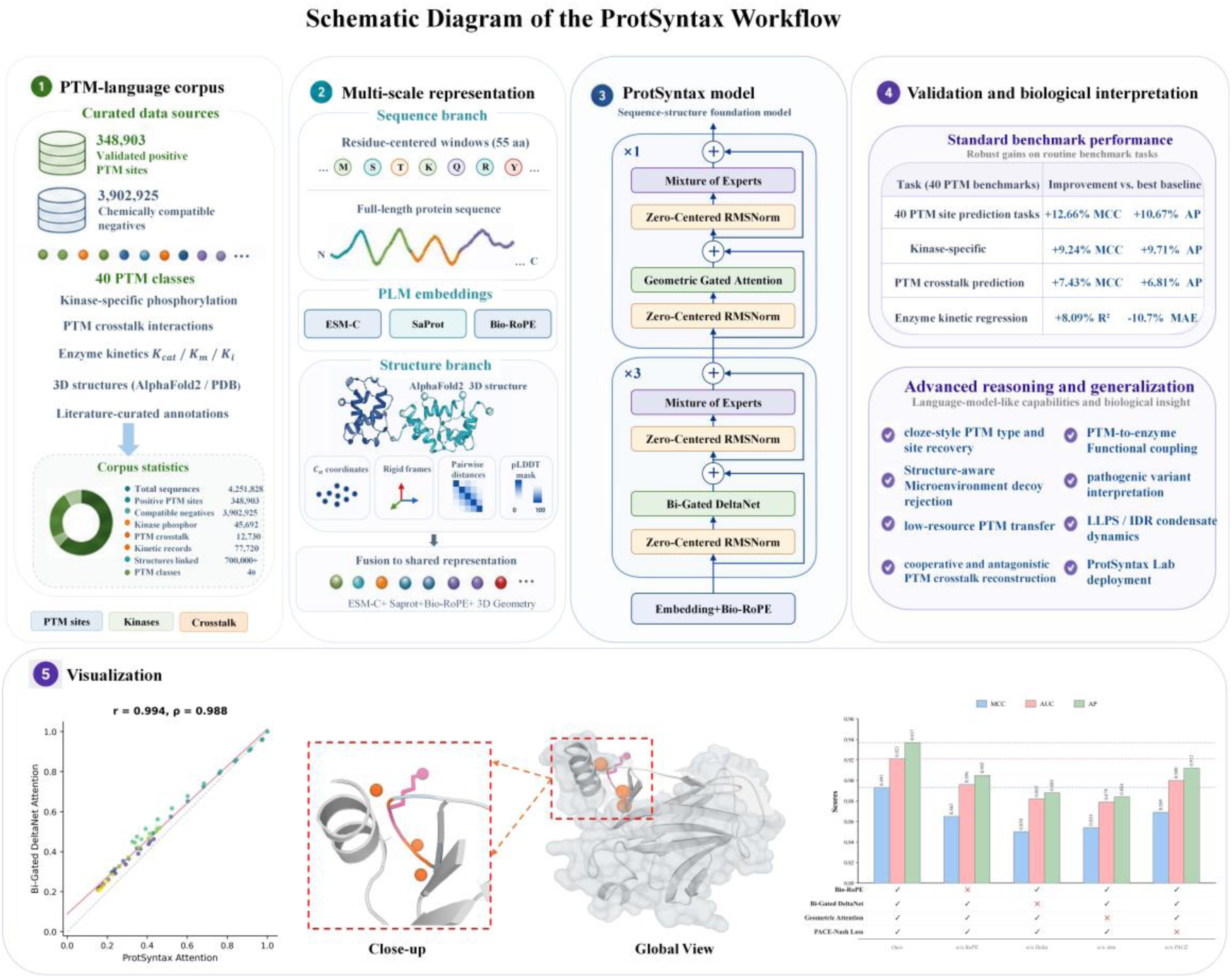
Overview of ProtSyntax. Experimentally supported PTM annotations, kinase– substrate relationships, PTM-crosstalk records and enzyme-kinetic measurements are converted into task-specific sequence and structure inputs. Frozen ESM-C and SaProt representations and AlphaFold2-derived residue geometry are integrated by a shared sparse mixture-of-experts encoder containing Bio-RoPE, Bi-Gated DeltaNet and Geometric Gated Attention blocks. PACE-Nash coordinates residue-level classification and protein-level regression objectives. Evaluation comprises matched benchmarks, contextual and geometric representation tests, low-resource transfer, in silico perturbation analyses and biological case studies.

**Figure 2.**
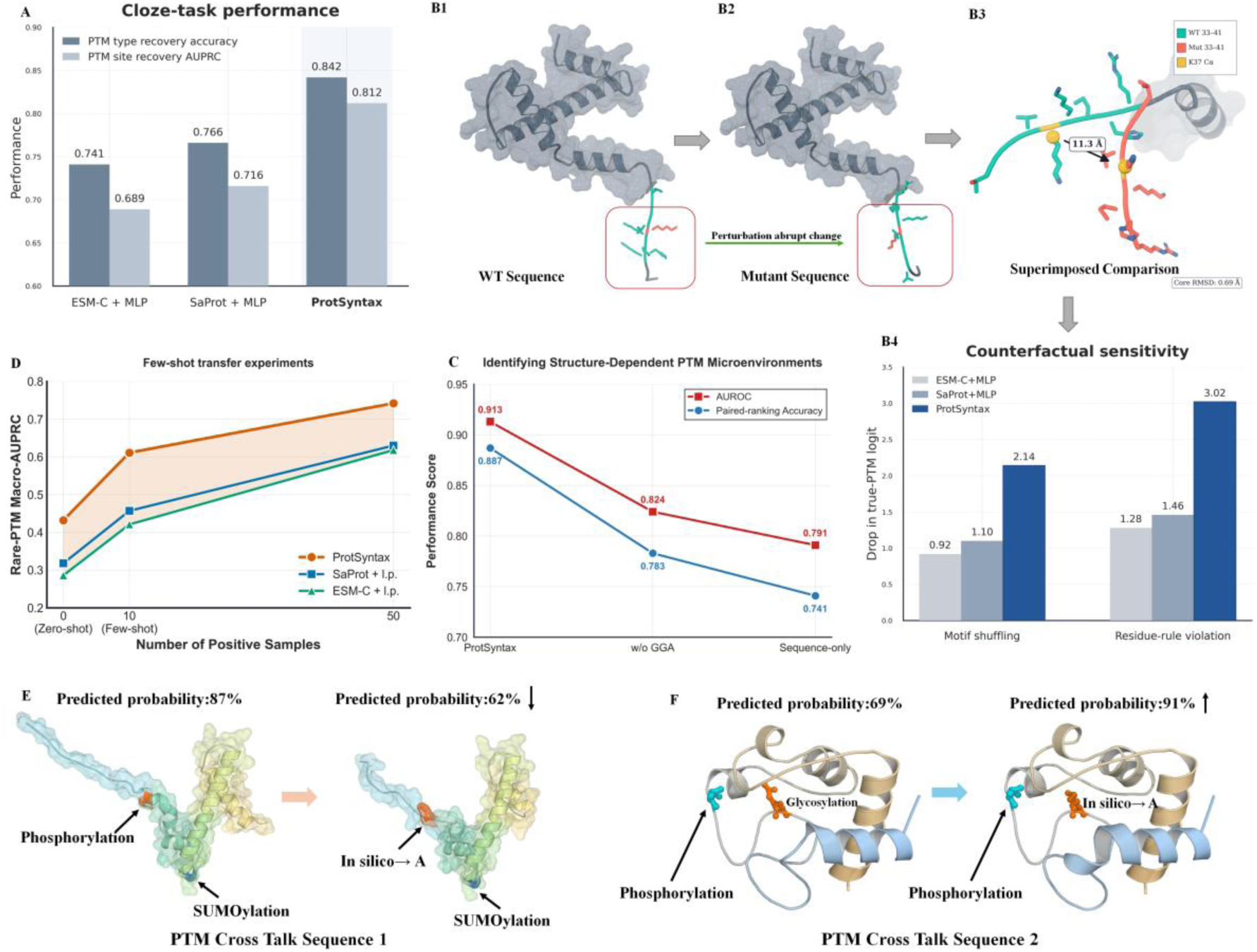
Validation of the ProtSyntax learning paradigm. (A) Performance comparison on the PTM language cloze test. (B1-B4) Counterfactual perturbation analysis, detailing protein structural comparisons and corresponding quantitative results. (C) Structural visualizations and quantitative analysis demonstrating the identification of structure-dependent PTM microenvironments by ProtSyntax. (D) Few-shot transfer learning performance of ProtSyntax on low-resource PTM types. (E-F) Comparative analysis illustrating the successful reconstruction of combinatorial PTM crosstalk grammar.

To distinguish PTM syntax learning from shallow motif recognition, we introduced two counterfactual perturbation settings. Motif shuffling disrupted residue order while approximately preserving amino acid composition, whereas residue-rule violation replaced the central residue with an amino acid chemically incompatible with the target PTM (Figure 2.B1–B4). Both perturbations induced larger decreases in the true-PTM logit for ProtSyntax than for ESM-C+MLP (fine-tuned), SaProt+MLP (fine-tuned) or AstraPTM2. These results indicate that ProtSyntax is sensitive to the joint constraints among residue identity, motif order and PTM type, supporting its role as a PTM-aware protein language model rather than a conventional site-level classifier.

#### 2.1.2 ProtSyntax resolves structure-dependent PTM microenvironments

PTM site formation is often determined by spatially proximal residues, conformational accessibility and domain context in addition to linear motifs^8^. We therefore designed a structure-aware microenvironment decoy test to evaluate whether ProtSyntax uses structural information in a functionally selective manner. For each experimentally validated PTM site, we constructed three classes of paired decoys: sequence-matched decoys with similar local motifs but distinct three-dimensional neighborhoods, structure-matched decoys with similar local spatial environments but incompatible residue chemistry or motifs, and random same-residue decoys from unmodified positions in the same protein. The model was required to rank the validated PTM site above its paired decoys. The comprehensive implementation details of the experiments are provided in Section 4.7.2 of the supplementary material.

ProtSyntax effectively distinguished true PTM sites from structural decoys, achieving a paired-ranking accuracy of 0.887, an AUROC of 0.913 and an AUPRC of 0.786 (Figure 2.C). These values exceeded those of the strongest competing model, AstraPTM2, by 6.2, 6.7 and 7.3 percentage points, respectively. Removing Geometric Gated Attention reduced the paired-ranking accuracy to 0.783 and the AUROC to 0.824, while the sequence-only variant showed a further decline in performance. The largest improvement was observed for sequence-matched decoys, indicating that explicit geometric modeling enables ProtSyntax to reject motif-like false positives that lack a permissive structural microenvironment. Coordinate perturbation analysis further showed that ProtSyntax was robust to mild structural noise but sensitive to disruption of residue-frame correspondence, supporting the conclusion that the model uses genuine three-dimensional microenvironment information rather than nonspecific structural cues.

#### 2.1.3 ProtSyntax transfers PTM syntax to low-resource modification types

A PTM language model should reuse biochemical rules learned from abundant PTM classes to recognize sparsely annotated modification types. We evaluated this property using a leave-one-PTM-type-out transfer setting. Each low-resource PTM type, including lactoylation, formylation, butyrylation, N-palmitoylation, C-linked glycosylation, sulfoxidation, nitration and dephosphorylation, was removed during pretraining and subsequently evaluated under zero-shot representation and few-shot adaptation settings with 0, 10 or 50 positive samples. The comprehensive implementation details of the experiments are provided in Section 4.7.2 of the supplementary material.

ProtSyntax exhibited substantially higher sample efficiency than fine-tuned ESM-C+linear probe, SaProt+linear probe and PTMGPT2 (Figure 2.D). In the zero-shot setting, ProtSyntax achieved a rare-PTM macro-AUPRC of 0.432, compared with 0.286, 0.318 and 0.274 for ESM-C, SaProt and PTMGPT2, respectively. With only 10 positive samples, ProtSyntax improved to 0.611, whereas ESM-C, SaProt and PTMGPT2 reached 0.421, 0.457 and 0.402, respectively. With 50 positive samples, ProtSyntax reached a macro-AUPRC of 0.742, approaching the fully supervised regime. Ablation of PACE-Nash impaired performance under both few-shot settings, suggesting that physicochemical contrastive regularization and multi-objective coordination facilitate transfer across PTM chemistries. Representation analysis further showed that ProtSyntax organizes low-resource PTMs according to biochemical relationships rather than label frequency alone. Lysine acylation-related PTMs occupied neighboring regions in the latent space, enabling rules learned from acetylation, succinylation and crotonylation to transfer to less frequent acylation events. Similar transfer was observed among cysteine-centered PTMs, whereas transfer was weaker between chemically unrelated modification classes. These findings support the interpretation that ProtSyntax learns a chemically structured PTM syntax. Detailed prediction results for each rare PTM type are provided in Supplementary Section 4.3.

#### 2.1.4 ProtSyntax reconstructs combinatorial PTM crosstalk grammar

PTM regulation is frequently combinatorial: one modification can promote, inhibit or rewire the probability of another depending on sequence and structural context^22^. To test whether ProtSyntax captures such higher-order syntax, we performed in silico perturbation experiments on unseen proteins. Candidate regulatory sites were substituted with alanine, and the resulting changes in the predicted probabilities of associated PTM events were quantified to infer cooperative or antagonistic dependencies (Figure 2.E-F). The comprehensive implementation details of the experiments are provided in Section 4.7.2 of the supplementary material.

In a protein segment containing coupled phosphorylation and SUMOylation events, alanine substitution of the phosphorylation site reduced the predicted SUMOylation probability from 0.87 to 0.62, consistent with a cooperative dependency. Conversely, perturbing a glycosylation site increased the predicted phosphorylation probability from 0.69 to 0.91, consistent with an antagonistic relationship. To further assess whether these perturbation responses generalize beyond individual examples, we curated 100 experimentally annotated PTM crosstalk sites from the PTMcode v2 dataset and applied the same in silico knockout strategy. ProtSyntax correctly recovered the expected crosstalk relationship for 91% of these sites, substantially outperforming DeepPCT (73%) and ProXTalk (70%). Together, these examples show that ProtSyntax does not treat PTM events as independent labels; instead, it encodes conditional dependencies among modification events and responds to perturbations in a manner consistent with combinatorial PTM regulation.

#### 2.1.5 ProtSyntax links local PTM syntax to enzyme functional consequences

We next asked whether the PTM syntax learned by ProtSyntax is linked to protein-level functional consequences. This analysis exploited the shared encoder used for residue-level PTM prediction and full-length enzyme kinetic regression, which allows local modification representations to be trained together with global functional readouts^23^. We curated enzymes with experimentally annotated or high-confidence PTM sites and grouped them according to the spatial relationship between the PTM site and functional regions, including catalytic neighborhoods, substrate-binding pockets, putative allosteric regions and distal solvent-exposed surfaces. We then performed in silico modification-blocking or modification-mimetic substitutions and compared the resulting changes in ProtSyntax PTM scores with changes in K_cat_, K_m_ and K_i_. The comprehensive implementation details of the experiments are provided in Section 4.7.2 of the supplementary material.

ProtSyntax-predicted ΔPTM scores were strongly associated with perturbations in enzyme kinetics. Among enzymes with available kinetic annotations, ΔPTM scores correlated with Δlog Kcat, Δlog Km and Δlog Ki, with Spearman’s ρ values of 0.628, 0.547 and 0.512, respectively. These correlations exceeded those achieved by CatPred (0.501, 0.422 and 0.395) and DKEP (0.477, 0.405 and 0.356). PTM perturbations near catalytic residues or substrate-binding pockets also produced substantially larger predicted functional effects than distal, solvent-exposed perturbations, with mean predicted Δlog Kcat values of 0.61 and 0.18, respectively. By comparison, the corresponding values were 0.455 and 0.14 for CatPred and 0.418 and 0.13 for DKEP. Ablation of the enzyme kinetic regression objective reduced the correlation between ΔPTM scores and Δlog Kcat to 0.421, whereas further removal of PACE-Nash loss decreased it to 0.386, indicating that joint kinetic supervision and coordinated multi-objective optimization promote representations that couple PTM states to functional consequences. ProtSyntax-derived uncertainty estimates were also consistent with the reliability of its functional perturbation predictions. Within the high-confidence subset, ProtSyntax correctly predicted the direction of Kcat changes with an accuracy of 0.781, compared with 0.594 in the low-confidence subset. In contrast, CatPred achieved accuracies of 0.671 and 0.452, whereas DKEP achieved 0.623 and 0.401, respectively. When the analysis was restricted to PTM sites near catalytic pockets, substrate-binding regions or annotated regulatory domains, the directional accuracy of ProtSyntax increased further to 0.823, compared with 0.698 for CatPred. Collectively, these results demonstrate that ProtSyntax extends beyond local PTM site recognition to quantitatively infer the functional consequences of modification events, thereby linking PTM syntax to protein functional regulation.

We further demonstrate the advantages of ProtSyntax in predicting highly evolutionarily conserved protein sequences and protein sequences harboring combinatorial PTM sites in the Supplementary Materials, with detailed results provided in Supplementary Section 4.4.

### 2.2 ProtSyntax outperforms task-specific state-of-the-art models across PTM and enzyme benchmarks

We benchmarked ProtSyntax against recent task-specific models across general PTM site prediction, kinase-specific phosphorylation, PTM crosstalk prediction and enzyme kinetic regression. Baselines included PTMGPT2, PTM-Mamba, MTPrompt-PTM and AstraPTM2 for general PTM prediction, DCPPS^24^ for kinase-specific phosphorylation, DeepPCT^25^ for PTM crosstalk and CatPred^26^ for enzyme kinetic parameters. All models were trained and evaluated using matched data partitions and evaluation protocols. ProtSyntax consistently improved over the strongest baseline across classification and regression tasks. Across 40 PTM site prediction tasks, ProtSyntax increased mean MCC and AP by 12.66% and 10.67%, respectively, relative to the runner-up model. For four kinase-specific phosphorylation tasks, ProtSyntax improved mean MCC and AP by 9.24% and 9.71%, and for PTM crosstalk prediction it improved MCC and AP by 7.43% and 6.81%. For enzyme kinetic regression, ProtSyntax increased mean R² by 8.09% and reduced MAE by 10.7%. The detailed performance metrics of the comparative models for each task are reported in Tables S7–S14 in Supplementary Section 6.

These improvements across heterogeneous PTM and enzyme benchmarks indicate that ProtSyntax benefits from a unified PTM-language representation rather than from optimization for a single task. Finally, we comparatively evaluated the prediction speeds of ProtSyntax and other models across three diverse scenarios, with detailed results presented in Supplementary Section 4.2.

**Table 1.**
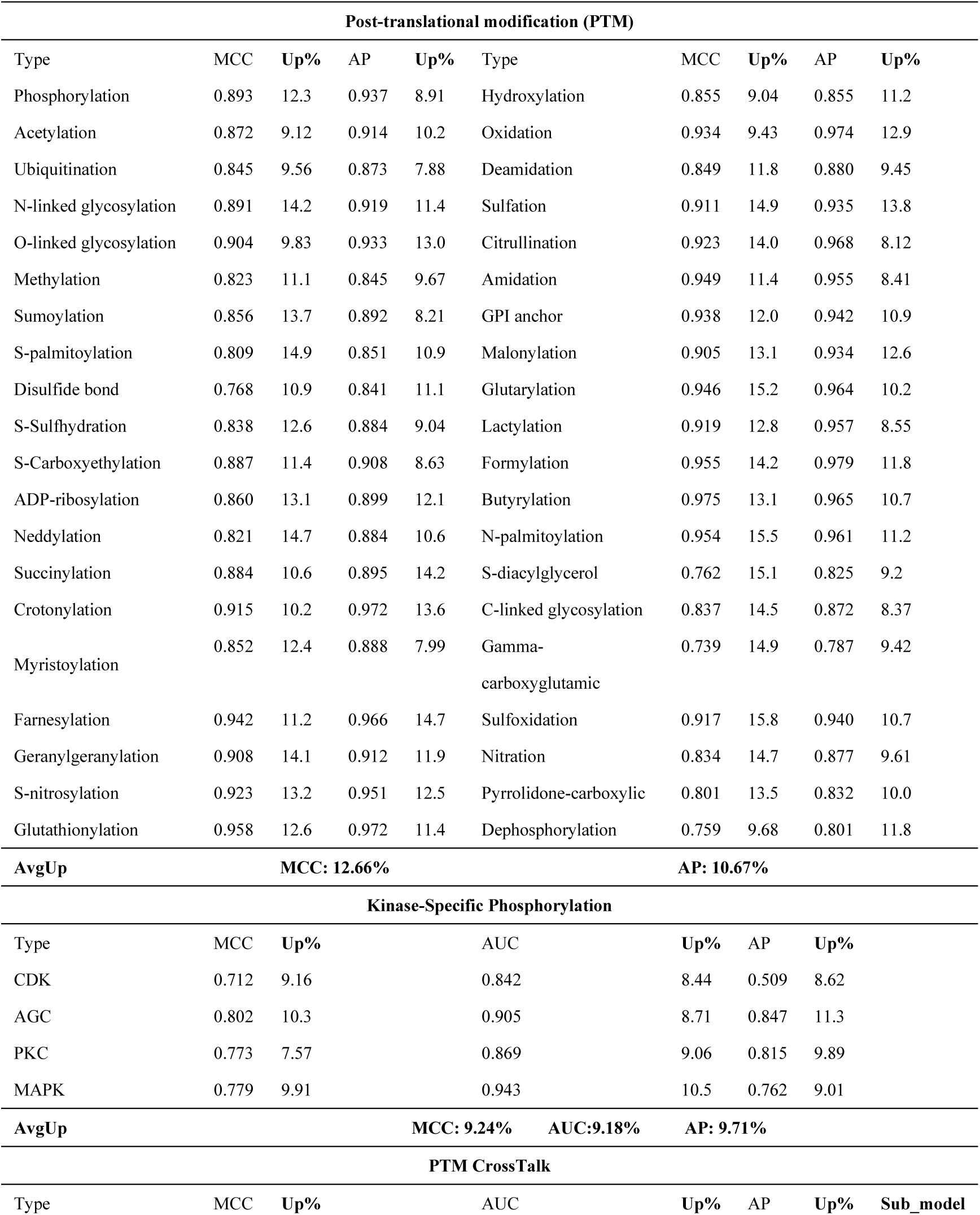
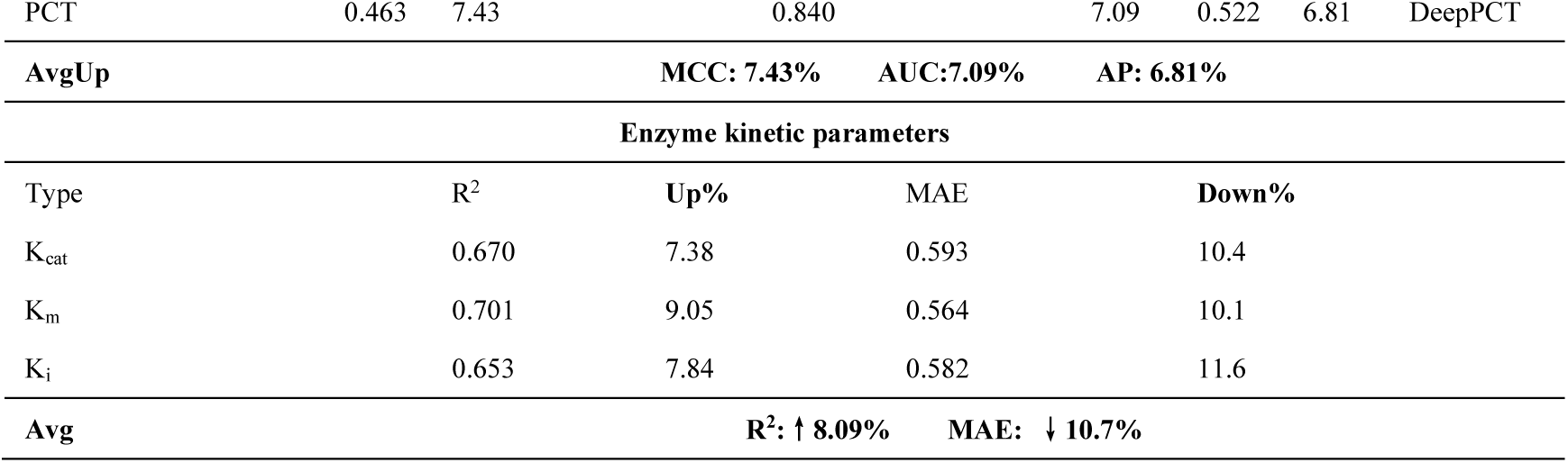
Performance comparison across large-scale benchmarks. The ‘Up’ and ‘Down’ values denote the relative percentage improvement and reduction, respectively, in ProtSyntax’s performance metrics compared to the runner-up model. For brevity and visual clarity, this condensed table presents only the absolute performance of ProtSyntax alongside its relative margins. Details regarding the specific second-best baseline models for each task are described in the main text.

### 2.3 Disease-associated case studies of ProtSyntax

To determine whether the learned PTM syntax extends to disease-associated network rewiring, we tasked ProtSyntax with zero-shot variant effect prediction^27^. Using clinical missense and somatic cancer mutations (ClinVar/COSMIC) paired with benign polymorphisms (gnomAD), we performed in silico mutagenesis and local structural relaxation on variants near validated PTM sites (see Figure 3.A-D). ProtSyntax accurately discriminated pathogenic from neutral variants strictly via local PTM syntax disruption (AUROC = 0.885, AUPRC = 0.812), surpassing the comparative model PTMGPT2 (AUROC = 0.801, AUPRC = 0.729). Crucially, by leveraging the spatial sensitivity of the Geometric Gated Attention (GGA) module, the model identified mutations that leave the modifiable residue chemically intact but sterically abolish its 3D permissive microenvironment. For example, in a simulated PTEN tumor suppressor mutant, a spatially adjacent but sequentially distant substitution induced a severe predicted loss of ubiquitination (Δ*P_ptm_* = −0.47). The corresponding GGA spatial penalty increased 3.2-fold, mechanistically mapping the mutation to a steric occlusion of the E3 ligase interface. Thus, ProtSyntax directly translates variant-induced geometric perturbations into interpretable, residue-level regulatory consequences. The comprehensive implementation details of the experiments are provided in Section 4.7.3 of the supplementary material.

**Figure 3.**
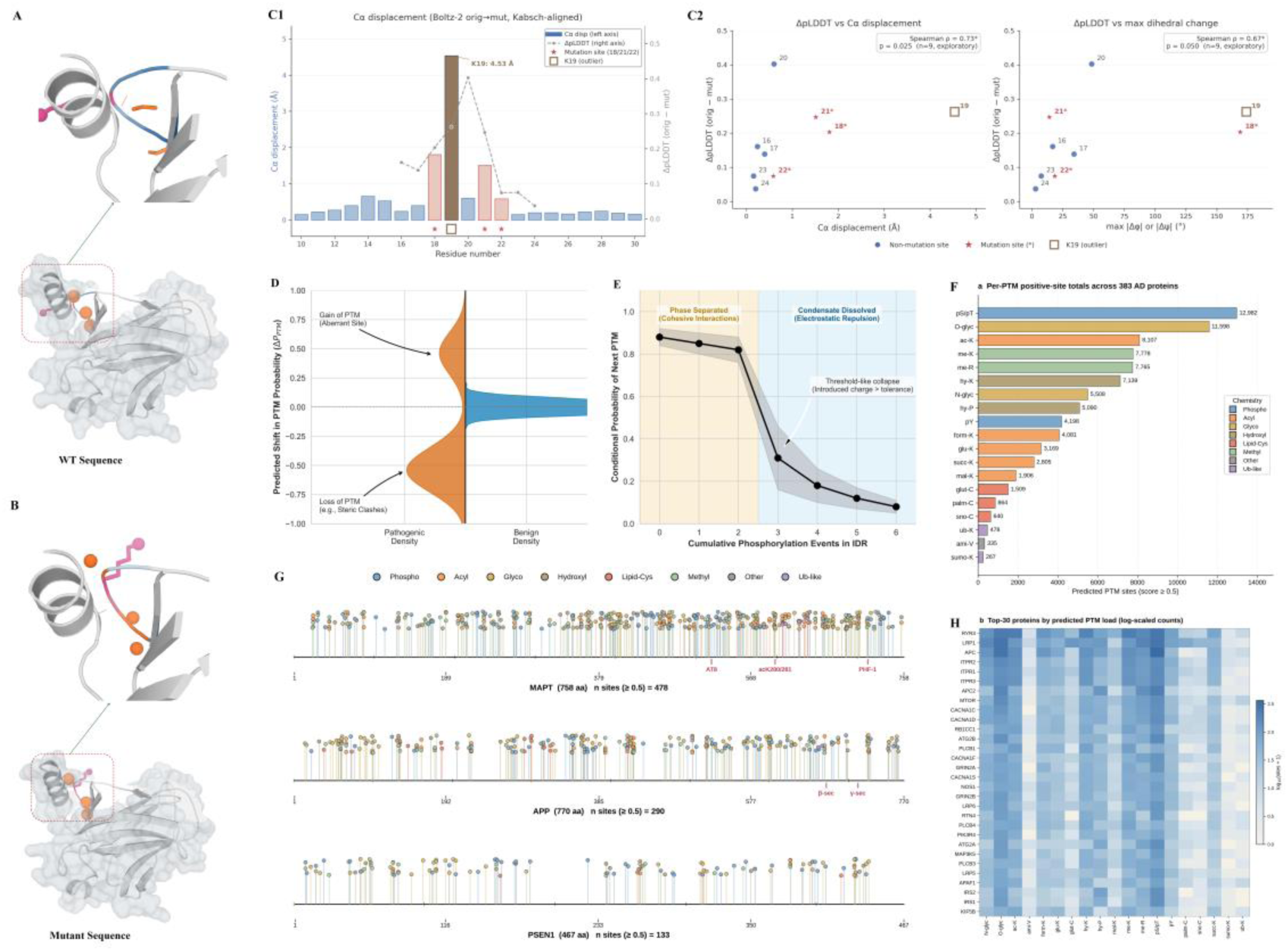
Hypothesis-generating applications of ProtSyntax. (A, B) Reference and mutant structures for the representative PTM-proximal variant, with the perturbed neighbourhood enlarged. (C1-C2) Relationship between local coordinate displacement and the reported structural-confidence change after the mutant-structure procedure. (D) Distribution of PTM-score changes for pathogenic or cancer-associated variants and matched benign variants. (E) Conditional probability of an additional modification across the in-silico FUS phosphorylation series; background shading denotes low- and high-phosphorylation regimes used for visualization and does not represent experimentally established phases. (F) Numbers of positive-scoring residue–PTM pairs across 383 KEGG Alzheimer’s disease pathway proteins at a score threshold of 0.5. (G) Predicted PTM landscapes for MAPT, APP and PSEN1; curated landmarks are indicated for reference. (H) Length-unadjusted PTM-score counts for the 30 proteins with the largest predicted burden across 19 classes.

We next asked whether ProtSyntax could resolve the combinatorial PTM grammar underlying highly dynamic biomolecular processes, using liquid–liquid phase separation (LLPS) as a representative case^28^. Biomolecular condensates are often regulated by multivalent PTMs within intrinsically disordered regions (IDRs). Because such regions generally lack reliable tertiary structural constraints, we masked residues with low-confidence AlphaFold coordinates (pLDDT < 50), allowing ProtSyntax to operate in a sequence-dominant mode driven primarily by the Bi-Gated DeltaNet module. Across validated LLPS-associated proteins, iterative in silico modification analysis showed that the ProtSyntax-predicted multi-site modification propensity was strongly associated with experimentally measured shifts in saturation concentration (C_sat_, *Spearman_ρ_* = −0.71, *p* < 10^−4^), substantially outperforming the strongest comparator, PTMGPT2 (*Spearman_ρ_* = −0.438, *p* < 10^−2^). In a simulated hyperphosphorylation trajectory of the FUS RNA-binding domain, analysis of Bi-Gated DeltaNet cross-gating weights revealed a system-level, phase-like transition. As serine and threonine residues were sequentially modified, model attention progressively shifted from cohesive short-range clustering toward long-range electrostatic repulsion. Notably, the introduction of a third proximal phosphorylation event induced a threshold-like collapse in the conditional probability of subsequent modifications, from 0.82 to 0.31 (Figure 3.E), whereas the strongest comparator, MTPrompt-PTM, showed only a modest decrease from 0.70 to 0.55. These results indicate that ProtSyntax captures nonlinear PTM-dependent regulatory logic associated with condensate behavior, even when inference is driven predominantly by learned sequence syntax rather than explicit structural information. The comprehensive implementation details of the experiments are provided in Section 4.7.3 of the supplementary material.

As an additional disease-focused case study, we used ProtSyntax to construct a residue-level PTM atlas for 383 reviewed human proteins in the KEGG Alzheimer’s disease pathway (hsa05010; Figure 3F–H). Across 214,479 chemically compatible residue– PTM pairs spanning 19 modification classes, ProtSyntax identified 86,219 sites with scores ≥0.5, including 18,011 stringent predictions (score ≥0.8), with at least one predicted site in every pathway protein. The atlas recapitulated established AD biology: functional enrichment converged on autophagy, synaptic signalling, calcium regulation and PI3K–AKT pathways, while kinase-profile analysis revealed marked enrichment of tau-associated CMGC kinases among predicted MAPT phosphosites. Comparison with 10,507 curated UniProt and PhosphoSitePlus sites showed strong recovery across major PTM classes and key proteins, including the AT8 and PHF-1 tau phospho-epitopes and the K280/K281 acetylation cluster. Moreover, ten PTM classes largely absent from previous prediction frameworks contributed 23,297 additional candidates. These results establish a chemically broad, experimentally anchored AD PTM landscape that can prioritize sites for targeted proteomics, mutagenesis and therapeutic investigation. Further details are provided in Supplementary Section 4.5.

As a complementary case study, we demonstrate that ProtSyntax can translate disease-associated PTM landscapes into actionable therapeutic hypotheses by prioritizing kinase–site–substrate intervention axes in temozolomide-resistant glioblastoma with full analyses provided in Supplementary Section 4.6.

Finally, ProtSyntax Lab provides an open-source bilingual web interface for PTM prediction and protein language model analysis; details are provided in Supplementary Section 4.9.

### 2.4 Component ablation and robustness to uncertain structures

To assess the contribution of key ProtSyntax components, we performed ablation experiments on the phosphorylation dataset, focusing on Bio-RoPE, Bi-Gated DeltaNet, Geometric Gated Attention and PACE-Nash loss. As shown in Figure 4.A-B, replacing conventional modules commonly used in large language models, such as standard RoPE and Gated DeltaNet, with ProtSyntax-specific designs consistently improved performance. Bio-RoPE, Bi-Gated DeltaNet, Geometric Gated Attention and PACE-Nash loss increased MCC by 2.8%, 4.3%, 3.9% and 2.4%, respectively. To assess robustness to uncertain structures, we stratified the phosphorylation test set by pLDDT (<50, 50–70 and >70). ProtSyntax consistently outperformed PTMGPT2 across all strata, retaining a 12.6% MCC advantage even at pLDDT <50 (Figure 4.C), demonstrating robust integration of sequence and structural context (Supplementary Section 4.1).

**Figure 4.**
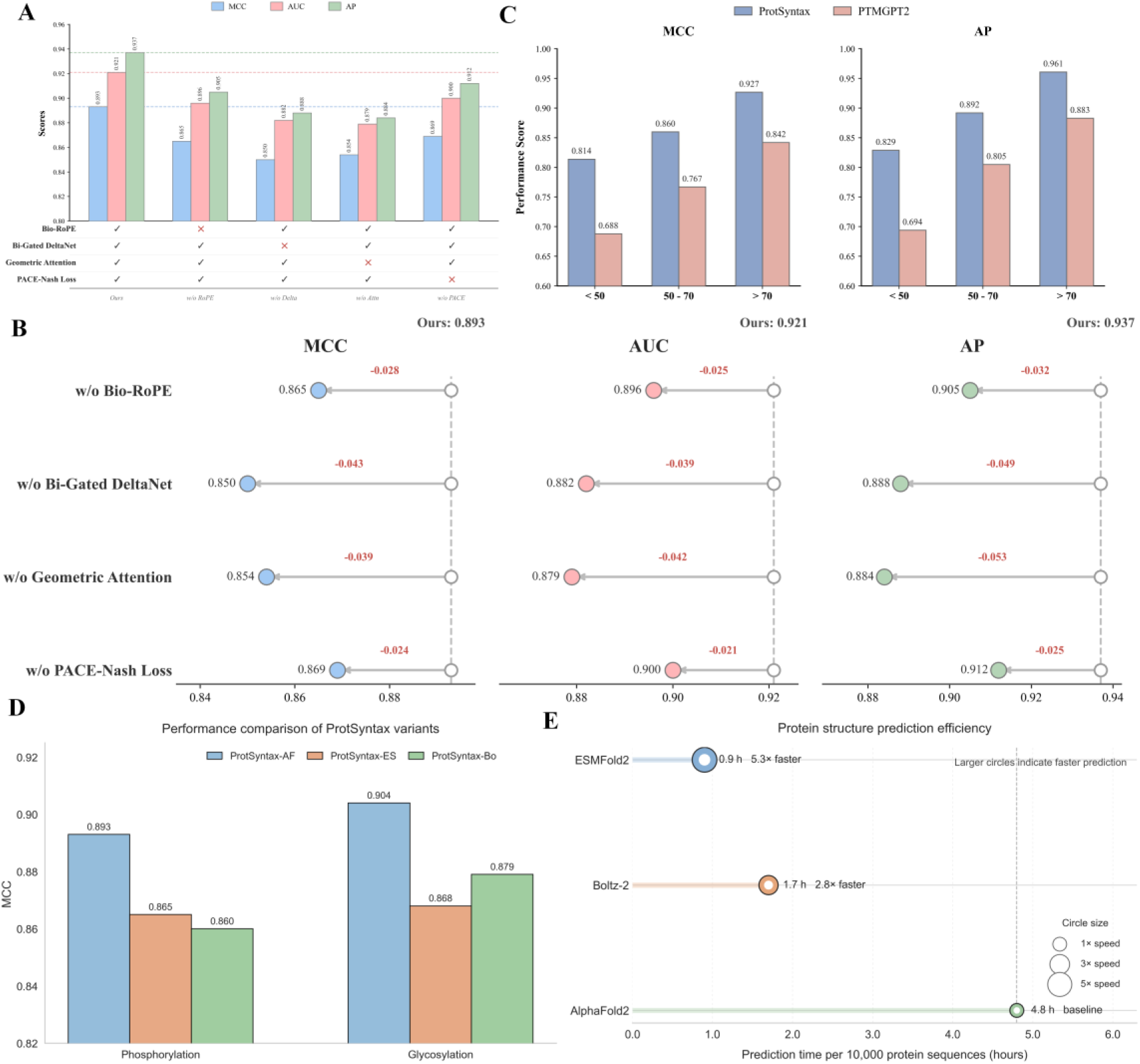
Visualizations of ablation and generalization experiments. (A–B) Visualizations of the ablation study on four proposed modules. (C) Performance comparison of ProtSyntax under varying protein pLDDT scores. (D–E) MCC comparison and prediction-speed analysis of ProtSyntax variants using AlphaFold2, ESMFold or Boltz-2 as the structure-generation backbone.

We also compared structures generated by AlphaFold2, ESMFold and Boltz-2. AlphaFold2-derived inputs produced the highest MCC on the phosphorylation and glycosylation benchmarks, whereas ESMFold and Boltz-2 reduced structure-generation time under the stated hardware conditions. This comparison defines an accuracy– efficiency trade-off rather than identifying a universally optimal structure generator (Figure 4.E).

### 2.5 Interpretable local–global reasoning in ProtSyntax

ProtSyntax integrates residue-level PTM syntax with protein-scale sequence and structural context^29^. As shown in Figure 5, Bi-Gated DeltaNet propagates bidirectional long-range information to distinguish coherent PTM-associated patterns from incidental local motifs, whereas Geometric Gated Attention constrains residue interactions using three-dimensional proximity. Their 3:1 interleaving enables ProtSyntax to infer PTM susceptibility from the joint effects of local chemistry, distal sequence context and structural–functional organization. A detailed analysis of model interpretability is provided in Supplementary Section 4.8.

**Figure 5.**
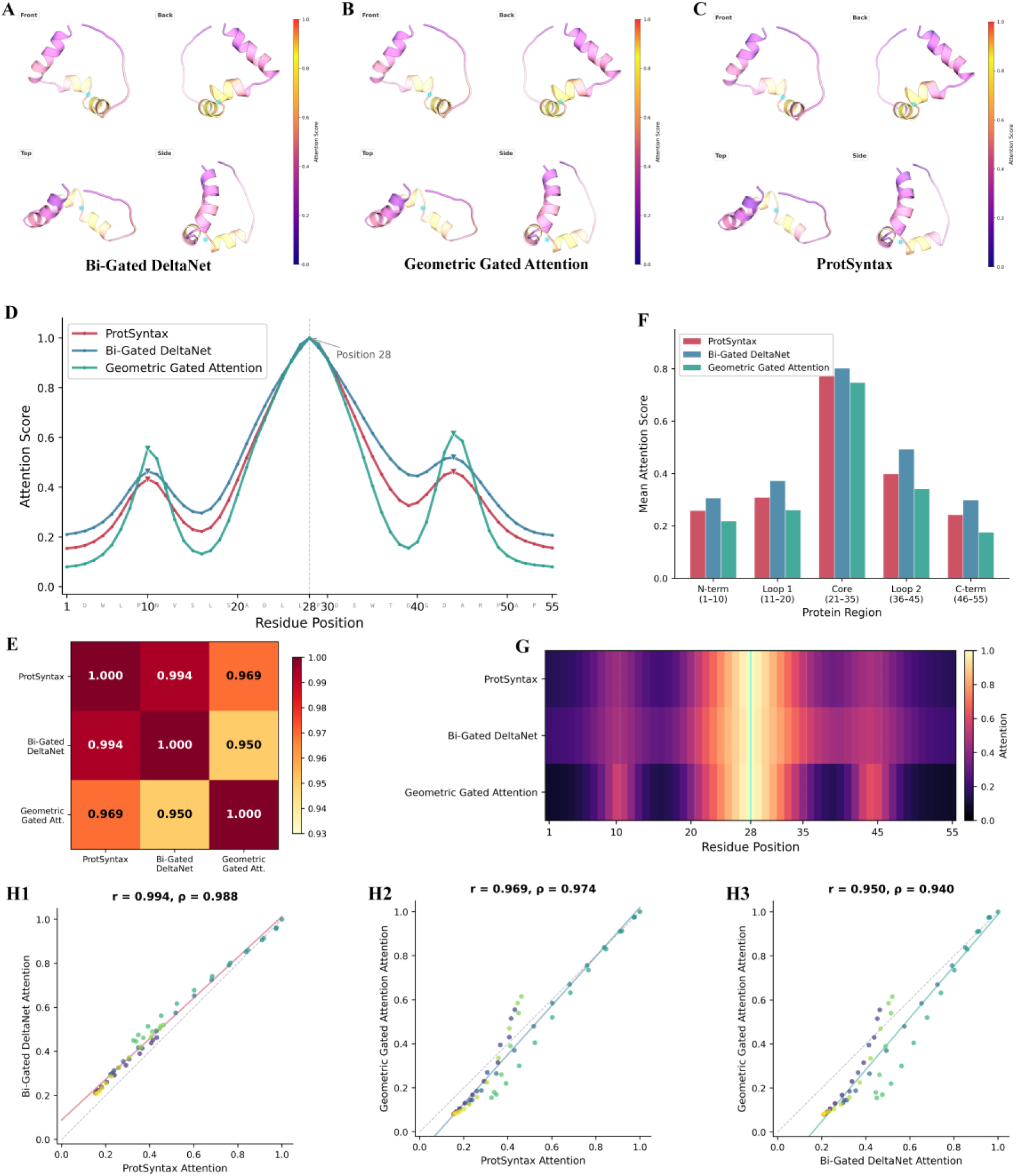
Residue-level attention-pattern analysis. (A-C) Visual comparison of attention scores for ProtSyntax, Bi-Gated DeltaNet, and Geometric Gated Attention mapped onto the protein structure graphs. (D) Attention Score Profiles. Residue-wise distribution of attention scores for each module along the target protein sequence. (E) Pearson Correlation. Matrix of Pearson correlation coefficients evaluating the relationships among the attention scores of the three modules. (F) Regional Mean Attention. Average attention scores aggregated by distinct protein structural regions (N-terminus, Loop 1, Core, Loop 2, and C-terminus). (G) Residue-Level Attention Heatmap. Heatmap illustrating the residue-wise attention scores across the three modules. (H1-H3) Pairwise Scatter Plots. Scatter plots comparing residue-wise attention scores between module pairs. Data points are color-coded according to sequence position (gradient from purple at the N-terminus to yellow at the C-terminus). The diagonal dashed line represents the reference line of perfect agreement, and the solid line indicates the linear regression fit.

## 3. Discussion

ProtSyntax reframes PTM modeling from independent site annotation to multiscale regulatory inference. Here, PTM syntax denotes conditional rules whereby residue chemistry and motif order define local compatibility, long-range sequence context and three-dimensional geometry determine structural permissiveness, and interacting modifications shape functional consequences. By learning these dependencies jointly, ProtSyntax treats site selection, PTM identity, crosstalk and function as coupled aspects of a context-dependent regulatory event, thereby bridging proteome-scale PTM mapping with biological interpretation.

ProtSyntax implements this formulation through Bio-RoPE for PTM-aware motif organization, Bi-Gated DeltaNet for bidirectional long-range context, Geometric Gated Attention for residue-frame structural constraints and PACE-Nash for coordinating PTM learning with uncertainty-aware kinetic supervision. Evidence for transferable PTM syntax extends beyond benchmark gains: ProtSyntax recovered masked PTM information, rejected structurally invalid motif decoys, transferred to low-resource modifications, reconstructed conditional crosstalk and linked predicted PTM perturbations to kinetic changes. These complementary results argue against simple motif memorization or task-specific fitting. Applications to disease variants and condensate regulation further illustrate its capacity to generate hypotheses across sequence, structure and function. Thus, the principal contribution of ProtSyntax is not merely broader PTM coverage, but a unified framework for inferring where modifications occur, the contexts that permit them and their potential functional consequences. An extended discussion is provided in Supplementary Section 7.

Despite these advances, several limitations remain. ProtSyntax is inherently constrained by the incompleteness and experimental bias of current PTM annotations, particularly for rare, transient and condition-specific modifications^30^. Furthermore, although the geometric branch enables structure-aware reasoning, it still relies on predicted or static protein conformations and therefore cannot fully represent conformational ensembles, enzyme–substrate encounter states or highly dynamic molecular environments^31^. Likewise, the current framework links PTM syntax to function primarily through enzyme kinetic supervision and in silico perturbation, but does not yet explicitly model PTM occupancy, temporal signaling dynamics, proteoform diversity or cell-type-specific regulation. Future work should therefore integrate quantitative and time-resolved proteomics, perturbation datasets, interaction networks and dynamic structural information to enable context-dependent modeling of PTM regulation^32^. More broadly, PTM modeling should evolve from predicting whether a residue can be modified to understanding when, where, under which regulatory context and with what functional consequence modification occurs^33^. By framing PTM biology as a foundation-model problem spanning protein sequence, structure and function, ProtSyntax provides a scalable and interpretable framework for decoding the regulatory syntax of the modified proteome and for generating mechanistic hypotheses in protein engineering, functional proteomics and precision medicine.

## 4. Methods

### 4.1 PTM-language corpus construction and task formulation

To operationalize PTM language learning, we constructed a multi-task protein corpus in which each supervision source captures a distinct level of PTM grammar. General PTM site prediction teaches the model residue-level modification compatibility; kinase-specific phosphorylation prediction introduces enzyme-recognition specificity; PTM crosstalk prediction captures conditional dependencies between modification events; and enzyme kinetic regression links local modification syntax to protein-level biochemical function. Thus, ProtSyntax was not trained as a collection of independent predictors, but as a shared sequence–structure model exposed to residue identity, motif order, three-dimensional microenvironment, PTM co-occurrence and functional consequence within a unified representation space.

For general PTM site prediction, experimentally supported residue-level PTM annotations were collected from public protein resources and the literature^34^. Records were mapped to reference protein sequences, standardized by modification type and residue coordinate, and deduplicated at the sequence–site level^35^. We removed annotations with ambiguous modification types, non-unique residue localization, evidence inferred only from homology transfer or text mining, or sequences that could not be reliably matched to a reference protein^36^. The resulting PTM-language corpus contained 348,903 positive site-centered samples and 3,902,925 chemically compatible negative site-centered samples across 40 PTM classes. Each sample was represented as a 55-residue window centered on the candidate residue. Positive samples corresponded to experimentally validated PTM sites, whereas negative samples were sampled from residues chemically compatible with the target PTM type but lacking an annotation for that modification. To reduce homology-driven leakage, protein sequences were clustered with CD-HIT at 50% sequence identity before data splitting, and all site-centered windows from proteins in the same cluster were assigned to the same partition. Training, validation and test sets were split at a ratio of 8:1:1 using random seed 42.

For kinase-specific phosphorylation, kinase–substrate annotations were integrated from the DCPPS^24^ study and complementary public resources, followed by redundancy reduction and residue-centered window extraction. The final dataset contained four kinase-specific phosphorylation classes with 3,803 positive and 41,889 negative samples. For PTM crosstalk prediction, we built on the DeepPCT^25^ dataset and incorporated additional curated crosstalk annotations, resulting in 262 positive and 12,468 negative samples. For protein functional supervision, we used CatPred-DB^26^, comprising 77,020 experimentally measured enzyme kinetic records, including 23,917 K_cat_, 41,174 K_m_ and 11,929 K_i_ samples.

Owing to space limitations in the main text, the detailed composition of the dataset is provided in Supplementary Section 2 and Supplementary Table S1, which summarizes the annotation statistics and characteristics of all 40 PTM types included in this study. In addition, we performed a comprehensive visual analysis of the PTM dataset to further characterize its distributional properties, residue preferences, functional enrichment patterns and modification co-occurrence relationships, as shown in Figure S1.

Because PTM annotation and enzyme kinetics operate at different scales, ProtSyntax uses 55-residue-centered windows for PTM training and full-length sequences for kinetic regression. The window defines the training unit rather than the inference limit: proteins up to 1,024 amino acids can be profiled using overlapping sliding windows, with predictions mapped to native residue coordinates. All tasks share the same tokenizer and backbone, use task identifiers and task-homogeneous batches, and integrate overlength sequences through pooled window representations.

### 4.2 Protein representations and PTM-aware positional encoding

#### 4.2.1 Bio-RoPE positional encoding

PTM syntax depends on more than token order. A candidate modification site is shaped by the relative arrangement of flanking residues, secondary-structure periodicity, residue chemistry and long-range protein context. Standard rotary positional encoding captures relative sequence order but does not distinguish whether a position is occupied by a residue with compatible physicochemical properties for a given modification^37^. We therefore developed Bio-RoPE, a PTM-aware rotary positional encoding that decomposes positional phase into structural-periodic, physicochemical and standard long-range channels. This design allows ProtSyntax to encode not only where a residue appears in a protein window, but also whether its positional context is chemically and structurally compatible with PTM grammar.

Given a residue representation *x_i_* at position *i*, Bio-RoPE partitions the hidden channels into three groups: structural-periodic channels, physicochemical phase-modulation channels and standard long-range rotary channels. The structural-periodic channels encode canonical protein periodicities using the period set = {2.0,3.0,3.6,5.1}, corresponding to extended strand-like periodicity, 3_10_-helices periodicity, α-helical periodicity and π-helical periodicity. For a period *p_j_*, the structural phase is defined as:

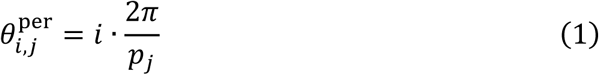

The physicochemical channels introduce amino-acid-dependent phase offsets. For residue identity*a_i_*, ProtSyntax retrieves a frozen descriptor vector *r*(*a_i_*), encoding hydrophobicity, helical propensity and side-chain steric bulk. This descriptor is projected into phase offsets and added to the base rotary phase:

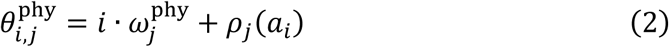

The remaining channels retain the standard RoPE formulation, 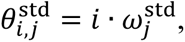 to preserve long-range relative positional modeling. The final phase vector is obtained by concatenating the three channel groups and applying rotary transformation to query and key representations. In this way, residues at the same sequence position can receive different phase modulations according to amino acid chemistry, allowing ProtSyntax to encode PTM-compatible residue syntax together with local structural periodicity. The detailed mathematical formulation of Bio-RoPE is provided in Supplementary Section 3.1.

#### 4.2.2 Sequence and structure representation

ProtSyntax integrates protein language representations with explicit residue-level geometry. For each protein sequence, contextual residue embeddings were generated offline from ESM-C and SaProt using their official pretrained checkpoints. These pretrained encoders were used strictly as frozen feature extractors; their parameters were not updated during ProtSyntax training, and no gradients were propagated back into ESM-C^38^ and SaProt^39^. ESM-C provides sequence-derived semantic representations learned from large-scale protein sequences, whereas SaProt contributes structure-aware residue representations. The resulting fixed embeddings were projected into a shared hidden space through trainable projection layers and combined with amino acid token embeddings before being passed into the ProtSyntax encoder.

For structural modeling, AlphaFold2-predicted PDB structures were used to derive residue-level geometric features^40^. For each residue, we computed Cα coordinates, backbone-based local rigid frames and pairwise spatial relationships. These features were not treated as independent descriptors appended after sequence encoding; instead, they were used by Geometric Gated Attention to constrain residue–residue interactions within the encoder. Residues with missing coordinates or unreliable structural confidence were masked in the geometric branch, allowing the corresponding attention heads to revert to sequence-based semantic attention. This design enables ProtSyntax to use three-dimensional information when reliable structures are available while remaining applicable to low-confidence structures, intrinsically disordered regions and sequence-only inputs.

### 4.3 ProtSyntax architecture

ProtSyntax is a PTM-centered protein large language model designed to decode post-translational modification syntax and its functional consequences. Unlike conventional PTM predictors that treat each candidate residue as an independent classification target, ProtSyntax formulates PTM regulation as a structured protein language problem. In this framework, PTM syntax refers to the hierarchical rules by which residue chemistry, local motif order, long-range sequence context, three-dimensional accessibility and existing modification states jointly determine whether a residue can be modified, which PTM type is contextually compatible, how different PTMs condition one another, and how local modification events are translated into protein-level functional outcomes. ProtSyntax implements this syntax-to-function formulation through a shared sequence– structure encoder coupled with task-specific heads for general PTM-site prediction, kinase-specific phosphorylation, PTM crosstalk modeling and enzyme kinetic regression.

The architecture of ProtSyntax is therefore organized around the biological requirements of PTM grammar rather than inherited unchanged from general-purpose protein language models. ProtSyntax is trained from scratch as a PTM-centered large protein model and uses a sparse Mixture-of-Experts backbone to expand representational capacity while maintaining computational efficiency. Within this backbone, Bi-Gated DeltaNet serves as the long-range syntax propagation module: it scans protein sequences bidirectionally from N-to-C and C-to-N directions and cross-gates the two directional states, allowing the model to reinforce residues supported by coherent upstream and downstream evidence while suppressing locally plausible but globally incidental motif signals. Geometric Gated Attention serves as the structural grammar module: by injecting residue-frame-based three-dimensional constraints directly into attention logits, it enables ProtSyntax to distinguish residues that are merely sequence-compatible from those embedded in genuinely permissive PTM microenvironments. By default, ProtSyntax interleaves Bi-Gated DeltaNet and Geometric Gated Attention blocks at a 3:1 ratio^41^, allowing most layers to efficiently propagate long-range PTM syntax while selected layers perform structure-constrained residue interaction reasoning. Together with PTM-aware positional encoding and multi-objective syntax–function supervision, this design enables ProtSyntax to move beyond site annotation toward rare-PTM generalization, combinatorial crosstalk reconstruction and quantitative inference of PTM-driven functional effects. The overall architecture of ProtSyntax is illustrated in Figure 6.

**Figure 6.**
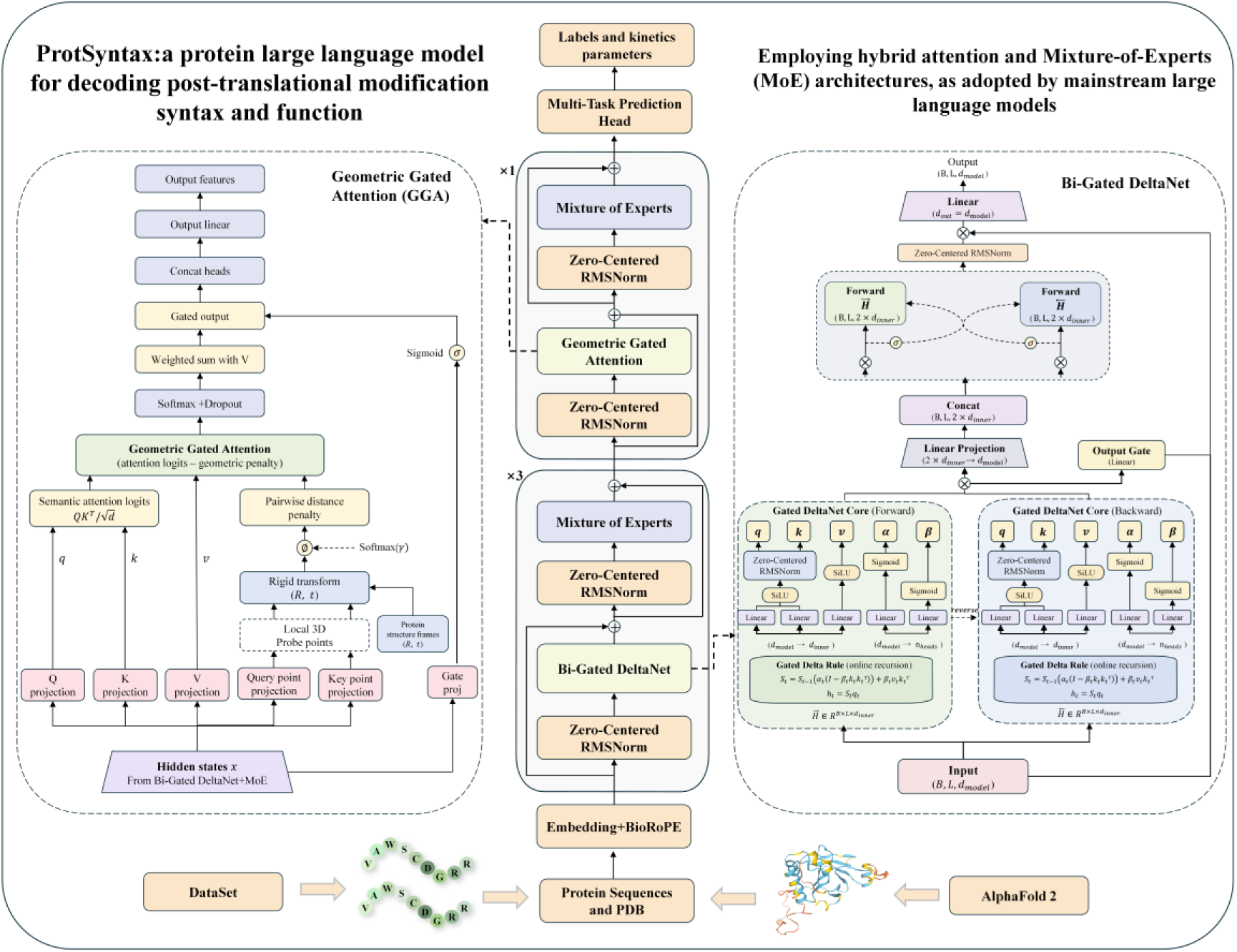
Architecture of ProtSyntax. ProtSyntax is built upon a modern hybrid attention and mixture-of-experts architecture. Within this architecture, the Bi-Gated DeltaNet is implemented as a linear attention module aimed at breaking the quadratic computational bottleneck of standard attention. Meanwhile, the Geometric Gated Attention functions as a global attention mechanism; by retaining the standard Transformer’s capability to precisely align any token with all historical tokens in a sequence, it ensures lossless global information retrieval.

It is important to note that the detailed mathematical derivations of Bio-RoPE, Bi-Gated DeltaNet, Geometric Gated Attention and the PACE-Nash loss are provided in Supplementary Section 3. In the main text, we focus on the architectural rationale, computational workflow and biological interpretation of these modules, whereas the supplementary material provides the complete recurrent update equations, geometric attention formulation and multi-objective optimization details required to ensure technical transparency and reproducibility.

#### 4.3.1 Bi-Gated DeltaNet for bidirectional PTM syntax propagation

Bi-Gated DeltaNet was designed to propagate PTM-relevant sequence evidence along both directions of a protein chain with linear sequence complexity. PTM occurrence is often determined by a candidate residue together with upstream and downstream motifs, local physicochemical context, domain boundaries and distal sequence cues. A single-direction recurrent module can efficiently summarize sequence history, but it may overemphasize one-sided incidental motifs and fail to evaluate whether N-terminal and C-terminal contexts support the same modification event. Bi-Gated DeltaNet addresses this limitation by using two parameter-independent Gated DeltaNet cores that scan the sequence in opposite directions^41^, followed by a zero-parameter cross-gating operation that conditionally recalibrates the two directional states at each residue. This makes the module function as a bidirectional syntax filter: residues are strengthened when both sides provide coherent PTM evidence and weakened when a motif is locally plausible but globally unsupported.

For an input representation X, each Gated DeltaNet core projects the input into multi-head query, key and value representations:

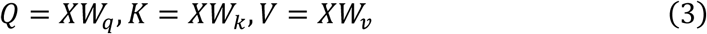

The query and key vectors are normalized and passed through a SiLU activation before recurrent state writing. At residue position *t* and head *h*, the core computes a decay gate *α_t_*_,ℎ_ and a write gate *β_t_*_,ℎ_, which control memory retention and incorporation of the current residue. The head-wise recurrent state *S_t_*_,ℎ_ is updated using a gated Delta rule that selectively decays outdated memory and writes current residue information through a rank-one update. The output at each position is retrieved by querying the recurrent state with the normalized query vector.

Bi-Gated DeltaNet applies this recurrence in both sequence directions. The forward core summarizes N-terminal-to-C-terminal context, whereas the backward core summarizes C-terminal-to-N-terminal context and is then realigned to the original residue coordinates. The two directional representations are fused by cross-gating:

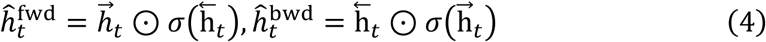

This operation allows downstream context to gate upstream evidence and upstream context to gate downstream evidence without adding learnable parameters. The cross-gated states are concatenated, projected back to the model hidden dimension and filtered by an input-conditioned output gate. In PTM prediction, this mechanism reduces false-positive signals arising from one-sided incidental motifs and strengthens residues supported by coherent bidirectional context.

#### 4.3.2 Geometric Gated Attention for structure-constrained PTM microenvironment syntax

Geometric Gated Attention (GGA) was introduced to represent PTM microenvironment syntax that cannot be resolved from linear sequence alone. Many modification events depend on spatially proximal residues, backbone conformation, solvent exposure and domain organization. A residue may be chemically compatible with a PTM in sequence space but inaccessible or incorrectly oriented in the folded protein^42^. Therefore, instead of concatenating structural descriptors to sequence embeddings, GGA injects geometry directly into the attention logits. This converts attention from a purely semantic residue interaction mechanism into a structure-constrained interaction mechanism, enabling ProtSyntax to identify PTM sites supported by both sequence grammar and three-dimensional permissiveness.

Given input representation *X*, GGA first computes standard multi-head query, key and value projections. For each residue *i*, attention head hhh and geometric probe mmm, the module predicts query-side and key-side probe points in local residue coordinates. These points are transformed into global coordinates using residue-wise backbone rigid frames (*R_i_*, *t_i_*):

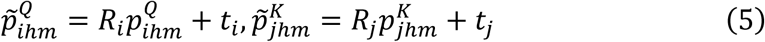

The squared Euclidean distance between query-side and key-side probes is then used as a head-specific geometric penalty on the semantic attention score:

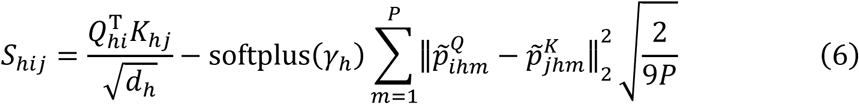

The learned scaling parameter *γ*_ℎ_ controls the strength of geometric correction for each head. This formulation suppresses residue pairs that are semantically compatible but geometrically inconsistent, while preserving high attention for residue interactions supported by both sequence context and three-dimensional proximity. After attention aggregation, a query-dependent sigmoid gate regulates the contribution of each head, filtering redundant structural–semantic responses that are not informative for PTM discrimination. To stabilize optimization, the geometric point projections were initialized to zero, making GGA behave like standard semantic attention at the beginning of training and allowing geometric constraints to emerge progressively under PTM supervision.

#### 4.3.3 PACE-Nash loss for coupling PTM syntax with protein function

ProtSyntax was trained with Physicochemical-Aware Contrastive and Evidential Nash-Bargaining loss, abbreviated PACE-Nash, to align residue-level PTM syntax with protein-level functional supervision. PTM-centered protein learning faces three coupled challenges: modified residues are sparse relative to chemically compatible unmodified residues; kinetic measurements contain heteroscedastic experimental uncertainty; and residue-level classification objectives can conflict with full-length protein regression objectives. PACE-Nash addresses these challenges by combining asymmetric focal classification, correlation-aware contrastive learning, uncertainty-aware kinetic regression, physicochemical manifold regularization and Nash bargaining-based adaptive task weighting^43^.

For PTM prediction, we combined an asymmetric focal classification loss with a correlation-aware contrastive objective. The focal term reduces the dominance of abundant negative residues, whereas the contrastive term encourages proteins or residue windows with related PTM patterns to occupy neighboring regions in the latent space^44^. For enzyme kinetic regression, ProtSyntax predicts both the mean and uncertainty of log-transformed Kcat, Km and Ki values, allowing uncertain kinetic measurements to contribute proportionally during training. A physicochemical manifold regularization term further constrains representation distances to remain consistent with distances computed from protein physicochemical descriptors. The resulting task objectives can be summarized as:

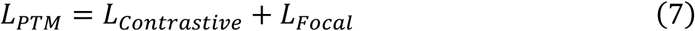

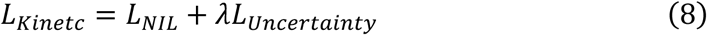

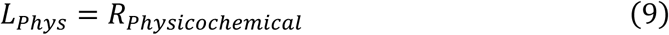

Rather than manually specifying task weights, PACE-Nash formulates multi-objective optimization as a Nash bargaining problem. Gradients generated by the PTM objective, kinetic objective, and physicochemical regularization objective are projected into a shared parameter space, and adaptive bargaining coefficients are obtained by maximizing the collective utility of all objectives. The final optimization target is therefore expressed as:

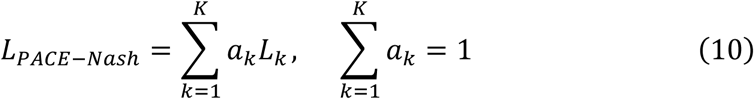

where *a_k_* denotes the dynamically optimized bargaining coefficient for task *k*. This formulation enables adaptive coordination between PTM classification, enzyme kinetic regression, and physicochemical regularization, yielding a Pareto-efficient training process and improving the biological fidelity and generalization capability of the learned protein representations.

### 4.4 Hyperparameters and Model Evaluation Metrics

For model evaluation, we used Matthews correlation coefficient (MCC), area under the receiver operating characteristic curve (AUC) and average precision (AP) for classification tasks, and the coefficient of determination (R^2^) and mean absolute error (MAE) for regression tasks. ProtSyntax was trained on a cluster of 24 NVIDIA H100 GPUs and can be deployed for inference on a H100/A100/5090/4090 GPU server.

In ProtSyntax, each sparse MoE layer comprises 16 experts and uses Top-2 routing, such that only a small subset of experts is activated for each token. The model contains approximately 4 billion (4B) trainable parameters, with approximately 1.0 billion (A1B) active parameters per token, thereby balancing large model capacity with efficient sparse computation. In brief, ProtSyntax used a 33-layer hybrid backbone with a 3:1 interleaving ratio between Bi-Gated DeltaNet and Geometric Gated Attention blocks and a hidden dimension of 1,280. Key settings included 20% structural-periodic and 30% physicochemical channels in Bio-RoPE, 16 heads in Bi-Gated DeltaNet, four geometric probes in Geometric Gated Attention and a contrastive temperature of 0.7 in PACE-Nash optimization. The model was optimized with AdamW using a peak learning rate of 4e-4, 5,000 warm-up steps and a task-dependent batch size of 256 or 512. Detailed information regarding the hyperparameters can be found in Section 5 of the Supplementary Materials. To facilitate broad accessibility, we developed ProtSyntax Lab, a web-based platform that enables biologists and bioinformaticians to perform PTM-related prediction tasks through an intuitive online interface.

For all experiments requiring fine-tuning or retraining of comparison models, we used a harmonized training protocol to ensure a fair evaluation against ProtSyntax. All models were trained on the same data partitions with identical sample definitions, preprocessing procedures, training epochs, early-stopping criteria and checkpoint-selection rules. Whenever architecturally compatible, we also matched the optimization strategy, learning-rate schedule, batch construction and random seeds. Model-specific settings were retained only when required by the original architecture or officially recommended implementation. Hyperparameter selection was performed exclusively on the validation set, and the test set remained untouched until final evaluation. Thus, the reported differences reflect model capability rather than unequal training budgets, data access or optimization conditions.

## Supporting information

Supplementary for main text

## Declarations

## Ethical approval

This article does not contain any studies with animals performed by any of the authors.

## Author Contributions

Yiyu Lin: Methodology, Writing – Original Draft. Jiahui Wu: Software, Validation. You Zhou: Data Curation, Investigation. Xinye Ni: Formal analysis, Visualization. Shan Chang: Resources. Jun Ding: Conceptualization. Yan Wang: Resources. Xin Gao: Methodology. Sen Yang: Methodology and Investigation.

## Data Availability

The dataset associated with this work is publicly available on Hugging Face at: https://huggingface.co/datasets/Ethan-Lin/PTMBenchmark-ProtSyntax.

## Code Availability

The source code for ProtSyntax and ProtSyntax Lab has been made publicly available on GitHub at: https://github.com/Yuqiu-rgb/ProtSyntax, and the model weights have been released on Hugging Face. Additionally, the GitHub repository provides comprehensive documentation and detailed instructions on how to use the model.

## Acknowledgements

The authors express their gratitude to those who provided support for this research. We further recognize the Alibaba Qwen Team for their pioneering efforts in advancing open-source foundation models. Specifically, the implementation of ProtSyntax adapts the optimized engineering infrastructure from the Qwen3-Next family.

## Reference

1. Zhang, D. et al. Metabolic regulation of gene expression by histone lactylation. Nature 574, 575–580 (2019).

2. Li, Y. et al. Pan-cancer proteogenomics connects oncogenic drivers to functional states. Cell 186, 3921–3944.e25 (2023).

3. Gadd, M. S. et al. Structural basis of PROTAC cooperative recognition for selective protein degradation. Nat Chem Biol 13, 514–521 (2017).

4. Decrypting the functional design of unmodified translation elongation factor P. Cell Reports 43, 114063 (2024).

5. Mijit, M., Caracciolo, V., Melillo, A., Amicarelli, F. & Giordano, A. Role of p53 in the Regulation of Cellular Senescence. Biomolecules 10, 420 (2020).

6. Russo, A. A., Jeffrey, P. D. & Pavletich, N. P. Structural basis of cyclin-dependent kinase activation by phosphorylation. Nat Struct Mol Biol 3, 696–700 (1996).

7. Wang, J. et al. Lactylation of PKM2 Suppresses Inflammatory Metabolic Adaptation in Pro-inflammatory Macrophages. Int J Biol Sci 18, 6210–6225 (2022).

8. Bludau, I. et al. The structural context of posttranslational modifications at a proteome-wide scale. PLOS Biology 20, e3001636 (2022).

9. Lin, H. et al. Understanding the immunosuppressive microenvironment of glioma: mechanistic insights and clinical perspectives. J Hematol Oncol 17, 31 (2024).

10. Chen, L. & Chen, Y. RMTLysPTM: recognizing multiple types of lysine PTM sites by deep analysis on sequences. Brief Bioinform 25, bbad450 (2024).

11. Kim, D. N. et al. Artificial Intelligence Transforming Post-Translational Modification Research. Bioengineering 12, 26 (2025).

12. Gerken, J. E. et al. Geometric deep learning and equivariant neural networks. Artif Intell Rev 56, 14605–14662 (2023).

13. Kundu, B. K. & Tanti, B. Decoding plant physiology through systems biology: Integrative multi-omics and computational perspectives for next-generation crop design. Plant Communications 7, 101668 (2026).

14. Li, Y.-Y. et al. A Systematic Review of Computational Methods for Protein Post-Translational Modification Site Prediction. Arch Computat Methods Eng 33, 4287–4307 (2026).

15. Shrestha, P., Kandel, J., Tayara, H. & Chong, K. T. Post-translational modification prediction via prompt-based fine-tuning of a GPT-2 model. Nat Commun 15, 6699 (2024).

16. Han, Y., He, F., Shao, Q., Wang, D. & Xu, D. MTPrompt-PTM: A Multi-Task Method for Post-Translational Modification Prediction Using Prompt Tuning on a Structure-Aware Protein Language Model. Biomolecules 15, 843 (2025).

17. AstraPTM2: A Context-Aware Transformer for Broad-Spectrum PTM Prediction.

18. Tan C. et al. MeToken: Uniform Micro-environment Token Boosts Post-Translational Modification Prediction. International Conference on Learning Representations 2025, 74895–74924 (2025).

19. Koh, H. Y. et al. AI-driven protein design. Nat Rev Bioeng 3, 1034–1056 (2025).

20. Beltrao, P., Bork, P., Krogan, N. J. & van Noort, V. Evolution and functional cross-talk of protein post-translational modifications. Mol Syst Biol 9, MSB134521 (2013).

21. Rives, A. et al. Biological structure and function emerge from scaling unsupervised learning to 250 million protein sequences. Proceedings of the National Academy of Sciences 118, e2016239118 (2021).

22. Venne, A. S., Kollipara, L. & Zahedi, R. P. The next level of complexity: Crosstalk of posttranslational modifications. PROTEOMICS 14, 513–524 (2014).

23. Catalytic activity regulation through post-translational modification: the expanding universe of protein diversity. in Advances in Protein Chemistry and Structural Biology vol. 122 97–125 (Academic Press, 2020).

24. Liu, M., Wang, X., Sun, Z.-L., Yang, X. & Chen, X. DCPPS: Prediction of Kinase-Specific Phosphorylation Sites Using Dynamic Embedding and Cross-Representation Interaction. Interdiscip Sci Comput Life Sci 17, 1056–1073 (2025).

25. Huang, Y.-X. & Liu, R. Improved prediction of post-translational modification crosstalk within proteins using DeepPCT. Bioinformatics 40, btae675 (2024).

26. Boorla, V. S. & Maranas, C. D. CatPred: a comprehensive framework for deep learning in vitro enzyme kinetic parameters. Nat Commun 16, 2072 (2025).

27. Krassowski, M. et al. ActiveDriverDB: human disease mutations and genome variation in post-translational modification sites of proteins. Nucleic Acids Res 46, D901–D910 (2018).

28. Owen, I. & Shewmaker, F. The Role of Post-Translational Modifications in the Phase Transitions of Intrinsically Disordered Proteins. International Journal of Molecular Sciences 20, 5501 (2019).

29. Salih A. M. et al. A Perspective on Explainable Artificial Intelligence Methods: SHAP and LIME. doi:10.1002/aisy.202400304.

30. Cui, X. et al. Beyond static structures: protein dynamic conformations modeling in the post-AlphaFold era. Brief Bioinform 26, bbaf340 (2025).

31. Franciosa, G., Locard-Paulet, M., Jensen, L. J. & Olsen, J. V. Recent advances in kinase signaling network profiling by mass spectrometry. Curr Opin Chem Biol 73, 102260 (2023).

32. Smith, L. M. & Kelleher, N. L. Proteoform: a single term describing protein complexity. Nat Methods 10, 186–187 (2013).

33. Victoriano, M. et al. From virtual experiments to biomedical insight with synthetic data. Nat Mach Intell 8, 866–879 (2026).

34. The UniProt Consortium. UniProt: a worldwide hub of protein knowledge. Nucleic Acids Res 47, D506–D515 (2019).

35. Li, Z. et al. dbPTM in 2022: an updated database for exploring regulatory networks and functional associations of protein post-translational modifications. Nucleic Acids Res 50, D471–D479 (2022).

36. Hornbeck, P. V. et al. PhosphoSitePlus: a comprehensive resource for investigating the structure and function of experimentally determined post-translational modifications in man and mouse. Nucleic Acids Res 40, D261–D270 (2012).

37. Lazos, L., Poovendran, R. & Capkun, S. Rope: robust position estimation in wireless sensor networks. in *IPSN 2005*. Fourth International Symposium on Information Processing in Sensor Networks, 2005. 324–331 (IEEE, Los Angeles, CA, USA, 2005). doi:10.1109/IPSN.2005.1440942.

38. Hayes, T. et al. Simulating 500 million years of evolution with a language model. Science 387, 850–858 (2025).

39. Su, J. et al. SaProt: Protein Language Modeling with Structure-aware Vocabulary. International Conference on Learning Representations 2024, 6987–7009 (2024).

40. Yang, Z., Zeng, X., Zhao, Y. & Chen, R. AlphaFold2 and its applications in the fields of biology and medicine. Sig Transduct Target Ther 8, 115 (2023).

41. Yang, A., et al. Qwen3 Technical Report. Preprint at 10.48550/arXiv.2505.09388 (2025).

42. Qiu, Z. et al. Gated Attention for Large Language Models: Non-linearity, Sparsity, and Attention-Sink-Free. in Advances in Neural Information Processing Systems vol. 38 100092–100118 (Curran Associates, Inc., 2025).

43. The Bargaining Problem - The Econometric Society. http://www.econometricsociety.org/publications/econometrica/1950/04/01/bargaining-problem.

44. Wen, B. et al. DeepMVP: deep learning models trained on high-quality data accurately predict PTM sites and variant-induced alterations. Nat Methods 22, 1857–1867 (2025).

