## Supplementary for main text for "ProtSyntax: a protein large language model for decoding post-translational modification syntax and function"

### Supplementary Materials

Given the length and breadth of the Supplementary Information, we provide a brief overview of the purpose of each section to facilitate navigation and improve readability:

**1.Related work and methodological positioning of ProtSyntax.** This section extends the discussion of prior studies presented in the main text. It reviews the evolution of model architectures in the PTM prediction field and provides a detailed analysis of the major methodological challenges and unresolved limitations.

**2.Dataset Description.** This section provides additional details on the datasets used in this study, including their sources, construction, preprocessing and partitioning procedures.

**3.Methods.** This section expands upon the Methods described in the main text and provides additional technical details regarding the model architecture, individual components, training objectives and implementation procedures.

**4.Results.** This section comprehensively supplements the Results presented in the main text. In addition to reporting detailed results for the primary experiments, it includes further validation experiments and more in-depth analyses.

**5.Hyperparameters.** This section lists the detailed hyperparameter settings used for model training and inference.

**6.Detailed performance metrics of the comparative models.** This section reports the complete quantitative results of the comparative evaluations between ProtSyntax and existing models. Because it contains many data-intensive tables, it is placed near the end of the Supplementary Information to preserve the continuity and readability of the preceding sections.

**7.Extended Discussion.** This section provides a more detailed discussion of the methodological implications, limitations and broader significance of this work.

#### 1.Related work and methodological positioning of ProtSyntax

Early computational studies of post-translational modification focused primarily on phosphorylation, for which relatively abundant experimental annotations provided a tractable setting for developing site-prediction algorithms. These methods generally treated PTM recognition as a supervised pattern-classification problem in which the biochemical status of a candidate residue was inferred from its surrounding sequence and a predefined set of descriptors. NetPhos[1] represented an important departure from rigid consensus-motif matching by using artificial neural networks to learn nonlinear sequence patterns surrounding serine, threonine and tyrosine residues. This work established that modification competence could be captured statistically rather than encoded exclusively through manually specified motifs. GPS [2]subsequently introduced a hierarchical formulation of kinase-specific phosphorylation prediction, organizing kinases into groups, families, subfamilies and individual enzymes and using amino acid substitution similarity to characterize substrate preferences at different levels of specificity. Its major contribution was to connect candidate phosphosites to the recognition logic of particular kinase classes, rather than treating phosphorylation as a single homogeneous event. Musite further combined local amino acid composition, similarity to experimentally annotated phosphosites and intrinsic-disorder propensity within an ensemble-learning framework, enabling both general and kinase-specific prediction across multiple organisms and at proteome scale. Collectively, these methods established the basic computational architecture of PTM-site prediction: experimentally annotated sites were converted into residue-centred examples, biologically motivated features were extracted, and a classifier was trained to distinguish modified from unmodified residues. Their success demonstrated that PTM occurrence is computationally learnable, but the scope of the inferred biology remained largely constrained by the descriptors, similarity functions and prediction targets specified in advance.

The emergence of deep learning shifted the field from manually engineered features toward end-to-end representation learning. MusiteDeep[3] directly processed amino acid sequences with convolutional neural networks, allowing local PTM-recognition patterns to be learned from data rather than prescribed through handcrafted descriptors. Its attention mechanisms emphasized informative sequence positions and feature channels, while transfer from general phosphorylation data to kinase-specific tasks showed that sequence representations learned from abundant annotations could be reused when kinase-specific labels were limited. This was a substantive conceptual advance because it recast PTM prediction as representation learning rather than feature selection. PDeepPP[4] extended this direction by combining pretrained ESM-2 representations with task-adaptive residue embeddings and parallel convolutional and Transformer branches. The convolutional component captured short-range sequence regularities, whereas the Transformer component modeled broader contextual dependencies. Its information-maximization objective further addressed the unstable decision boundaries induced by severe class imbalance, illustrating how pretrained protein representations and task-specific optimization could be integrated within a general peptide-classification framework. MTPrompt-PTM[5] advanced multi-PTM modeling by applying prompt-based adaptation to a structure-aware protein language model. A shared encoder supported several PTM tasks, PTM-specific heads preserved modification-dependent decision boundaries, and knowledge distilled from single-task teacher models transferred task-specific information into a unified student model. The main contribution of this generation of methods was therefore not only improved predictive accuracy, but also a shift from isolated, manually designed classifiers toward reusable representations that could share information across modification types and benefit from large-scale protein pretraining.

More recent approaches have incorporated generative language modeling, state-space architectures and full-length protein representations, reflecting a broader transition toward PTM-specialized foundation models. PTMGPT2 [6]reformulated site prediction as prompt-conditioned label generation, using a GPT-2-derived protein model to generate modification labels from protein fragments and task prompts. This formulation demonstrated that PTM prediction could be embedded within a generative language-modeling objective, while decoder attention was used to inspect candidate motifs and mutation-sensitive sequence regions. PTM-Mamba[7] introduced a different representation principle by treating modified residues as explicit vocabulary elements rather than only as output labels. Bidirectional Mamba blocks encoded wild-type and modified protein sequences, and PTM-focused masked modeling encouraged the network to recover modification-specific tokens. By representing modified proteoforms directly, the model broadened PTM learning beyond conventional site classification to disease association, druggability, interaction modeling and zero-shot analyses. AstraPTM2[8] pursued broad-spectrum prediction across full-length proteins by combining pretrained sequence representations, AlphaFold2-derived structural information and protein-level descriptors within a context-aware Transformer. Multi-scale processing, curriculum learning, imbalance-aware optimization and label-specific calibration enabled the simultaneous prediction of dozens of PTM types. Together, these models substantially expanded PTM coverage, increased the amount of sequence and structural context available to the predictor, and demonstrated the value of PTM-oriented pretraining and multi-task inference. They mark an important progression from independent site classifiers toward models that represent broader aspects of the modified proteome.

Despite this progression, most computational PTM models remain organized around residue-level label assignment rather than around the regulatory logic that determines why a modification is permissible, selective and functionally consequential. The limitations of the earlier methods follow directly from their methodological formulations. NetPhos and GPS 2.0 learned phosphorylation propensity from local sequence patterns, kinase hierarchies and motif similarity, whereas Musite supplemented such signals with disorder and neighbourhood-based descriptors; these approaches were effective for motif-driven recognition but could only represent biochemical relationships encoded in the predefined feature space. Deep-learning models reduced this dependence on manual descriptors, yet their objectives remained similarly site-centric. MusiteDeep used convolutional filters and attention to learn discriminative local patterns, but its transfer from general to kinase-specific phosphorylation primarily reused motif-level representations rather than a unified biochemical organization of PTM classes. PDeepPP combined ESM-2 embeddings with convolutional and Transformer branches and improved robustness to class imbalance, but its latent space was optimized for classification separation rather than for preserving relationships among residue chemistry, modification mechanisms and functional compatibility. MTPrompt-PTM enabled parameter sharing through prompts, PTM-specific heads and teacher–student distillation; however, knowledge transfer was mediated by shared parameters and teacher predictions, not by explicit constraints that chemically related PTMs should share mechanistic rules while unrelated modifications should remain distinct. Similarly, PTMGPT2, PTM-Mamba and AstraPTM2 expanded PTM coverage and representation capacity, but their principal supervision still emphasized the recovery or assignment of modification labels. Consequently, cross-PTM sharing may reflect statistical co-occurrence, dataset frequency or representation similarity rather than genuinely transferable biochemical syntax, which limits reliable generalization to rare, weakly annotated or previously unseen modification types.

A related methodological gap lies in the distinction between broad contextual encoding and mechanistic permissiveness. PTMGPT2 uses prompt-conditioned generative prediction over protein fragments, allowing contextual sequence information to influence label generation, but decoder attention does not explicitly test whether distal evidence from both sides of a candidate residue coherently supports the same regulatory event, nor does it impose physical constraints on residue interactions. MusiteDeep and PDeepPP extend the effective receptive field through convolutional and Transformer components, yet their context aggregation remains primarily statistical: a strong local motif can still dominate even when it is inconsistent with domain organization, distal regulatory elements or enzyme-recognition context. PTM-Mamba improves long-range efficiency through bidirectional state-space propagation and introduces explicit tokens for modified residues, but its central representation is optimized for sequence-state modeling and PTM-token recovery rather than for determining whether a wild-type candidate site occupies a physically admissible modification microenvironment. MTPrompt-PTM incorporates structure-aware pretrained representations, and AstraPTM2 further combines full-length sequence embeddings with AlphaFold2-derived structural features and protein-level descriptors; nevertheless, in both cases structural information is principally supplied as an enriched representation or auxiliary feature source. Such integration can capture correlations with secondary structure, solvent exposure or global fold context, but it does not necessarily require residue–residue interactions to obey explicit three-dimensional geometry. This distinction is biologically important because sequence-distant residues may converge to form a catalytic, binding or regulatory microenvironment, whereas sequence-adjacent residues may be sterically occluded or inaccessible to the modifying enzyme. A mechanistically grounded PTM model must therefore do more than associate a candidate site with broad sequence or structural context: it must jointly evaluate local chemical compatibility, coherent bidirectional evidence and the explicit three-dimensional permissiveness of the surrounding residue environment.

A third limitation is that broad-spectrum and multi-task prediction do not by themselves recover the conditional organization of the modified proteome. Most existing frameworks produce several PTM labels from a shared encoder but still optimize each label primarily as an independent prediction target. Consequently, the presence of multiple predicted modifications does not reveal whether one event promotes, suppresses or alters the functional interpretation of another. Cooperative, antagonistic and order-dependent PTM relationships are fundamental to signalling, degradation, chromatin regulation and phase separation, yet they cannot be reliably inferred from co-occurrence alone. A shared regulatory representation must encode how the probability and meaning of one modification change when another site is added, removed or perturbed.

Most importantly, the field has largely separated PTM occurrence from PTM function. A high-confidence site prediction indicates that a residue resembles previously observed modification sites, but it does not establish whether the event changes catalytic turnover, substrate affinity, protein stability, interaction specificity, conformational communication or disease-associated behaviour. Site prediction and functional-effect modeling are therefore commonly treated as distinct problems, despite the fact that PTMs are biologically important precisely because local chemical changes propagate into protein-level consequences. Without shared supervision linking residue-level events to quantitative functional measurements, a model may become increasingly accurate at annotation while remaining unable to distinguish functionally consequential modifications from biochemically tolerated ones. Thus, residue chemistry, positional organization, long-range contextual consistency, three-dimensional permissiveness, PTM crosstalk and functional consequence remain only partially connected in current computational frameworks.

These limitations motivate a more fundamental formulation of PTM modeling. PTMs should not be treated merely as labels appended to protein sequences, but as elements of a regulatory language through which proteins encode context-dependent functional states. In this language, amino acid identity and physicochemical properties define the available vocabulary; residue order and motif organization establish local syntactic constraints; protein-scale sequence context and three-dimensional geometry determine whether that syntax is valid in a specific molecular setting; interactions among modification events provide conditional grammar; and the resulting effects on protein activity supply functional semantics. Learning such a language requires more than improving classification performance on common PTM benchmarks. It requires a model that can extract rules shared across chemically related modifications, distinguish locally plausible motifs from contextually and geometrically permissible sites, represent dependencies among multiple PTMs, and connect local modification events to measurable protein-level effects.

This need is amplified by the nature of current PTM data. Experimental annotations are incomplete, heavily imbalanced across modification classes, biased toward well-studied proteins and conditions, and particularly sparse for transient or low-abundance PTMs. Under these conditions, task-specific predictors can exploit recurring patterns in abundant classes but are poorly positioned to recover the underlying regulatory principles required for low-resource generalization. A PTM language model offers a different scientific objective: rather than fitting each modification class as an isolated endpoint, it seeks to infer a shared system of chemical, positional, structural and functional constraints from heterogeneous PTM observations. Such a representation should allow knowledge learned from abundant modifications to be reused for rare PTMs when their underlying chemistries are related, while avoiding transfer between superficially similar but mechanistically incompatible events.

Motivated by this perspective, ProtSyntax was developed as a protein large language model designed specifically to understand PTM language. The central objective is not simply to increase the number of PTM types that can be predicted, but to determine whether diverse residue-level annotations, kinase–substrate relationships, modification crosstalk and enzyme-kinetic measurements can be organized into a coherent representation of PTM regulation. This formulation asks a broader set of questions than conventional site prediction: why is a chemically compatible residue modified at one position but not another; which local and protein-scale contexts make the event permissible; how do neighbouring or distal modifications alter its probability and interpretation; and how can a local modification propagate into a measurable change in protein function? Addressing these questions is necessary for reliable generalization to low-resource PTMs, mechanistic interpretation of combinatorial modification patterns and computational analysis of disease-associated regulatory perturbations. ProtSyntax therefore reframes PTM prediction as the learning of a regulatory protein language in which modification identity, contextual permissiveness, inter-PTM dependency and functional consequence are different levels of the same biological syntax.

ProtSyntax was designed to address the major limitations of current PTM predictors by integrating residue chemistry, sequence context, three-dimensional geometry and protein-level function within a unified PTM-language framework.

1) **Bio-RoPE encodes chemically and structurally informed positional syntax.** Conventional positional encodings primarily represent residue order and assign the same positional phase independently of amino-acid identity, limiting their ability to distinguish chemically meaningful motifs from positionally similar but biologically incompatible patterns. Bio-RoPE addresses this limitation by combining canonical protein structural periodicities, residue-dependent physicochemical phase modulation and standard long-range relative-position channels. This formulation embeds motif order, side-chain chemistry and local structural rhythm directly into positional representations, facilitating the transfer of biochemical rules across related PTMs while reducing reliance on label frequency or superficial sequence similarity.
2) **Bi-Gated DeltaNet resolves bidirectional contextual consistency.** Existing long-context encoders can associate candidate sites with distant sequence regions but do not explicitly determine whether N- and C-terminal evidence coherently supports the same modification event. Bi-Gated DeltaNet performs parallel N-to-C and C-to-N state propagation and uses cross-gating to conditionally recalibrate the two directional representations at each residue. Consequently, globally coherent PTM signals are reinforced, whereas locally plausible but one-sided incidental motifs are suppressed. Its linear sequence complexity further enables efficient propagation of PTM-relevant information across extended protein contexts.
3) **Geometric Gated Attention models explicit three-dimensional permissiveness.** Structural embeddings or concatenated descriptors may provide structural correlations without ensuring that residue interactions obey physical geometry. Geometric Gated Attention instead introduces backbone-frame-based probe points and pairwise geometric penalties directly into the attention logits. It therefore prioritizes residue interactions supported by both semantic compatibility and spatial organization, while suppressing sequence-compatible contacts that are inaccessible or geometrically inconsistent. This design enables ProtSyntax to distinguish motif-like false positives from genuinely permissive PTM microenvironments and to capture spatial coupling between residues that are distant in sequence.
4) **PACE-Nash couples PTM syntax to transferable and uncertainty-aware**

**functional learning.** PTM datasets are strongly imbalanced, rare modifications are sparsely annotated, enzyme-kinetic measurements contain heterogeneous uncertainty, and residue-level classification can conflict with protein-level regression. PACE-Nash integrates asymmetric focal classification, correlation-aware contrastive learning, physicochemical manifold regularization, evidential kinetic regression and Nash bargaining-based adaptive task weighting. These objectives organize related PTM chemistries within a transferable latent space, reduce domination by abundant negatives, account for uncertainty in kinetic supervision and mitigate destructive gradient interference. Rather than allowing PTM annotation and functional prediction to coexist as weakly connected tasks, PACE-Nash coordinates them within a shared representation, thereby linking local modification syntax and inter-PTM dependencies to measurable protein-level consequences.

#### 2. Dataset Description

We selected a CD-HIT sequence-identity threshold of 0.5 as a prespecified compromise between minimizing homology-driven information leakage and preserving sufficient sequence and PTM-class diversity. Because homologous proteins can share conserved acceptor residues, recognition motifs and structural microenvironments, all proteins within the same cluster—and all corresponding residue-centred samples—were assigned to a single data partition. More stringent thresholds, such as 0.3 or 0.4, could unnecessarily collapse remotely related proteins and disproportionately reduce rare-PTM coverage. We did not perform exhaustive threshold comparisons because each setting would require complete reconstruction and rebalancing of the multi-million-sample dataset, regeneration of sequence and structural inputs, and full retraining of the multi-billion-parameter model, ideally across multiple random seeds. Such an analysis would impose a computational cost that is prohibitive for an academic research setting and disproportionate to this methodological choice. We therefore fixed the 0.5 threshold before training and applied it consistently across all dataset partitions and evaluations, avoiding post hoc selection based on test performance.

For PTM site prediction, we adopted a residue-centered window strategy rather than using full-length proteins as direct classification inputs. This design reflects the biological nature of PTM recognition, in which modification propensity is primarily determined by the candidate residue, its flanking sequence motif, local physicochemical environment and nearby structural context. By centering each sample on the candidate residue, the model is explicitly guided to learn PTM-relevant local syntax while reducing interference from distant sequence regions that may be unrelated to the specific modification event. We selected a 55-amino-acid window, corresponding to 27 residues upstream and downstream of the candidate site, as a balanced context size. This window is sufficiently broad to capture enzyme-recognition motifs, short-range residue dependencies, secondary-structure-associated periodic patterns and local microenvironmental cues, while remaining compact enough to avoid excessive background noise, redundant padding and unnecessary computational cost. Therefore, the 55-residue residue-centered formulation provides an effective compromise between biological contextual completeness and modeling efficiency for large-scale PTM site prediction.

Although PTM-site supervision was constructed using 55-residue, site-centered windows, this formulation defines the training sample rather than the inference scope of ProtSyntax. During inference, ProtSyntax supports sequence-wide PTM profiling for proteins of up to 1,024 amino acids through an overlapping sliding-window procedure, generating residue-level prediction scores for candidate sites and mapping them back to their native positions along the protein sequence. This strategy preserves the PTM-relevant local context on which the classifier was trained while enabling systematic screening across substantially longer protein sequences. Thus, ProtSyntax is not restricted to isolated residue-centered fragments; the window-based formulation is primarily used to concentrate supervision on modification-relevant sequence and structural microenvironments, whereas deployment supports whole-protein PTM annotation.

Table S1. Detailed Dataset Distribution

| No | PTM Type | Rarity | Positive | Negative |
|---|---|---|---|---|
| 1 | Phosphorylation | Mainstream | 119995 | 1349347 |
| 2 | Acetylation | Mainstream | 37713 | 424081 |
| 3 | Ubiquitination | Mainstream | 32570 | 366251 |
| 4 | N-linked glycosylation | Mainstream | 22285 | 250593 |
| 5 | O-linked glycosylation | Mainstream | 18856 | 212040 |
| 6 | Methylation | Mainstream | 23999 | 269870 |
| 7 | Sumoylation | Mainstream | 10285 | 115658 |
| 8 | S-palmitoylation | Mainstream | 8571 | 96382 |
| 9 | Disulfide bond | Mainstream | 13714 | 154211 |

| | | | | |
|---|---|---|---|---|
| 10 | S-Sulfhydration | Relatively mainstream | 3500 | 22738 |
| 11 | S-Carboxyethylation | Relatively mainstream | 2561 | 24910 |
| 12 | ADP-ribosylation | Relatively mainstream | 4790 | 53974 |
| 13 | Neddylation | Relatively mainstream | 2743 | 30842 |
| 14 | Succinylation | Relatively mainstream | 4810 | 53974 |
| 15 | Crotonylation | Relatively mainstream | 3428 | 38533 |
| 16 | Myristoylation | Relatively mainstream | 2743 | 30842 |
| 17 | Farnesylation | Relatively mainstream | 2057 | 23132 |
| 18 | Geranylgeranylation | Relatively mainstream | 1714 | 19276 |
| 19 | S-nitrosylation | Relatively mainstream | 4110 | 46263 |
| 20 | Glutathionylation | Relatively mainstream | 3086 | 34697 |
| 21 | Hydroxylation | Relatively mainstream | 4118 | 46263 |
| 22 | Oxidation | Relatively mainstream | 3428 | 38553 |
| 23 | Deamidation | Relatively mainstream | 2743 | 30822 |
| 24 | Sulfation | Relatively mainstream | 1701 | 19296 |
| 25 | Citrullination | Relatively mainstream | 1727 | 19270 |
| 26 | Amidation | Relatively mainstream | 1371 | 15427 |
| 27 | GPI anchor | Relatively mainstream | 1028 | 11566 |
| 28 | Malonylation | Rare | 1886 | 21204 |
| 29 | Glutarylation | Rare | 1543 | 17349 |
| 30 | Lactylation | Rare | 2125 | 23902 |
| 31 | Formylation | Rare | 514 | 5783 |
| 32 | Butyrylation | Rare | 548 | 6168 |
| 33 | N-palmitoylation | Rare | 343 | 3896 |
| 34 | S-diacylglycerol | Rare | 309 | 3471 |
| 35 | C-linked glycosylation | Rare | 340 | 3813 |

| | | | | |
|---|---|---|---|---|
| 36 | Gamma-carboxyglutamic acid | Rare | 449 | 5012 |
| 37 | Sulfoxidation | Rare | 309 | 3473 |
| 38 | Nitration | Rare | 411 | 4646 |
| 39 | Pyrrolidone-carboxylic acid | Rare | 274 | 3084 |
| 40 | Dephosphorylation | Rare | 206 | 2313 |
| | **Total** | | **348,903** | **3,902,925** |
| **No** | **Kinase-Specific Phosphorylation** | **Significance** | **Positive** | **Negative** |
| 1 | MAPK | Highest | 755 | 8026 |
| 2 | CDK | Highest | 1008 | 9924 |
| 3 | AGC | High | 1351 | 16824 |
| 4 | PKC | High | 689 | 7115 |
| | Total | | 3803 | 41889 |
| **No** | **PTM CrossTalk** | **-** | **Positive** | **Negative** |
| 1 | **-** | **-** | **262** | **12468** |
| **No** | **Enzyme kinetic parameters** | **-** | **Sequence** | |
| 1 | $K_{cat}$ | | 23917 | |
| 2 | $K_m$ | | 41174 | |
| 3 | $K_i$ | | 11929 | |
| | **Total** | | **77020** | |

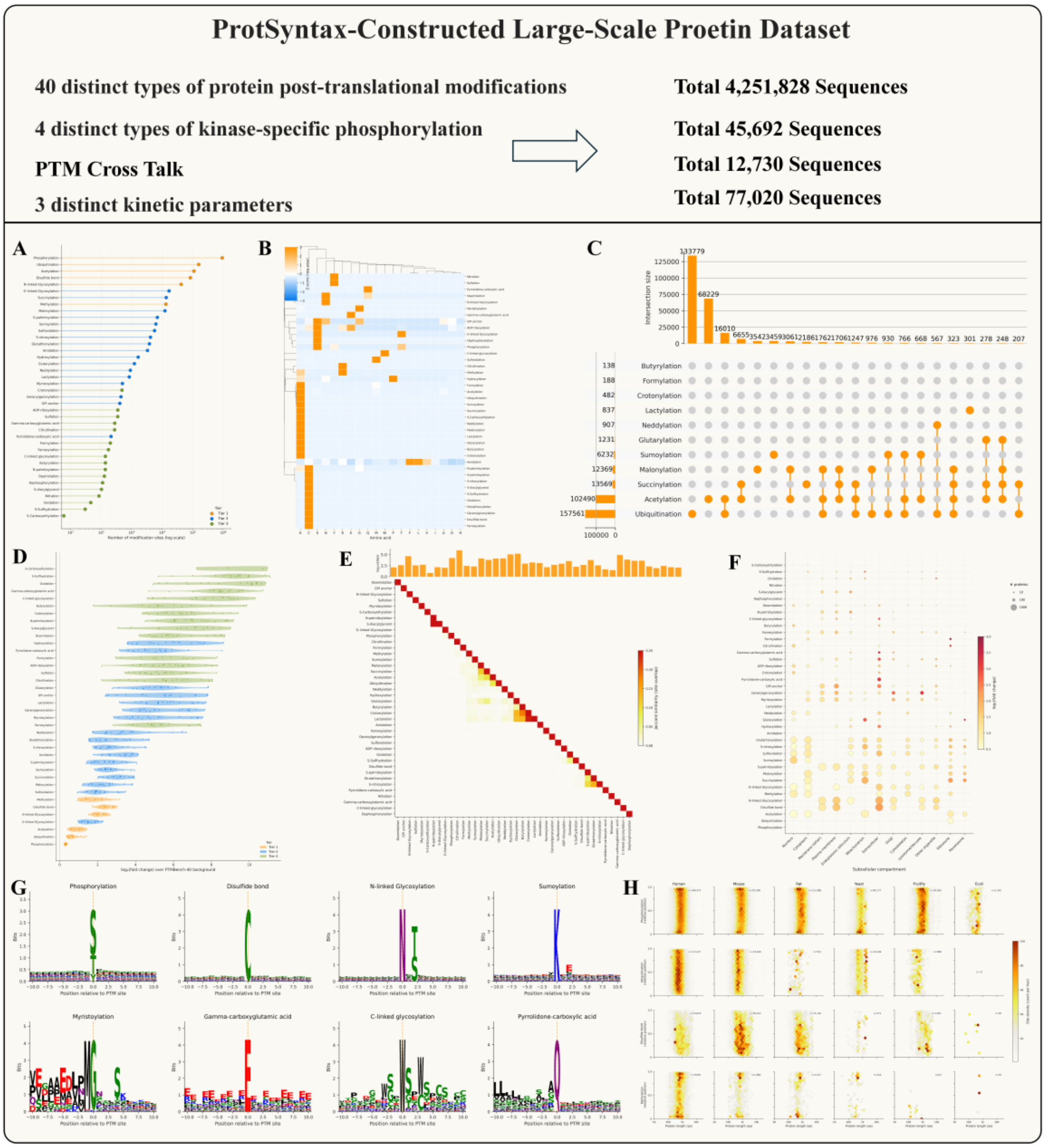


Figure S1. Dataset characterization and systematic analysis of PTM annotations. This figure summarizes the composition, sequence preferences, functional associations and evolutionary distribution of the curated post-translational modification (PTM) dataset. (A) Distribution of different PTM types in the dataset. (B) Biclustering heatmap showing amino acid preference patterns across PTM types. (C) Multi-label UpSet plot illustrating the overlap among lysine-centered modifications. (D) Violin and scatter plots summarizing Gene Ontology biological process (GO-BP) enrichment profiles associated with different PTM groups. (E) Lower-triangular heatmap depicting PTM co-occurrence patterns. (F) Bubble plot showing subcellular compartment enrichment across PTM types. (G) Motif logo analysis revealing sequence preferences surrounding representative PTM sites. (H) Hexbin density plots comparing the distributional patterns of four representative PTM types across six species.

Comprehensively, these 8 subplots (Figure S1) demonstrate that the dataset is not only extensive in scale but also exhibits high fidelity to authentic biological principles and complex modification networks. First, the data presents a natural "imbalance" in both scale and species distribution that accurately reflects real-world biological contexts; for instance, phosphorylation spans six orders of magnitude to maintain absolute dominance, while the ecological niches for species-specific modifications—such as those unique to E. coli, sea hares, and legumes—are precisely preserved. Second, the dataset possesses exceptionally high evidence quality and structural fidelity, with newly incorporated rare modifications almost entirely supported by gold-standard manual annotations, and spontaneously clustered amino acid preferences and sequence motifs perfectly aligning with established chemical mechanisms. Furthermore, the visualizations profoundly reveal the intricate networks and crosstalk among PTMs, capturing not only classic competitive switch mechanisms at identical sites (e.g., between acetylation and ubiquitination) but also clearly illustrating the highly complex characteristics of multiple overlapping modifications on lysine (K) residues. Finally, these modifications exhibit strong spatial distribution patterns and explicit functional indications; the precise functional enrichment of various PTMs within specific subcellular compartments (such as GPI anchors at the plasma membrane), along with the spatial positional preferences of modification sites across proteins of varying lengths (such as the N-terminal enrichment of methylation), collectively provide robust evidence for the biological validity of this dataset.

#### 2.1 Supplementary discussion: Rationale for the training scale of ProtSyntax

The four-million-scale PTM corpus used in this study should not be directly compared with the sequence collections used for unsupervised pretraining of general-purpose protein language models. Models such as the ESM family are trained on tens of millions to billions of largely unannotated protein sequences to acquire broad evolutionary, biochemical and structural regularities that support diverse downstream tasks. ProtSyntax has a different scope: it is a domain-specialized protein language model designed to resolve the regulatory syntax of PTMs and their functional consequences, rather than a universal model intended to reconstruct the entire protein sequence distribution. Its training corpus therefore comprises densely supervised residue-level examples spanning 40 PTM classes, together with kinase-specific phosphorylation, PTM crosstalk and enzyme kinetic annotations. Each example provides substantially more task-specific information than an unannotated sequence used in a conventional masked-language-modeling objective. Accordingly, the relevant quantity is not only the number of protein sequences, but also the density, diversity and biological organization of the available supervision. To our knowledge, this corpus represents one of the largest systematically curated PTM-supervised resources assembled for model development.

Importantly, ProtSyntax does not attempt to learn general protein biology de novo from these PTM annotations alone. ESM-C and SaProt are used as frozen upstream encoders, allowing ProtSyntax to inherit sequence-level evolutionary representations and structure-conditioned protein features acquired through large-scale unsupervised pretraining. The trainable ProtSyntax architecture is initialized and optimized on the PTM-centered objectives, but its input representation is already grounded in general protein knowledge. The current strategy is therefore more accurately described as training a PTM-specific architecture from random initialization on top of frozen pretrained protein representations, rather than pretraining an entirely independent protein language model from raw amino acid sequences. Conceptually, this corresponds to the proposed two-stage strategy: broad level-0 protein knowledge is first acquired by general-purpose PLMs, after which the four-million-scale annotated corpus is used to specialize that knowledge toward PTM chemistry, recognition context, structural permissiveness, regulatory crosstalk and protein-level functional effects.

The four-million-scale corpus should consequently not be interpreted as evidence that complex PTM regulation is intrinsically easy to learn or that large-scale general protein pretraining is unnecessary. Instead, ProtSyntax combines transferred protein knowledge with information-dense PTM supervision, explicit three-dimensional constraints and multi-task functional learning. Long-range and structurally mediated regulatory relationships are not inferred solely from the frequency of local PTM labels: they are supported by pretrained sequence and structure representations, modeled through bidirectional context propagation and geometric attention, and further constrained by full-length enzyme kinetic supervision. We assessed whether this specialization extends beyond conventional site-pattern recognition through homolog-separated evaluation, structure-matched decoy discrimination, rare-PTM transfer, crosstalk reconstruction and PTM-to-function coupling. These evaluations provide empirical evidence that the model uses transferable sequence–structure–function relationships, while avoiding the stronger and unsupported claim that four million PTM examples are sufficient to relearn the complete protein language from first principles.

### 3.Methods

#### 3.1 Bio-RoPE positional encoding

Bio-RoPE was introduced to make positional encoding compatible with the biochemical nature of PTM syntax. In conventional protein language models, rotary positional encoding mainly provides a relative ordering mechanism, implicitly assuming that positions with the same index share the same phase structure regardless of residue identity. This assumption is suboptimal for PTM modeling, because modification propensity is jointly shaped by residue order, local secondary-structure periodicity, side-chain chemistry and long-range protein context. Bio-RoPE therefore reformulates positional phase as a biologically modulated representation rather than a purely geometric index. Its structural-periodic channels encode canonical residue periodicities associated with strand-like, 3_10-helical, α-helical and π-helical arrangements, allowing the model to sense recurring spatial patterns that often organize enzyme recognition motifs or modification-prone microenvironments. Its physicochemical channels further introduce residue-dependent phase offsets derived from hydrophobicity, helical propensity and side-chain steric bulk, enabling residues occupying the same sequence position to receive different rotary phases according to their biochemical compatibility. The remaining standard RoPE channels preserve long-range relative-position extrapolation. Through this decomposition, Bio-RoPE embeds motif order, structural rhythm and amino-acid chemistry into the attention phase space, allowing ProtSyntax to distinguish chemically meaningful PTM syntax from positionally similar but biologically implausible residue contexts.

Given a hidden representation $x_i$ at residue position $i$, Bio-RoPE partitions the feature dimension into three channel groups: structural-periodic channels, physicochemical phase-modulation channels and standard long-range RoPE channels. The structural-periodic channels encode canonical residue periodicities observed in protein secondary structures, using the period set $P = \{2.0,3.0,3.6,5.1\}$, corresponding to extended strand-like periodicity, $3_{10}$-helices periodicity, $\alpha$-helices periodicity and $\pi$-helices periodicity. For a period $p_j \in \mathcal{P}$, the structural phase is defined as:

$$\theta_{i,j}^{\text{per}} = i \cdot \frac{2\pi}{p_j}, p_j \in \mathcal{P} \tag{1}$$

The physicochemical channels introduce amino-acid-dependent phase offsets. For residue identity $a_i$, ProtSyntax retrieves a frozen descriptor vector $r(a_i)$ encoding hydrophobicity, helical propensity and side-chain steric bulk. The descriptor is expanded across the physicochemical channels and added to the RoPE base phase:

$$\theta_{i,j}^{\text{phy}} = i \cdot \omega_j^{\text{phy}} + \rho_j(a_i) \tag{2}$$

where $\omega_j^{\text{phy}}$ denotes the base rotary frequency and $\rho_j(a_i)$ is the residue-specific phase offset. The remaining channels retain the standard RoPE formulation, $\theta_{i,j}^{\text{std}} = i \cdot \omega_j^{\text{std}}$, to preserve long-range relative positional modeling.

The final phase vector is obtained by concatenating the three groups, $\theta_i = [\theta_i^{\text{per}}; \theta_i^{\text{phy}}; \theta_i^{\text{std}}]$, and is applied to query and key representations as:

$$\text{BioRoPE}(x_i) = x_i \odot \cos(\theta_i) + \mathcal{R}(x_i) \odot \sin(\theta_i) \quad (3)$$

where $\mathcal{R}(\cdot)$ denotes the half-dimensional rotation of paired feature dimensions. In this formulation, residues at the same sequence position can receive distinct phase modulations according to amino acid chemistry, allowing ProtSyntax to encode PTM-relevant sequence syntax together with local structural periodic priors.

##### 3.2 Bi-Gated DeltaNet for bidirectional PTM context modeling

Bi-Gated DeltaNet was designed to propagate PTM-relevant sequence evidence across protein chains with linear sequence complexity while preserving bidirectional biological context. PTM sites are rarely determined by the central residue alone; rather, their modification status reflects the coordinated contribution of upstream and downstream motifs, local physicochemical patterns, domain boundaries and distal sequence cues. A unidirectional recurrent module can efficiently accumulate contextual information, but it may overemphasize one-sided motif evidence and fail to determine whether both sides of a candidate residue support the same regulatory event. Bi-Gated DeltaNet addresses this limitation by using two independents Gated DeltaNet cores that scan the protein in opposite directions, followed by a zero-parameter cross-gating mechanism that allows the forward and backward states to conditionally recalibrate each other at every residue. The recurrent core uses decay and writes gates to selectively retain informative historical context, suppress outdated or conflicting memory components and incorporate current residue information through efficient rank-one state updates. The bidirectional cross-gating step further acts as a syntax filter: PTM signals are strengthened when N-terminal and C-terminal contexts provide coherent evidence but weakened when a residue appears motif-compatible only from one side. This design enables ProtSyntax to model long-range protein context efficiently while maintaining residue-level sensitivity to sparse, context-dependent PTM signals.

Given an input protein representation X, respectively, each unidirectional Gated DeltaNet core first projects the input into multi-head query, key, and value representations:

$$Q = XW_q, K = XW_k, V = XW_v \quad (4)$$

where $W_q$, $W_k$, and $W_v$ are bias-free linear projection matrices. The projected features are reshaped into $H$ heads with head dimension D. To improve the numerical stability of the recurrent state update, the query and key vectors are processed by zero-centered RMS normalization followed by a SiLU activation:

$$\tilde{q}_{t,h} = \mathrm{SiLU}\left( \frac{q_{t,h}}{\| \frac{q_{t,h} \|_2}{\sqrt{D}} + \epsilon} \odot (1 + \gamma_q) \right) \tag{5}$$

$$\tilde{k}_{t,h} = \mathrm{SiLU}\left( \frac{k_{t,h}}{\| \frac{k_{t,h} \|_2}{\sqrt{D}} + \in} \odot (1 + \gamma_k) \right), \tilde{v}_{t,h} = \mathrm{SiLU}(v_{t,h}) \tag{6}$$

Here, $\gamma_q$ and $\gamma_k$ are initialized to zero, allowing the normalized representations to start from an approximately identity-scaled regime. This normalization reduces the influence of unusually large token activations on state writing, which is particularly important for maintaining stable contextual representations over long protein sequences.

At each residue position t and attention head h, the unidirectional core computes a decay gate and a write gate:

$$\alpha_{t,h} = \sigma(W_{\alpha,h} x_t + b_h^{\mathrm{struct}}), \beta_{t,h} = \sigma(W_{\beta,h} x_t) \tag{7}$$

where $\alpha_{t,h}$ controls the retention of historical information, $\beta_{t,h}$ controls the incorporation of the current residue, and $b_h^{\mathrm{struct}}$ is a learnable head-specific structural bias. This bias modulates the decay behavior of different heads, allowing the module to allocate some heads to short-range motif-sensitive patterns and others to longer-range sequence or domain-level dependencies. The model maintains a head-wise recurrent state matrix $S_{t,h}$, which is updated by a gated Delta rule:

$$S_{t,h} = S_{t-1,h}\left[\alpha_{t,h}\left(I - \beta_{t,h} \tilde{k}_{t,h} \tilde{k}_{t,h}^{\mathrm{T}}\right)\right] + \beta_{t,h} \tilde{v}_{t,h} \tilde{k}_{t,h}^{\mathrm{T}} \tag{8}$$

The first term selectively decays and corrects the previous state, whereas the second term writes the current residue information through a rank-one update. Compared with explicit global attention, this recurrent formulation updates the state by sequential scanning and therefore remains efficient for long protein sequences. The correction term associated with $\tilde{k}_{t,h}$ also suppresses outdated or conflicting memory components, enabling the state matrix to emphasize contextual features that are more informative for the current candidate PTM site. The unidirectional output at position *t* is then retrieved by querying the state matrix:

$$h_{t,h} = S_{t,h}\tilde{q}_{t,h} \tag{9}$$

The outputs from all heads are concatenated to produce the unidirectional hidden representation $h_{t,h}$. The above equation defines the new core gated DeltaNet recurrence. Bi-Gated DeltaNet applies this recurrent state evolution in both sequence directions using two independent Gated DeltaNet (GDN) cores:

$$\vec{h}_t = \mathrm{GDN}_{\mathrm{fwd}}(X)_t, \overleftarrow{\mathrm{h}}_t = \mathrm{GDN}_{\mathrm{bwd}}(X)_t \tag{10}$$

The backward core processes the reversed protein sequence from the C terminus to the N terminus and then reverses its output back to the original residue coordinates. As a result, each residue position receives two aligned representations: one summarizing the N-terminal context and the other summarizing the C-terminal context. This bidirectional formulation is particularly suitable for PTM prediction, because modification propensity often depends on the coordinated pattern of residues on both sides of the modified amino acid rather than on a single upstream or downstream motif alone. The two directional representations are fused using zero-parameter cross-gating:

$$\hat{h}_t^{\mathrm{fwd}} = \vec{h}_t \odot \sigma\left(\overleftarrow{\mathrm{h}}_t\right), \hat{h}_t^{\mathrm{bwd}} = \overleftarrow{\mathrm{h}}_t \odot \sigma\left(\vec{\mathrm{h}}_t\right) \tag{11}$$

This operation uses the backward state to gate the forward state and the forward state to gate the backward state, without introducing additional learnable parameters. Therefore, bidirectional evidence is not merely aggregated but conditionally recalibrated at each residue position. In the context of PTM site prediction, this mechanism can reduce false-positive signals arising from one-sided incidental motifs and strengthen residues for which both flanking contexts provide coherent modification-related evidence. The cross-gated directional states are concatenated and projected back to the model hidden dimension:

$$h_t^{\mathrm{fused}} = [\hat{h}_t^{\mathrm{fwd}}; \hat{h}_t^{\mathrm{bwd}}], z_t = W_o h_t^{\mathrm{fused}}, y_t = z_t \odot \mathrm{SiLU}\left(W_g x_t\right) \tag{12}$$

Finally, the fused representation is filtered by an input-conditioned output gate. $W_o$ and $W_g$ denote the output projection and output-gating matrices.

##### 3.3 Geometric Gated Attention for structure-constrained PTM reasoning

Geometric Gated Attention was introduced to model the three-dimensional microenvironmental component of PTM syntax, which cannot be fully resolved from linear sequence representations. Many PTM events depend on spatially proximal residues, backbone orientation, conformational accessibility and domain-level organization; therefore, a residue that is chemically compatible in sequence space may still be inaccessible, misoriented or structurally unsupported in the folded protein. Rather than treating structure as an auxiliary feature concatenated to sequence embeddings, Geometric Gated Attention injects residue-frame-based geometry directly into the attention logits. For each attention head, the module generates learnable local geometric probe points for query and key residues, maps them into global coordinates using backbone-derived rigid frames and converts their pairwise distances into a head-specific geometric penalty. This penalty suppresses semantically plausible but spatially inconsistent residue interactions, while preserving high attention for residue pairs supported by both sequence context and physical proximity. A query-dependent sigmoid gate further filters redundant structural–semantic responses after attention aggregation. Importantly, the geometric point projections are initialized to zero, allowing the module to behave like standard semantic attention at the beginning of training and to acquire structural selectivity progressively under PTM supervision. This design makes GGA particularly suitable for distinguishing motif-like false positives from genuinely permissive PTM microenvironments.

Given an input representation X, GGA first computes standard multi-head query, key and value projections:

$$Q = XW_Q, K = XW_K, V = XW_V \tag{13}$$

For each residue $i$, attention head $h$ and probe index $m$, GGA further predicts local query-side and key-side probe points $p_{ihm}^{Q}$ and $R_j p_{jhm}^{K}$. These local points are transformed into the global structural coordinate system using residue-specific rigid frames $(R_i, t_i)$ derived from the protein backbone:

$$\tilde{p}_{ihm}^{Q} = R_i p_{ihm}^{Q} + t_i, \tilde{p}_{jhm}^{K} = R_j p_{jhm}^{K} + t_j \tag{14}$$

This design enables each attention head to move beyond sequence-level semantic association and instead perceive spatial proximity and conformational relationships within a learnable three-dimensional probe space. GGA then injects the squared Euclidean distance between probe points into the attention logits as a geometric penalty:

$$S_{hij} = \frac{Q_{hi}^{\mathrm{T}} K_{hj}}{\sqrt{d_h}} - \mathrm{softplus}(\gamma_h) \sum_{m=1}^{P} \left\| \tilde{p}_{ihm}^{Q} - \tilde{p}_{jhm}^{K} \right\|_2^2 \sqrt{\frac{2}{9P}} \tag{15}$$

Here, $\gamma_h$ is a learnable geometric scaling parameter for the hhh-th attention head, constrained to be positive by the Softplus function. This mechanism suppresses residue pairs that are semantically related but geometrically inconsistent, while allowing residue pairs that are coherent in both sequence semantics and structural space to receive higher attention weights. Consequently, GGA converts attention from a purely sequence-driven residue interaction model into a joint semantic–structural modeling scheme, which is better suited for capturing the functional microenvironment of PTM sites shaped by local motifs, spatial contacts, and conformational accessibility.

After obtaining the structure-corrected attention weights, GGA aggregates the value vectors and applies a query-dependent head-wise sigmoid output gate:

$$G_{ih} = \sigma\left(X_i W_g + b_g\right), \hat{O}_{ih} = G_{ih} \cdot O_{ih} \tag{16}$$

Operating after contextual aggregation, this gate dynamically regulates the contribution of each attention head to filter out redundant structural–semantic responses irrelevant to PTM site discrimination. The gated outputs from all heads are then concatenated and projected back to the original feature dimension to form the input for the downstream prediction module. To stabilize optimization during this process, GGA initializes the three-dimensional point projections to zero, effectively making the module behave like standard semantic attention in the early stages. As training progresses, the geometric probes and head-specific penalties are learned under the PTM prediction objective to capture modification-related spatial conformational patterns.

#### 3.4 PACE-Nash Loss

PACE-Nash was developed to align residue-level PTM syntax with protein-level functional supervision under a unified multi-objective training framework. PTM-centered protein modeling presents several coupled optimization challenges: positive modification sites are sparse relative to chemically compatible negatives, different PTM types exhibit non-independent biochemical relationships, enzyme kinetic measurements contain heteroscedastic uncertainty and residue-level classification objectives may conflict with full-length functional regression objectives. PACE-Nash addresses these issues by integrating asymmetric focal classification, correlation-aware contrastive learning, evidential kinetic regression, physicochemical manifold regularization and Nash bargaining-based adaptive task weighting. The asymmetric focal term reduces the dominance of abundant negative residues, whereas the contrastive component encourages samples with related PTM patterns to occupy nearby regions in latent space, promoting transfer across chemically related modifications. The evidential regression branch predicts both the mean and uncertainty of log-transformed Kcat, Km and Ki, allowing noisy kinetic labels to contribute in proportion to their confidence. In parallel, physicochemical manifold regularization constrains latent distances to remain consistent with protein-level biochemical descriptors. Rather than manually assigning fixed loss weights, PACE-Nash adaptively balances these objectives by seeking a Pareto-efficient bargaining solution over shared model gradients. This enables ProtSyntax to learn representations that remain sensitive to local PTM grammar while retaining global functional information.

For PTM-related classification, PACE-Nash combines correlation-aware supervised contrastive learning with asymmetric focal classification. Given a batch of normalized protein representations $z_i$ and multi-label PTM annotation vectors $y_i$, the biological relatedness between samples *i* and *j* is measured using Jaccard overlap:

$$w_{ij} = \frac{|y_i \cap y_j|}{|y_i \cup y_j| + \epsilon} \tag{17}$$

Pairs with $w_{ij}$ below a threshold $\tau$ are ignored. The correlation-aware contrastive loss is then defined as:

$$\mathcal{L}_{\text{con}} = -\frac{1}{N} \sum_{i} \frac{\sum_{j \neq i} w_{ij} \log \frac{\exp\left(\frac{z_i^{\mathrm{T}} z_j}{T}\right)}{\sum_{k \neq i} \exp\left(\frac{z_i^{\mathrm{T}} z_k}{T}\right)}}{\sum_{j \neq i} w_{ij} + \epsilon} \tag{18}$$

where *T* is the temperature parameter. This objective encourages proteins or residue windows with related PTM patterns to occupy neighboring regions in the latent space. The contrastive term is combined with an asymmetric focal binary cross-entropy loss to reduce domination by abundant negative labels:

$$\mathcal{L}_{\text{PTM}} = \mathcal{L}_{\text{con}} + \mathcal{L}_{\text{focal}} \tag{19}$$

For enzyme kinetic prediction, ProtSyntax predicts both the mean μ\muμ and log-variance $\log\sigma^2$ of the log-transformed kinetic target. The regression loss is a Gaussian negative log-likelihood in log space with variance regularization:

$$\mathcal{L}_{\text{kin}} = \frac{1}{2}\left[\log(2\pi) + \log\sigma^2 + \frac{(y-\mu)^2}{\sigma^2}\right] + \lambda_{\text{var}} \parallel \sigma^2 \parallel_2 \tag{20}$$

This formulation allows the model to represent heteroscedastic experimental uncertainty in kinetic measurements.

To preserve coarse physicochemical topology in the learned representation space, PACE-Nash includes a physicochemical manifold penalty. Let $d_z(i,j)$ denote the Euclidean distance between latent representations and $d_p(i,j)$ denote the distance between precomputed physicochemical descriptor vectors. The penalty is:

$$\mathcal{L}_{\text{phys}} = \frac{1}{N^2 - N} \sum_{i \neq j} \max\left(0, m d_p(i,j) - d_z(i,j)\right) \quad (21)$$

where $m$ is a margin. This term discourages representation collapse between proteins with substantially different physicochemical profiles.

The three task losses are assembled as: $\mathbf{L} = [\mathcal{L}_{\text{PTM}}, \mathcal{L}_{\text{kin}}, \mathcal{L}_{\text{phys}}]$. Instead of assigning fixed manual task weights, PACE-Nash uses a Nash bargaining strategy to determine adaptive coefficients. Gradients from each loss are computed on shared parameters, flattened into task-specific gradient vectors and used to construct a gradient inner-product matrix. The task coefficients $a_k$ are constrained to the probability simplex and updated to improve the joint utility of all objectives. The final optimization target is:

$$L_{PACE-Nash} = \sum_{k=1}^{K} a_k L_k, \quad \sum_{k=1}^{K} a_k = 1 \quad (22)$$

This adaptive weighting scheme reduces destructive gradient interference between PTM classification, enzyme kinetic regression and physicochemical regularization, enabling ProtSyntax to learn a shared representation that remains sensitive to residue-level PTM syntax while preserving global protein functional information.

### 4.Results

#### 4.1 Ablation Study

For the ablation baselines, we selected representative modules that are widely used in contemporary large language models, including standard RoPE and cross-entropy loss, as well as Gated DeltaNet and Gated Attention from the Qwen model family. These components represent advanced and competitive design choices in current large-model architectures, thereby providing strong reference baselines rather than weak substitutes. The consistent improvements achieved by ProtSyntax-specific modules over these competitive counterparts further demonstrate the effectiveness and superiority of the ProtSyntax design. The data presented in Tables S2 and S3 provide the original ablation results for ProtSyntax.

Table S2. Ablation results for ProtSyntax

| Model | MCC | AUC | AP |
|---|---|---|---|
| Bio-RoPE | 0.893 | 0.921 | 0.937 |
| RoPE | 0.865 | 0.896 | 0.905 |
| Bi-Gated DeltaNet | 0.893 | 0.921 | 0.937 |

| | | | |
|---|---|---|---|
| Gated DeltaNet | 0.850 | 0.882 | 0.888 |
| Geometric Gated Attention | 0.893 | 0.921 | 0.937 |
| Gated Attention | 0.854 | 0.879 | 0.884 |
| PACE-Nash Loss | 0.893 | 0.921 | 0.937 |
| Cross-Entropy Loss | 0.869 | 0.900 | 0.912 |

We next examined the robustness of ProtSyntax to structural uncertainty by stratifying the phosphorylation test set into three pLDDT ranges: <50, 50–70 and >70. Importantly, these results should be interpreted in the context of confidence-aware structural masking. ProtSyntax incorporates frozen SaProt representations, for which the 3Di structural token of each residue with pLDDT <70 is replaced by a structure-mask token while its amino-acid identity is retained. Consequently, low-confidence regions contribute primarily sequence-derived information rather than potentially unreliable structural states. ProtSyntax applies a complementary strategy in its geometric branch: residues with unreliable coordinates are excluded from geometric attention, allowing the corresponding interactions to revert to sequence-based semantic attention. Thus, the performance observed in the lower-pLDDT groups does not indicate that ProtSyntax extracts precise geometric information from uncertain structures. Instead, it demonstrates that the model can suppress unreliable structural evidence while preserving residue chemistry, motif organization and bidirectional long-range sequence context. ProtSyntax consistently outperformed PTMGPT2 in both MCC and AP across all three pLDDT ranges. Even in the pLDDT <50 group, ProtSyntax achieved an MCC of 0.814, representing a 12.6-percentage-point improvement over PTMGPT2. Performance further increased with structural confidence, reaching an MCC of 0.927 and an AP of 0.961 in the pLDDT >70 group, where both SaProt-derived structural representations and explicit geometric constraints were more broadly available.

Table S3. Performance comparison of ProtSyntax across different pLDDT ranges

| pLDDT | Model | MCC | AP |
|---|---|---|---|
| <50 | ProtSyntax | 0.814 | 0.829 |
| | PTMGPT2 | 0.688 | 0.694 |
| 50-70 | ProtSyntax | 0.860 | 0.892 |
| | PTMGPT2 | 0.767 | 0.805 |
| >70 | ProtSyntax | 0.927 | 0.961 |
| | PTMGPT2 | 0.842 | 0.883 |

**4.2 Comparison of model inference speed**

ProtSyntax incorporates substantial engineering optimizations, in part by drawing on the implementation principles and code architecture of Alibaba's open-source large language model Qwen3-Next. These optimizations markedly improve the inference efficiency of ProtSyntax. Assuming that the feature embeddings required by each model, such as ESM-2, ESM-C and PDB-derived structural features, had been precomputed, we quantitatively compared the inference speed of ProtSyntax, PTMGPT2 and AstraPTM2 under identical computational settings on a single NVIDIA H100 GPU. Three representative inference scenarios were evaluated: PTM site-centered window prediction using a 55-aa input, a single full-length forward pass for a 300–500-aa protein, and whole-protein scanning of candidate PTM sites in a 500-aa protein. The results are reported in Table S4. ProtSyntax achieved the fastest inference speed across all three scenarios, demonstrating that it not only substantially improves predictive performance over existing models but also achieves a favorable balance between model complexity and computational efficiency.

Table S4. Comparison of inference speed across models under three representative scenarios. Where --- indicate that the corresponding model does not natively support the given inference scenario.

| Model | Scene | Seq/s |
|---|---|---|
| ProtSyntax | PTM site-centered window(55aa) | 3283 |
| PTMGPT2 | | 1150 |
| AstraPTM2 | | 3092 |
| ProtSyntax | Single full-length forward pass for a 300–500-aa protein | 247 |
| PTMGPT2 | | --- |
| AstraPTM2 | | 212 |
| ProtSyntax | Whole-protein scanning of candidate PTM sites in a 500-aa protein | 78 |
| PTMGPT2 | | 22 |
| AstraPTM2 | | 51 |

##### 4.3 Supplementary evaluation of ProtSyntax transferability to low-resource PTM types

We report the AUC values of ProtSyntax for each of the eight rare PTM types under 0-, 10- and 50-positive-sample transfer settings, providing a detailed supplement to the main text. The results are shown in Figure S2.

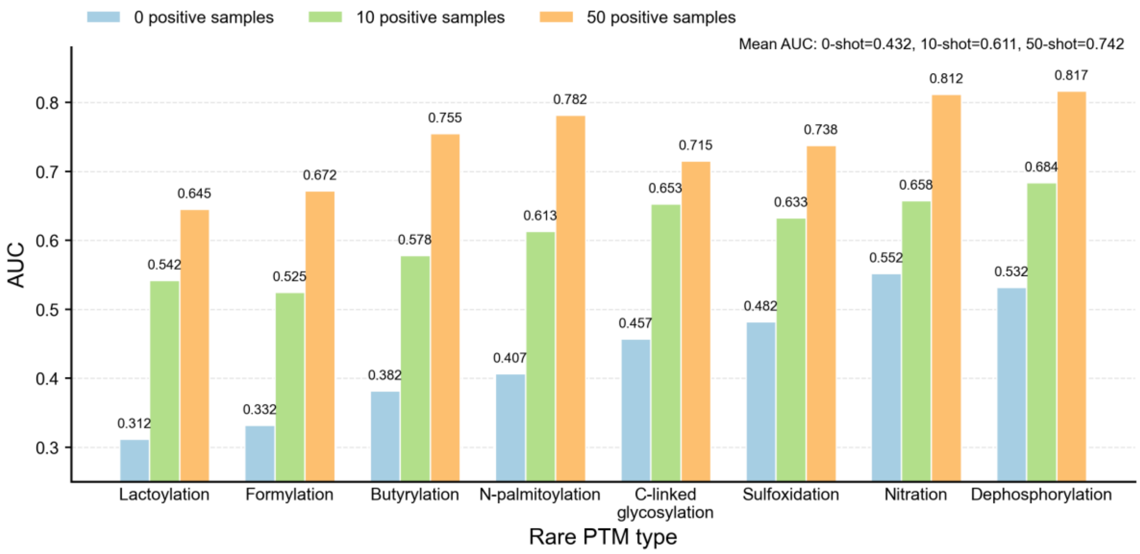


Figure S2. Results of the sample-transfer experiments for ProtSyntax across eight rare PTM types.

##### 4.4 Additional experiments supporting ProtSyntax as a PTM-aware protein language model

Evolutionary conservation within protein sequences serves as a robust indicator of functional significance. PTMs occurring at highly conserved sites often denote evolutionarily ancient and essential regulatory mechanisms vital for core signaling and cellular homeostasis, whereas those in rapidly evolving regions may represent species-specific adaptations or noise. Recognizing that prioritizing conserved sites is key to identifying high-confidence regulatory switches, we conducted a comparative analysis using the established ConSurf tool to partition phosphorylation data based on conservation scores (thresholds >0.8 and >0.9). As shown in Figure S3.A, benchmarking results demonstrate that ProtSyntax achieved the highest accuracy across both conservation thresholds. This superior performance not only underscores the model's capability in identifying critical conserved PTM sites but also validates its potential to advance from local site prediction to a system-level understanding of regulatory networks.

A major bottleneck in current research is predicting combinatorial PTM codes on individual proteins. We addressed this by selecting 20 representative "unseen" sequences containing multiple (⩾2) PTM types for comparative analysis against state-of-the-art generalist models, MTPrompt-PTM and PTMGPT2. Adopting a strict joint-probability threshold (>0.7 for all sites), ProtSyntax demonstrated exceptional robustness. As illustrated in Figure S3B, our model attained a 90% success rate, surpassing MTPrompt-PTM (65%) and PTMGPT2 (50%) by a wide margin. This performance disparity highlights ProtSyntax's advanced capability to capture latent correlations between different modifications, confirming its paradigm shift from a static site predictor to a model capable of systemic network comprehension.

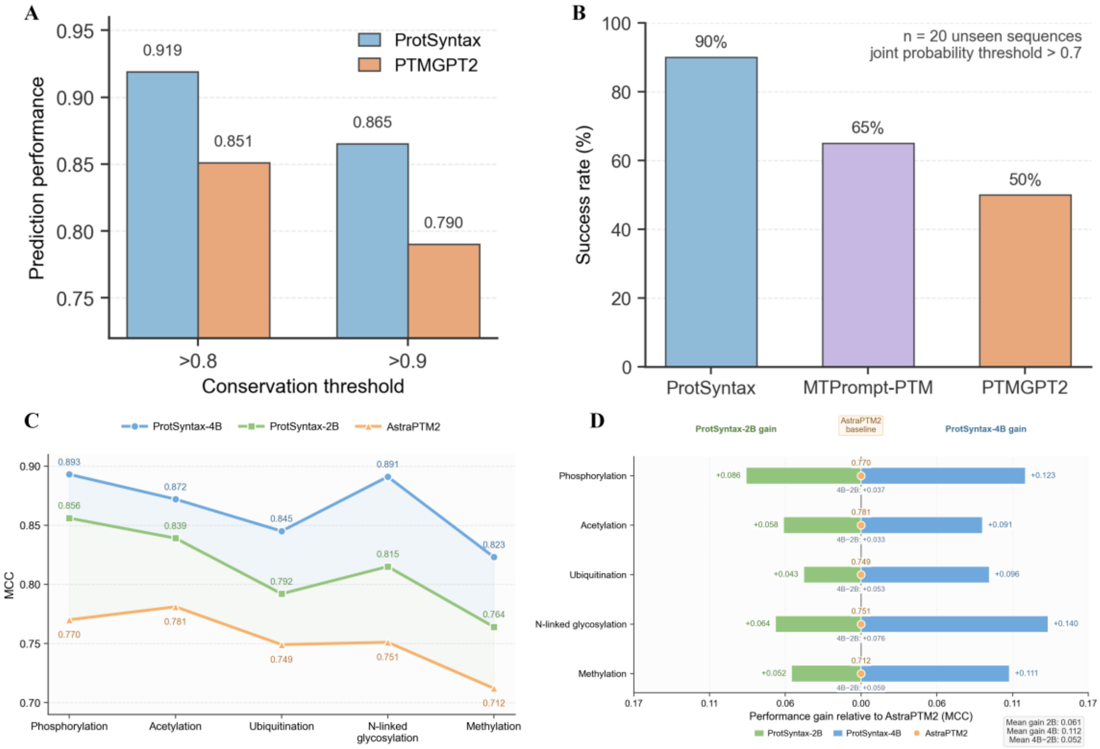


Figure S3. Performance comparison on additional validation tasks and across the ProtSyntax model family. (A,B) Visualization of the comparative results on additional validation tasks. (C,D) Performance comparison between different members of the ProtSyntax model family.

ProtSyntax is released in two configurations: ProtSyntax-4B, which is reported in the main text, and a lightweight variant, ProtSyntax-2B. ProtSyntax-2B is designed to improve deployability and inference efficiency while incurring only a modest reduction in predictive accuracy. We therefore report a direct performance comparison between these two model scales in Figure S3.C-D. Together, these two configurations provide flexible options for researchers with different computational resources and application requirements.

##### 4.5 Pathway-scale mapping of PTM syntax in Alzheimer’s disease

As an additional case study, we used Alzheimer’s disease (AD)—in which tau hyperphosphorylation promotes neurofibrillary-tangle formation, multisite modification of APP regulates amyloid-β processing and secretase components such as PSEN1 are phosphoregulated—to examine whether ProtSyntax could organize disease-associated PTM syntax at proteome scale. Starting from the KEGG AD pathway (hsa05010), gene-symbol mapping and redundancy removal yielded 383 reviewed human UniProt proteins. ProtSyntax scored all chemically compatible residues across 19 PTM classes, spanning phosphorylation, acetylation, methylation, glycosylation, ubiquitination and ubiquitin-like modifications, generating 214,479 candidate residue–PTM pairs. Of these, 86,219 scored ≥0.5 (40.2%) and 18,011 passed the stringent threshold of ≥0.8 (8.4%), substantially exceeding the coverage achieved by PTMGPT2(Figure S4); every protein contained at least one positive-scoring site, with a median of 162 sites per protein. Functional and chemical stratification revealed a coherent AD-associated landscape. g: Profiler identified 593 significantly enriched terms (adjusted $P<0.05$), converging on autophagy, endoplasmic-reticulum membranes, glutamatergic and cholinergic synapses, calcium-channel regulation, PI3K–AKT signalling and receptor-tyrosine-kinase cascades, with MAPT represented in 9 of 13 selected pathways. Eighteen of the 19 PTM classes were significantly enriched among the pathway proteins, led by phospho-Ser/Thr with a 5.8-fold foreground-to-background enrichment. Kinase-PSSM analysis of 12,982 predicted phospho-Ser/Thr sites using 303 kinase models further showed that CMGC-group kinases and CK2—including GSK3, CDK5 and DYRK1A—accounted for 41% of the 82 predicted MAPT phosphosites, compared with 18% across the complete atlas, recapitulating established tau-kinase biology. Against a curated reference set of 10,507 sites from UniProt and PhosphoSitePlus, recall reached 91% for N-linked glycosylation, 84% for Lys methylation, 80% for Lys acetylation and Tyr phosphorylation, and 57% for Ser/Thr phosphorylation; protein-level recall was 63%, 53% and 44% for MAPT, APP and PSEN1, respectively. ProtSyntax recovered the established AT8 and PHF-1 tau phospho-epitopes and the disease-associated K280/K281 acetylation cluster, while ten PTM classes largely absent from previous prediction frameworks contributed 23,297 additional candidates. Collectively, this residue-resolution atlas integrates pathway enrichment, PTM burden, kinase-family attribution and external-reference validation into a prioritized hypothesis space for targeted proteomics, site-directed mutagenesis and therapeutic-target nomination.

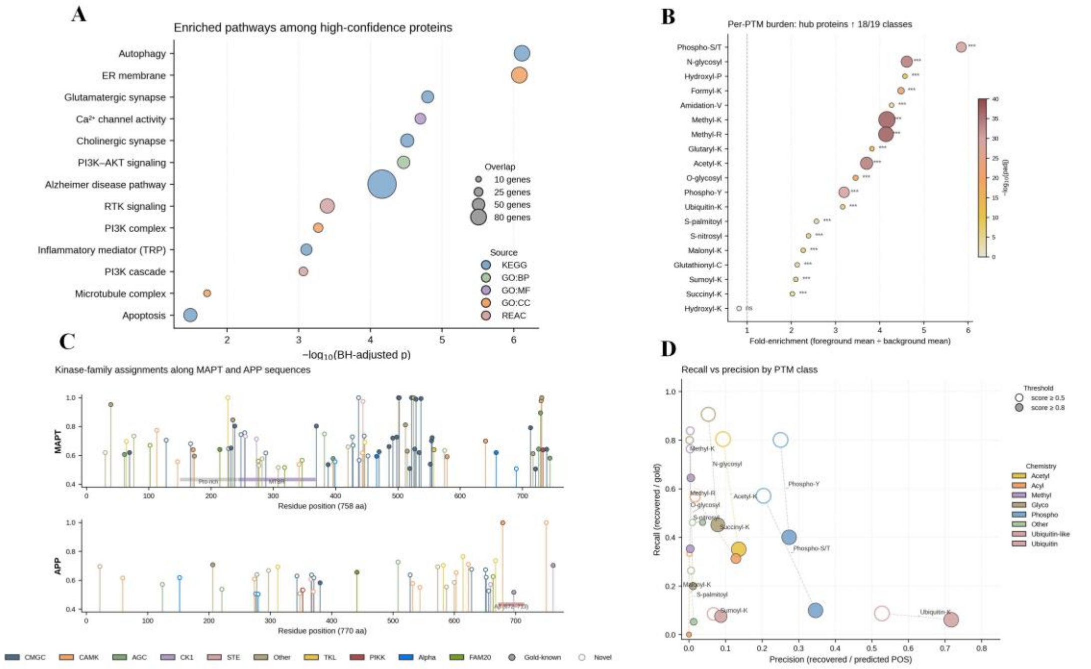


Figure S4. PTM landscape analysis and visualization. (A) Pathway enrichment bubble plot. Functional enrichment of high-confidence AD proteins (top 25%, n=96) against a 383-protein background, displaying 13 curated AD-relevant pathways. The y-axis ranks the adjusted p-value (padj), and bubble size reflects gene count. Autophagy and the ER membrane rank highest (padj$\approx 8\times10^{-7}$), highlighting a pathological spectrum involving synapses, calcium signaling, PI3K/AKT, microtubules, and apoptosis. (B) PTM chemical burden dot plot. Comparison of PTM site counts between foreground and background proteins across 19 categories. Dot size indicates the foreground mean, and color intensity reflects $-\log_{10}$(padj). Eighteen categories were significantly enriched; Phospho-S/T led with a ~5.8× enrichment, followed by Methyl-K/R, Acetyl-K, and N-glycosylation, demonstrating a substantial multi-modification burden on AD-associated proteins. (C) Kinase family mapping for MAPT and APP. Lollipop plots of high-confidence Phospho-S/T sites along MAPT and APP sequences. Colors indicate the top-predicted kinase family. Solid dots mark gold-standard sites (UniProt/PhosphoSitePlus), while hollow dots represent novel predictions. Bottom shaded bands denote key domains (MAPT Pro-rich/MTBR, APP Aβ). The plot emphasizes the dominant role of the CMGC kinase family in MAPT phosphorylation. (D) Recall-precision scatter plot across PTMs. Evaluation of 15 PTM categories with gold-standard coverage (10,507 sites). Hollow and solid dots denote score thresholds of $\geq 0.5$ and $\geq 0.8$, respectively. The results exhibit a discovery-oriented "high-recall, low-precision" profile, where precision acts as a conservative lower bound due to incomplete gold standards. Notably, Phospho-Y, N-glycosylation, and Acetyl-K achieved a strong balance of both metrics.

##### 4.6 ProtSyntax-guided prioritization of kinase–site–substrate intervention axes in

**temozolomide-resistant glioblastoma**

To assess whether ProtSyntax could translate disease-associated PTM alterations into actionable therapeutic hypotheses, we used temozolomide-resistant glioblastoma as a case study. Differential proteomic and phosphoproteomic profiles from resistant and sensitive samples were integrated at the protein level, yielding 422 candidate resistance-associated proteins[9]. ProtSyntax systematically scored all chemically compatible residues and identified 7,335 high-confidence PTM sites, of which 1,126 were located within or near catalytic pockets, protein–protein interaction interfaces, regulatory domains or resistance-associated mutation regions. We then constructed a residue-level prioritization framework integrating prediction confidence, disease-associated omic changes, predicted functional impact, PTM-crosstalk strength and pathway centrality, with scores normalized within each PTM class to reduce abundance and residue-frequency biases. This analysis retained 368 high-priority sites across 74 proteins, preferentially associated with receptor-tyrosine-kinase signalling, PI3K–AKT and MAPK pathways, cell-cycle regulation, DNA-damage responses and apoptosis. In silico perturbation further indicated that 143 of these sites could substantially alter at least one additional PTM event, suggesting that they may function as regulatory hubs within the resistance-associated network.

High-priority phosphosites were subsequently assigned to upstream kinases by jointly considering ProtSyntax phosphorylation scores, kinase–substrate specificity, kinase activity changes, substrate-network coverage and pathway concordance. Ninety-six candidate kinases were ranked after further incorporating druggability and normal-tissue essentiality. MAPK1, CDK2, AKT1, SRC and CSNK2A1 emerged as the five highest-ranked candidates; their predicted substrates encompassed 28 core resistance-associated proteins and collectively accounted for 39.2% of the prioritized phosphosites. Computational suppression of MAPK1-, CDK2- or AKT1-associated phosphorylation events reduced the predicted resistance-related PTM-network burden by 24.6–31.8%, substantially exceeding the mean reduction observed for matched random kinases (9.66%). Pairwise simulations indicated that combined MAPK1 and CDK2 blockade produced the strongest predicted network suppression while exerting comparatively limited effects on proteins unrelated to the resistant phenotype. The ranking remained stable across alternative confidence thresholds and bootstrap resampling, with MAPK1, CDK2 and AKT1 consistently retained among the leading candidates. Finally, residue-level evidence was consolidated into 16 candidate kinase–site–substrate intervention axes, including MAPK1–MYC, AKT1–GSK3B and CDK2–RB1 relationships, providing a focused set of hypotheses for targeted phosphoproteomics, site-directed mutagenesis and pharmacological perturbation. Collectively, this application demonstrates that ProtSyntax can reduce a complex disease-associated PTM landscape to a tractable set of mechanistically interpretable kinases, functional sites and substrate-specific intervention axes, thereby connecting disease omics with PTM mechanism analysis and drug-target prioritization.

#### 4.7 Supplementary notes on Sections 2.1.1–2.1.5 of the main text

##### 4.7.1 Task Description and Model Selection for Validation Experiments

To improve readability, Table S5 summarizes the comparison models included in each experiment described in Sections 2.1.1–2.1.5 of the main text.

The comparison models were selected according to their methodological relevance to each evaluation rather than imposing a uniform baseline set across all tasks. Fine-tuned ESM-C and SaProt were included as controlled general-purpose protein language model baselines to determine whether the observed gains could be attributed solely to large-scale pretrained sequence representations or structure-aware embeddings. Both models were adapted using the same task-specific training data and evaluated under protocols matched to ProtSyntax, thereby isolating the contribution of ProtSyntax-specific architecture and learning objectives. Except for these two fine-tuned foundation models, all remaining comparators were selected as the strongest task-specific baselines in their respective domains. AstraPTM2 represents a leading broad-spectrum, structure-aware PTM predictor and was therefore used for contextual recovery and structural microenvironment discrimination; PTMGPT2 provides a strong PTM-oriented generative baseline for low-resource transfer; DeepPCT and ProXTalk[10] are specialized state-of-the-art methods for PTM-crosstalk prediction; and CatPred serves as the leading reference for enzyme-kinetic modeling. This task-aligned design therefore benchmarks ProtSyntax against both powerful pretrained protein encoders and the strongest specialized methods available for each biological setting, while avoiding comparisons with models that do not support the corresponding prediction task.

Table S5. Comparison models included in each experiment described in Sections 2.1.1–2.1.5 of the main text.

| Task | ESM-C | SaProt | AstraPTM2 | PTMGPT2 | DeepPCT | ProXTalk | CatPred |
|---|---|---|---|---|---|---|---|
| PTM-language cloze recovery | √ | √ | √ | | | | |
| PTMmicroenvironment decoy discrimination | | | √ | | | | |
| PTM transfer | √ | √ | | √ | | | |

| | | | | | | |
|---|---|---|---|---|---|---|
| Combinatorial PTM crosstalk reconstruction | | | | √ | √ | |
| PTM-to-function coupling through enzyme kinetics | | | | | | √ |

Demonstrating that ProtSyntax understands PTM language requires evidence beyond improved site-classification accuracy, which may arise from memorizing residue frequencies, consensus motifs or dataset-specific correlations. We therefore selected five complementary experiments that interrogate progressively higher levels of PTM representation. The cloze-style tasks[11] test whether the model can reconstruct a missing PTM identity or select the correct modified residue from chemically compatible alternatives, thereby assessing its command of residue identity, motif order and contextual syntax. The structure-aware decoy experiment[12] examines whether this syntax is grounded in molecular geometry by requiring the model to reject sequence-plausible sites that lack an accessible and spatially coherent three-dimensional microenvironment. Low-resource transfer[13] provides a stringent test of abstraction: successful zero- and few-shot generalization indicates that knowledge is organized around transferable physicochemical and mechanistic relationships rather than modification-specific label frequency. PTM-crosstalk reconstruction[14] extends the evaluation from isolated events to conditional grammar, asking whether perturbing one modification alters the probability of another in a cooperative or antagonistic manner. Finally, PTM-to-function coupling[15] evaluates functional semantics by determining whether local modification perturbations are linked to quantitative changes in enzyme kinetics and whether these effects depend on proximity to catalytic, binding or regulatory regions. Collectively, these experiments span contextual completion, geometric permissiveness, cross-class compositional generalization, inter-PTM dependency and protein-level functional consequence. Their orthogonal design reduces the likelihood that success can be explained by any single shortcut and instead tests whether ProtSyntax has learned a coherent sequence–structure–function representation in which PTM identity, context, interaction and biological effect constitute interconnected levels of the same regulatory language.

###### 4.7.2 Detailed Description of the Experiments in Section 2.1 and 2.3

**1. Cloze-style recovery of residue-level PTM syntax**.We constructed the cloze-style evaluation from the protein-disjoint test partition of the 40-class PTM corpus described in Supplementary Section 2, which was assembled from experimentally supported annotations in UniProtKB, PhosphoSitePlus, dbPTM, iPTMnet and modification-specific literature. Records with ambiguous modification identities, uncertain residue coordinates, homology-only evidence or sequence mismatches were removed, and proteins were clustered with CD-HIT at 50% sequence identity before an 8:1:1 training–validation–test split using random seed 42. Two complementary tasks were formulated. In PTM-type recovery, the modification identity associated with an experimentally validated site was masked, whereas the central residue, its 55-residue sequence window and available AlphaFold2-derived geometry were retained; the model was required to recover the correct PTM among all 40 classes. The evaluation set was stratified by PTM type, and macro-averaged metrics were additionally reported to prevent abundant modifications from dominating the results. In PTM-site recovery, the PTM identity was supplied and protein segments containing at least three chemically compatible candidate residues were selected. Each candidate was evaluated using a centered window, and the experimentally modified residue was required to rank above the remaining unannotated candidates; residues annotated with the same PTM in any source were excluded from the negative set. Fine-tuned ESM-C+MLP, SaProt+MLP and AstraPTM2 were evaluated using identical protein partitions and candidate sets, with all hyperparameters selected exclusively on the validation set. PTM-type recovery was assessed by accuracy and macro-F1, whereas site recovery was evaluated by AUPRC and mean reciprocal rank. ProtSyntax achieved a type-recovery accuracy of 0.842 and a site-ranking AUPRC of 0.812. To exclude shallow motif or amino-acid-frequency shortcuts, we further generated counterfactual inputs by independently shuffling the residues flanking each central site while preserving amino-acid composition and by replacing the central residue with an amino acid incompatible with the target PTM. Ten shuffled controls were generated per site, and changes in the true-class logit were evaluated by protein-level bootstrap resampling. Both perturbations caused substantially larger score reductions in ProtSyntax than in the comparison models, whereas removal of Bio-RoPE reduced type-recovery accuracy to 0.801. These results indicate that ProtSyntax jointly represents acceptor-residue chemistry, motif order and modification identity rather than relying only on residue-level permissibility.

**2.Structure-aware discrimination of PTM microenvironments.** Experimentally validated sites from the homolog-separated test partition were mapped to experimentally determined structures where available and to AlphaFold2 models otherwise. The primary evaluation retained sites whose local structural neighborhoods had a mean pLDDT of at least 70; missing or low-confidence coordinates were masked rather than imputed. For each authentic site, the local microenvironment was defined by residues within 10 Å of the candidate residue and characterized by residue identity, relative backbone-frame orientation, solvent accessibility and the local contact-distance matrix. Three paired decoy classes were generated. Sequence-matched decoys contained the same chemically compatible residue and a highly similar 55-residue motif, as measured by local sequence identity and BLOSUM62 similarity, but differed in solvent exposure or three-dimensional organization. Structure-matched decoys exhibited similar local contact geometry and accessibility but lacked the required motif organization or residue chemistry. Random same-residue decoys were sampled from experimentally unannotated positions in the same protein. Decoys overlapping any known PTM annotation or located within 15 residues of the authentic site were excluded, and sequence and structural similarity thresholds were fixed using the validation partition. Matching sites within the same protein reduced confounding by protein identity, subcellular localization and global fold. ProtSyntax and AstraPTM2 were evaluated without adaptation, and performance was measured using paired-ranking accuracy, AUROC and AUPRC, with 95% confidence intervals obtained by protein-level bootstrap resampling. ProtSyntax achieved a paired-ranking accuracy of 0.887, an AUROC of 0.913 and an AUPRC of 0.786, exceeding AstraPTM2 by 6.2, 6.7 and 7.3 percentage points, respectively. The largest advantage occurred for sequence-matched decoys, for which local motif information was deliberately rendered non-discriminative. Removing Geometric Gated Attention reduced paired-ranking accuracy and AUROC to 0.783 and 0.824, and the sequence-only variant showed a further decline. As structural controls, isotropic Gaussian noise of increasing magnitude was added to the coordinates, and residue frames were separately permuted while preserving the global coordinate distribution. ProtSyntax remained stable under mild coordinate noise but deteriorated after disruption of residue-frame correspondence, demonstrating that its predictions depend on coherent relational geometry rather than nonspecific structural confidence, residue density or the mere availability of a structure.

**3. Transfer to low-resource PTM types.** Cross-modification transfer was evaluated using eight sparsely annotated PTMs—lactylation, formylation, butyrylation, N-palmitoylation, C-linked glycosylation, sulfoxidation, nitration and dephosphorylation—in a leave-one-PTM-type-out design. For each experiment, all positive annotations of the target PTM were removed from model training, and proteins homologous to the support or query proteins were excluded from the remaining corpus using the same 50% sequence-identity threshold. The query set was held fixed across models, whereas protein-disjoint support sets containing 0, 10 or 50 positive sites were constructed. Chemically compatible but unannotated residues were sampled as negatives at a 1:10 positive-to-negative ratio, and five independent support-set draws were evaluated for the 10- and 50-shot settings. To make the zero-shot condition independent of target-class site labels, each PTM class was represented by a physicochemical descriptor containing its compatible acceptor residues, chemical family, elemental or mass shift, charge change and approximate hydrophobicity change. For the held-out PTM, the target query was initialized from this descriptor through the physicochemical embedding used by PACE-Nash, without exposing the model to any positive site from that class. In the few-shot settings, only the class-specific query and lightweight adaptation parameters were updated, with early stopping based on a protein-disjoint validation subset; the shared encoder remained fixed to reduce overfitting. Fine-tuned ESM-C and SaProt linear probes and PTMGPT2 received the same support, validation and query samples. Macro-AUPRC across the eight PTMs was used as the primary metric because of class imbalance, with per-class AUROC and variability across support draws reported as secondary measures. ProtSyntax achieved macro-AUPRC values of 0.432, 0.611 and 0.742 under the 0-, 10- and 50-shot settings, respectively, compared with 0.286/0.421/0.618 for ESM-C, 0.318/0.457/0.630 for SaProt and 0.274/0.402 in the corresponding zero- and 10-shot PTMGPT2 settings. Removing PACE-Nash consistently impaired few-shot transfer. Representation-space analysis further showed significant neighborhood enrichment among chemically related modifications: lysine acylations clustered with acetylation, succinylation and crotonylation, whereas cysteine-centered PTMs formed a separate region, and transfer was weaker between chemically unrelated classes. These findings indicate that ProtSyntax organizes PTM classes according to reusable biochemical constraints, allowing a rare modification to be positioned within an existing chemical manifold rather than learned entirely from its limited annotations.

**4. Reconstruction of combinatorial PTM crosstalk**. The crosstalk branch was trained using 262 positive and 12,468 negative PTM-pair examples assembled from the DeepPCT dataset and complementary curated resources. Each positive record specified a source site, a target site, their PTM identities and an experimentally supported cooperative or antagonistic relationship. Negative examples were constructed from chemically compatible site pairs within the same proteins but lacked reported evidence of functional crosstalk; negatives were matched to positives by sequence separation and PTM-type combination to prevent the model from exploiting trivial distance or label-frequency differences. For independent evaluation, 100 relationships were curated from PTMcode v2, their directions were checked against the associated publications, and records overlapping the training proteins or their homologs were removed. Pairs in which both PTMs competed for the same residue were evaluated separately as mutually exclusive site-level events and were not included in the directional knockout analysis. For each remaining pair, ProtSyntax first calculated the target-site probability from the native sequence. The source residue was then replaced with alanine to remove its modification competence while leaving the remaining sequence and backbone geometry unchanged, and the target probability was recalculated. A reduction in target probability was interpreted as evidence of cooperative crosstalk, whereas an increase indicated antagonism. Effects with an absolute probability change below 0.05, a threshold selected on the validation data, were classified as unresolved and counted as incorrect. To test whether the conclusions depended specifically on alanine substitution, sensitivity analyses used residue-specific nonmodifiable substitutions and masked-site perturbations, and only directionally concordant relationships were regarded as robust. DeepPCT and ProXTalk were retrained on the same partition and evaluated using identical native and perturbed sequences. ProtSyntax recovered the annotated direction for 91% of the independent relationships, compared with 73% for DeepPCT and 70% for ProXTalk; label-permuted and distance-matched null pairs produced near-random directional agreement. In representative cases, removal of a phosphorylation site reduced the associated SUMOylation probability from 0.87 to 0.62, consistent with cooperative regulation, whereas perturbation of a glycosylation site increased phosphorylation probability from 0.69 to 0.91, indicating antagonism. Thus, ProtSyntax encodes PTMs as conditional events within a shared sequence–structure state rather than as independent residue labels. Because residue substitution also changes side-chain chemistry, these perturbation responses should be interpreted as model-derived regulatory hypotheses requiring experimental validation, rather than as direct evidence of biochemical causality.

**5.Coupling of PTM perturbations to enzyme function**. We assembled an independent enzyme-perturbation benchmark by integrating experimentally annotated or high-confidence PTM sites with matched wild-type and variant kinetic measurements from CatPred-DB, BRENDA, SABIO-RK and the corresponding primary literature. CatPred-DB comprised 77,020 kinetic records, including 23,917 kcat, 41,174 Km and 11,929 Ki measurements. Only wild-type–variant pairs measured using the same substrate, compatible assay conditions and convertible units were retained, and all evaluation enzymes and their close homologs were excluded from model training. Catalytic residues and substrate-binding pockets were annotated using UniProt, M-CSA and BioLiP, whereas regulatory and allosteric regions were obtained from UniProt and the Allosteric Database. Sites within 8 Å of a catalytic or ligand-contacting residue were classified as catalytic/pocket-proximal; sites overlapping an annotated regulatory region were classified separately; and solvent-exposed sites more than 20 Å from these regions served as distal controls. For each PTM site, we generated a modification-blocking substitution and, where a well-established proxy existed, a modification-mimetic substitution—for example, Ser/Thr-to-Ala and Ser/Thr-to-Asp/Glu for phosphorylation or Lys-to-Arg and Lys-to-Gln for lysine acylation. Native and perturbed sequences were processed through both the PTM and kinetic heads. Backbone coordinates were held fixed in the primary analysis to isolate the effect of residue chemistry, while a subset was re-evaluated using independently predicted variant structures to confirm robustness to local structural relaxation. The PTM-to-function response was defined as the perturbation-induced change in the kinetic prediction conditioned on the corresponding change in PTM compatibility. Predicted and experimentally measured kinetic changes were calculated in log space and compared using Spearman correlation; absolute changes were additionally analyzed to quantify effect magnitude independently of direction. CatPred was evaluated on the same wild-type–variant pairs and experimental targets. ProtSyntax achieved correlations of 0.628, 0.547 and 0.512 for kcat, Km and Ki, respectively, compared with 0.501, 0.422 and 0.395 for CatPred. Catalytic- or pocket-proximal perturbations produced a mean predicted $\Delta$ log Kcat of 0.61, compared with 0.18 for distal surface sites. Removing kinetic supervision reduced the kcat correlation to 0.421, and removal of PACE-Nash further reduced it to 0.386, supporting the contribution of joint residue- and protein-level optimization. Evidential uncertainty was assessed using prediction-variance quartiles: the accuracy of the predicted kcat-change direction was 0.781 in the high-confidence subset and 0.594 in the low-confidence subset, compared with 0.671 and 0.452 for CatPred. When restricted to PTMs near catalytic pockets, substrate-binding regions or annotated regulatory domains, ProtSyntax achieved a directional accuracy of 0.823, compared with 0.698 for CatPred. Collectively, these analyses establish a quantitative and uncertainty-aware connection between perturbation of local PTM syntax, structural-functional context and experimentally measurable changes in enzyme activity.

##### 4.7.3 Detailed Description of the Experiments in Section 2.1 and 2.3

**1.Zero-shot prediction of PTM-mediated variant effects.** We constructed an independent variant benchmark by integrating pathogenic or likely pathogenic germline missense variants from ClinVar, recurrent or curated somatic cancer mutations from COSMIC and population polymorphisms from gnomAD. ClinVar variants with conflicting clinical interpretations were excluded, and COSMIC variants were retained only when they occurred in Cancer Gene Census proteins or were supported as recurrent cancer-associated events. Benign controls comprised ClinVar benign or likely benign variants and gnomAD variants with no reported disease association and a population allele frequency above 0.1%. Experimentally supported PTM sites were obtained from UniProtKB and PhosphoSitePlus and mapped to the corresponding reference protein sequences. Variants were included when they directly affected a modifiable residue, occurred within 15 residues of a validated PTM site or lay within 10 Å of that site in an experimentally determined or AlphaFold2-predicted structure. To minimize confounding, disease-associated and benign variants were matched by protein where possible and otherwise by PTM type, wild-type residue, sequence and structural distance to the PTM site, local pLDDT and substitution severity measured by the Grantham score. No variant labels, ClinVar/COSMIC annotations or pathogenicity scores were used during ProtSyntax training or threshold selection; the task was therefore zero-shot with respect to variant-effect prediction. For each variant, ProtSyntax was applied to the reference and mutant proteins using identical PTM tasks. Mutant structures were generated with AlphaFold2 using the substituted sequence and subsequently subjected to local side-chain repacking and restrained structural relaxation, while residues outside a 12-Å radius of the mutation were constrained to the reference backbone. Reference and mutant structures were aligned using unaffected Cα atoms, and local coordinate displacement, solvent-accessibility change and residue-contact disruption were recorded. Changes in AlphaFold2 pLDDT were treated only as indicators of model confidence and not as evidence of physical destabilization. For every nearby PTM site jjj, we calculated $\Delta P_{PTM,J} = P_{mut,j} - P_{ref,j}$. A variant-level PTM-disruption score was then defined as the largest distance-weighted absolute change across nearby sites, while the signed site-level changes were retained to distinguish predicted PTM loss from aberrant PTM creation. This score used no conservation, allele-frequency, gene-disease or general pathogenicity features. ProtSyntax discriminated disease-associated variants from matched benign variants with an AUROC of 0.885 and an AUPRC of 0.812, compared with 0.801 and 0.729 for PTMGPT2. Performance was additionally evaluated separately for variants that directly altered the modified residue and variants that preserved the acceptor residue but perturbed its spatial neighborhood. The latter group provided a stringent test of geometric reasoning because residue compatibility and much of the linear motif remained unchanged. Sequence-only, GGA-ablated and residue-frame-permuted controls showed reduced discrimination in this subset, indicating that three-dimensional microenvironment disruption contributed information beyond local sequence alteration. In the representative PTEN example, a sequence-distant but spatially adjacent mutation reduced the predicted ubiquitination probability by 0.47 and increased the GGA-derived spatial incompatibility penalty 3.2-fold, consistent with disruption of the local

E3-ligase-accessible environment. All confidence intervals were estimated by protein-level bootstrap resampling, ensuring that multiple variants from the same protein were not treated as independent observations. These analyses demonstrate how ProtSyntax can translate variant-induced changes in local PTM syntax and geometry into interpretable residue-level regulatory hypotheses, although the predicted mechanisms require biochemical validation.

**2.Modeling PTM-dependent liquid–liquid phase separation.** Experimentally characterized LLPS-associated proteins were collected from LLPSDB, PhaSepDB and PhaSePro and supplemented with primary studies reporting saturation concentrations Csat for unmodified proteins and matched PTM-manipulated conditions. We retained experiments in which the reference and modified protein constructs were measured under matched temperature, pH, ionic strength, crowding-agent and nucleic-acid conditions. When absolute measurements originated from different studies, Csat values were not compared directly; instead, the PTM response was expressed as the within-study change $\Delta log_{10}C_{sat} = loh_{10}(C_{sat,modified}/C_{sat,reference})$. PTM states were supported by site-resolved mass spectrometry, enzyme-mediated modification or validated modification-mimetic constructs. Intrinsically disordered regions were identified using curated DisProt or MobiDB annotations and supplemented with IUPred2A predictions requiring a disorder score above 0.5 across at least 30 consecutive residues. Candidate PTM sites were restricted to the experimentally manipulated IDRs. Because static AlphaFold2 coordinates are unreliable in these regions, residues with pLDDT below 50 were structurally masked, setting their geometric contribution to zero and allowing inference to depend primarily on residue chemistry, motif organization and bidirectional sequence context through Bi-Gated DeltaNet. No LLPS labels or Csa measurements were included in model training. For each protein, ProtSyntax was first evaluated in its unmodified reference state. Experimentally reported PTMs were then introduced sequentially through PTM-state indicators that preserved the underlying amino-acid identity, and the probabilities of the remaining chemically compatible sites were recalculated after each step. The multi-site response score was defined as the change in the length-normalized mean conditional PTM propensity between the reference and imposed modification states; this formulation captured whether accumulating modifications promoted further modification, produced saturation or suppressed additional PTM events. Correlation with $log_{10}C_{sat}$ was evaluated using Spearman's ρ\rhoρ, with significance determined by study-stratified permutation testing and confidence intervals obtained by protein-level bootstrap resampling. ProtSyntax produced a correlation of ρ=−0.71, substantially stronger than PTMGPT2 (ρ=−0.438). The relationship remained evident after removing individual proteins or studies, indicating that it was not driven by a single extensively characterized condensate system. We further examined the low-complexity domain of the RNA-binding protein FUS as a controlled multisite phosphorylation trajectory. Serine and threonine sites were introduced sequentially according to their baseline phosphorylation probabilities, and alternative modification orders were evaluated to assess order dependence. Bi-Gated DeltaNet cross-gating weights were summarized over short-range and long-range residue interactions after each step. As proximal phosphorylation accumulated, the representation shifted away from short-range cohesive coupling toward longer-range, charge-sensitive interactions, consistent with increasing electrostatic repulsion. After the third proximal phosphorylation event, the conditional probability of a subsequent modification decreased from 0.82 to 0.31, whereas MTPrompt-PTM showed only a modest decline from 0.70 to 0.55. Site-order permutation, PTM-label permutation and matched modification of unrelated IDR residues did not reproduce the same response. This threshold-like behavior represents a nonlinear transition in the model's conditional PTM grammar rather than an experimentally established thermodynamic phase boundary. Collectively, the results suggest that ProtSyntax can connect multivalent PTM accumulation within structurally disordered regions to experimentally measured condensate behavior, while generating testable hypotheses concerning modification order, cooperativity and saturation.

###### 4.7.4 Interpretability Analysis of the Experiments in Section 2.1

The five validation tasks were designed to interrogate complementary levels of PTM language, ranging from residue-level completion to structural permissiveness, cross-modification transfer, combinatorial regulation and protein-level functional semantics. The performance of ProtSyntax should therefore not be attributed to a single architectural component. Rather, it emerges from the coordinated inductive biases of Bio-RoPE, Bi-Gated DeltaNet, Geometric Gated Attention (GGA), the sparse Mixture-of-Experts (MoE) backbone and PACE-Nash optimization. Their task-specific contributions can be interpreted as follows.

**1)** In the cloze-style PTM-type and PTM-site recovery experiments, ProtSyntax benefits from representing a modification event as a contextually constrained biochemical token rather than as an independent residue label. Bio-RoPE is particularly important because it makes positional encoding conditional on both motif order and residue physicochemistry. Consequently, residues occupying equivalent relative positions do not receive equivalent representations when their chemical properties differ, allowing the model to distinguish a well-formed PTM motif from a compositionally similar but order-disrupted or chemically incompatible sequence. Bi-Gated DeltaNet further integrates evidence from both N- and C-terminal contexts around the candidate site. Its cross-gating mechanism reinforces motifs supported by coherent bilateral context while suppressing one-sided or incidental motif matches, which is especially important when several chemically compatible residues occur within the same segment. GGA provides an additional constraint by distinguishing residues with similar linear contexts but different structural accessibility or spatial neighborhoods. Meanwhile, the sparse MoE architecture supplies sufficient conditional capacity for partially distinct residue chemistries and PTM classes without requiring them to be represented by a single homogeneous transformation. PACE-Nash further regularizes the shared representation through physicochemical and contrastive objectives, increasing the separation between biologically compatible contexts and counterfactual perturbations. The resulting PTM score therefore reflects the contextual well-formedness of a modification event, explaining both the improved cloze recovery and the stronger sensitivity to motif shuffling and residue-rule violation.

**2)** In the structure-dependent microenvironment decoy experiment, the principal advantage arises from GGA, because sequence-matched decoys were deliberately constructed to remove the shortcut of local motif recognition. GGA introduces residue-frame-based geometric relationships directly into the attention computation, enabling a candidate site to be evaluated according to the spatial arrangement, orientation and accessibility of its surrounding residues. A validated site can therefore receive convergent support from residues that are distant in primary sequence but form a permissive three-dimensional microenvironment, whereas a sequence-matched decoy is penalized when the corresponding spatial configuration is absent. Bio-RoPE and Bi-Gated DeltaNet remain complementary rather than redundant: the former verifies the chemical and positional validity of the local motif, whereas the latter supplies domain-scale sequence context that can distinguish a functional region from an incidental surface motif. The interleaving of sequence-dominant Bi-Gated DeltaNet blocks with periodic geometry-constrained GGA blocks allows structural evidence to refine, rather than replace, sequence semantics. Moreover, masking unreliable coordinates and gating geometric contributions prevent uncertain structures from dominating the representation, allowing the model to revert toward sequence-based reasoning when structural confidence is low. This architecture explains why ProtSyntax is robust to mild coordinate noise yet sensitive to disruption of residue-frame correspondence: the model has learned relational geometry associated with PTM permissiveness rather than merely exploiting generic structural descriptors.

**3)** The low-resource transfer results indicate that ProtSyntax decomposes PTM classes into reusable biochemical primitives rather than memorizing modification-specific labels. These primitives include acceptor-residue identity, side-chain reactivity, motif organization, steric accessibility and the broader sequence–structure context in which a chemical group can be installed. Bio-RoPE embeds residue chemistry into the positional representation, making chemically related modifications more likely to share compatible local syntax. The physicochemical-aware contrastive component of PACE-Nash further encourages modifications with related chemical mechanisms to occupy neighboring regions of the latent space while maintaining separation from incompatible classes. Nash-based objective coordination reduces destructive competition between abundant PTM classes, low-resource classes and protein-level functional supervision, thereby preventing the representation from being dominated exclusively by high-frequency labels. The sparse MoE backbone adds conditional capacity for both shared and modification-biased transformations: related PTMs can reuse common representational pathways, whereas specialized experts can capture distinctions in substrate chemistry or recognition context. Bi-Gated DeltaNet and GGA additionally provide PTM-label-independent information about long-range context and structural permissiveness. Consequently, when a rare PTM is introduced with only a few examples, adaptation mainly requires learning how the new label maps onto an already organized biochemical manifold rather than learning the underlying modification rules de novo. This interpretation is consistent with the strong transfer among lysine acylations and cysteine-centered modifications, together with weaker transfer between chemically unrelated PTM classes.

**4)** ProtSyntax reconstructs PTM crosstalk more effectively because each modification is encoded as a conditional event embedded within a shared sequence–structure state, rather than as an output that is statistically independent of other PTM predictions. Substitution of one regulatory residue therefore changes not only its local PTM score but also the contextual representation available to other candidate sites. Bi-Gated DeltaNet propagates this perturbation bidirectionally along the protein sequence, allowing changes at one site to influence distal residues through shared motifs, domain organization or regulatory regions. GGA provides a complementary spatial propagation route, particularly when interacting PTM sites are distant in sequence but adjacent within the folded structure. Bio-RoPE ensures that the perturbation is interpreted in terms of altered residue chemistry and motif order rather than as an arbitrary token replacement. The sparse MoE backbone is well suited to the heterogeneity of crosstalk logic, because cooperative, antagonistic and context-dependent relationships may require distinct nonlinear transformations rather than a single universal interaction rule. Finally, direct crosstalk supervision, coordinated by PACE-Nash with general PTM and kinetic objectives, encourages these perturbation responses to remain compatible with both residue-level modification syntax and protein-level functional state. Thus, the change in the probability of a second PTM after perturbing the first reflects an internally learned conditional grammar. These in silico responses should nevertheless be interpreted as model-inferred regulatory dependencies rather than direct evidence of biochemical causality, because amino-acid substitution simultaneously removes modification competence and alters the underlying side-chain chemistry.

**5)** In the PTM-to-function coupling experiment, the decisive feature of ProtSyntax is the shared encoder jointly trained on residue-level PTM prediction and full-length enzyme kinetic regression. This design forces local modification representations to remain informative about the global biochemical state of the protein. When a PTM site is perturbed, Bi-Gated DeltaNet can propagate the resulting representational change across the sequence toward catalytic motifs, substrate-recognition regions and distal regulatory domains. GGA simultaneously evaluates whether the perturbed site is spatially coupled to a catalytic pocket, binding interface or putative allosteric network, including interactions between residues that are widely separated in sequence. These local and non-local effects are integrated into the full-length protein representation and translated by the kinetic heads into changes in $K_{\text{cat}}$, $K_m$ and $K_i$. PACE-Nash is particularly important in this setting because residue classification and kinetic regression impose objectives at different biological scales and can otherwise generate conflicting gradients. Adaptive Nash coordination preserves a representation that is simultaneously sensitive to local PTM compatibility and global functional variation, while evidential regression associates kinetic predictions with estimates of predictive uncertainty. The sparse MoE backbone further accommodates heterogeneity among enzyme families, catalytic mechanisms and kinetic parameters by providing conditional computational pathways. The larger predicted effects for PTMs near catalytic or substrate-binding regions, together with the deterioration observed after removing kinetic supervision or PACE-Nash, therefore support a geometry- and context-dependent syntax-to-function mapping rather than a nonspecific association between mutation magnitude and kinetic change. Unlike a conventional kinetic predictor, ProtSyntax contains an explicit residue-level regulatory coordinate through which the functional consequences of individual modification events can be interrogated.

##### 4.8 Interpretable local–global reasoning in ProtSyntax

ProtSyntax enables interpretable PTM modeling by integrating residue-level syntax with protein-scale contextual constraint. The residue-centered PTM input first directs the encoder toward local determinants, including flanking motifs, amino-acid chemistry and short-range physicochemical patterns. These local cues are further contextualized through a shared multi-task encoder trained with full-length enzyme kinetic regression, which encourages retention of domain organization, catalytic and substrate-binding regions, and distal regulatory signals. As shown in Figure 5, Bi-Gated DeltaNet captures this local–global coupling by bidirectionally propagating N- and C-terminal sequence evidence and using cross-gating to recalibrate context at each residue, enabling the model to distinguish coherent PTM-associated patterns from incidental local motifs. Geometric Gated Attention complements this mechanism by imposing explicit three-dimensional constraints on residue interactions, suppressing semantically plausible but spatially inconsistent associations while emphasizing contacts supported by both sequence context and structural proximity. Through the 3:1 interleaving of Bi-Gated DeltaNet and Geometric Gated Attention, ProtSyntax therefore interprets PTM susceptibility as an emergent property jointly shaped by local residue syntax, distal sequence context and the global structural–functional state of the protein.

##### 4.9 Instructions for Using ProtSyntax Lab

ProtSyntax was used as a computational decision-support platform for the identification, prioritization, and interpretation of candidate post-translational modification sites from protein sequence data (ProtSyntax Lab, Figure S5). For each analysis, the input protein sequence was provided in standard one-letter amino-acid notation or uploaded in FASTA format, and the corresponding PTM prediction task was selected according to the biological question being addressed. When the modification chemistry or residue specificity was known in advance, residue-restricted tasks were used to focus the prediction space; for exploratory analyses, broader PTM screening was applied to identify potentially relevant candidate sites. Prior to inference, the input sequence was checked for length, valid amino-acid characters, candidate residue coverage, and local sequence composition, ensuring that downstream predictions were generated from model-compatible sequence data.

The model output was interpreted as a ranked set of computational predictions rather than direct experimental evidence. Each candidate site was evaluated using its residue position, predicted modification type, local motif window, calibrated probability score, confidence category, and distribution along the protein sequence. For computational analyses, these outputs provide structured features that can be integrated with downstream workflows such as comparative sequence analysis, structural modeling, variant effect assessment, proteome-scale screening, or machine-learning-based prioritization. For biological interpretation, predicted sites should be considered together with orthogonal evidence, including evolutionary conservation, domain architecture, solvent accessibility, intrinsically disordered regions, known functional motifs, available proteomic observations, and experimental validation when available. Confidence thresholds were selected according to the objective of the study. More stringent thresholds were used when prioritizing sites for validation or mechanistic follow-up, whereas lower thresholds were used in exploratory settings where sensitivity and candidate discovery were favored. The resulting predictions were therefore treated as hypothesis-generating evidence to guide experimental design, database comparison, or computational prioritization, rather than as definitive annotations. To ensure reproducibility, each prediction run recorded the input sequence identifier, organism context, selected PTM task, confidence threshold, maximum number of reported sites, ProtSyntax model version, inference date, and relevant model output fields.

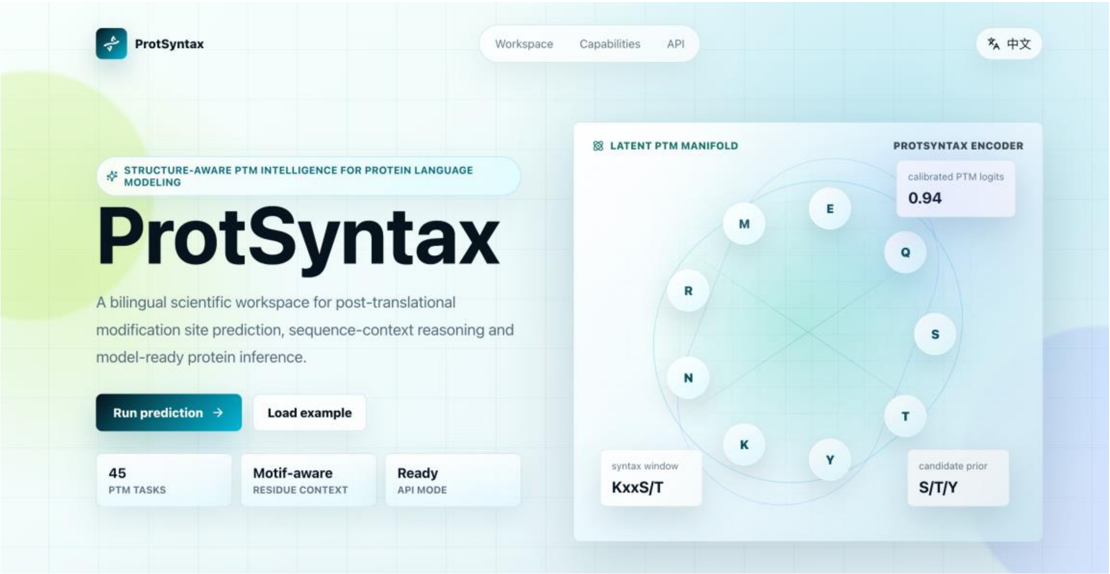


Figure S5. Schematic illustration of ProtSyntax Lab. This figure complements the schematic presented in the main text.

#### 5. Hyperparameters

The detailed hyperparameter table for ProtSyntax is provided below; please refer to Table S6. In addition, ProtSyntax has been integrated into open-source large-model frameworks, including Hugging Face, Ollama and vLLM, enabling researchers to readily access and deploy the model.

Table S6. The detailed hyperparameters of ProtSyntax

| Module | Hyperparameter | Value |
|---|---|---|
| Global Architecture | Number of Layers | 33 |
| | Block Interleaving Ratio | 3:1 |
| | Hidden Dimension | 1280 |
| | Number of Bi-Gated DeltaNet | 25 |
| | Number of GGA Blocks | 8 |
| | Activation Function | SwiGLU |
| | Residual Dropout | 0.1 |
| Input Configuration | PTM Window Length | 55 |
| | Maximum Sequence Length | 1024 |
| | Sliding-window Stride | 512 |
| | Dynamic Padding | True |
| | Structure-missing Mask | True |
| Sparse MoE | Number of Routed Experts | 16 |
| | Experts Activated per Token | 2 |
| | MoE Layer Frequency | Every |
| | Router Scoring Function | Softmax |
| | Load-balancing Loss Coefficient | 0.01 |
| | Token Dropping | False |
| Bio-RoPE | Structural Channels | 20% |
| | Physicochemical Channels | 30% |
| | Rotary Base Frequency | 10000.0 |
| Bi-Gated DeltaNet | Number of Heads | 16 |
| | Single Head Dimension | 80 |
| | RMS Norm Epsilon | 1e-6 |
| Geometric Gated Attention | Geometric Probes | 4 |
| | Geometric Scaling Init | 0.0 |
| | Attention Dropout | 0.1 |
| PACE-Nash Optimization | Contrastive Temperature | 0.7 |
| | Uncertainty Variance Reg | 0.01 |
| | Nash Update Frequency | 10 |
| | Nash Inner LR | 10 |
| | Initial Task Weights | 3*1/3 |
| | Probability Clipping | 0.05 |
| | Jaccard Similarity Threshold | 0.1 |
| | Nash Coefficient Learning Rate | 0.01 |
| Train&Optimization | Batch Size | 256/512 |

| | |
|---|---|
| Peak Learning Rate | 4e-4 |
| Warmup steps | 5000 |
| Optimizer | AdamW |
| Peak Learning Rate | 4e-4 |
| Minimum Learning Rate | 4e-5 |
| Learning-rate Scheduler | Cosine decay |
| Weight Decay | 0.1 |
| Numerical Precision | BF16 |
| Gradient Checkpointing | True |
| Validation Frequency | Every 500 Steps |

#### 6. Detailed performance metrics of the comparative models

Tables S7–S14 supplement Table 1 in the main text by reporting the detailed performance metrics of the remaining comparative models. To preserve the readability and continuity of the main supplementary text, these three tables are presented in the final section.

ProtSyntax is capable of predicting 40 distinct post-translational modification (PTM) types, making it the most comprehensive protein large language model (PLM) to date in terms of simultaneous PTM prediction capacity. The datasets used in the comparative experiments are described in Section 2 of the Supplementary Material. For the PTM site prediction task, we deliberately benchmarked ProtSyntax against universal prediction models capable of multi-PTM site identification. Among the 40 evaluated PTMs, 13 are supported by more than two baseline models, while the remaining 27 are exclusively predictable by ProtSyntax and AstraPTM2. The head-to-head comparison between ProtSyntax and AstraPTM2 for the other 27 PTM types is also available in Table S7. Detailed comparative performance for the 13 widely supported PTM types is provided in Table S8, supplementing the overarching results in Table S8. Notably, because S-sulfhydration and S-carboxyethylation are rare PTM types uniquely predicted by ProtSyntax, we leveraged the inherent fine-tunability of PTM-Mamba and fine-tuned it specifically to predict these two modifications to establish a comparative baseline.

Table S7. Detailed performance comparison for the PTM prediction task. MCC1 represents the MCC achieved by ProtSyntax, whereas MCC2 represents the MCC achieved by the second-best model. Sub_Model indicates the corresponding second-best comparative model. The same notation applies to AP.

| **PTM Type** | **MCC1** | **MCC2** | **AP1** | **AP2** | **Sub_Model** |
|---|---|---|---|---|---|
| Phosphorylation | 0.893 | 0.770 | 0.937 | 0.848 | PTMGPT2 |

| | | | | | |
|---|---|---|---|---|---|
| Acetylation | 0.872 | 0.781 | 0.914 | 0.812 | PTM-Mamba |
| Ubiquitination | 0.845 | 0.749 | 0.873 | 0.794 | PTMGPT2 |
| N-linked glycosylation | 0.891 | 0.749 | 0.919 | 0.805 | AstraPTM2 |
| O-linked glycosylation | 0.904 | 0.806 | 0.933 | 0.803 | MTPrompt-PTM |
| Methylation | 0.823 | 0.712 | 0.845 | 0.748 | MTPrompt-PTM |
| Sumoylation | 0.856 | 0.719 | 0.892 | 0.810 | PTM-Mamba |
| S-palmitoylation | 0.809 | 0.660 | 0.851 | 0.742 | PTMGPT2 |
| Disulfide bond | 0.768 | 0.659 | 0.841 | 0.730 | AstraPTM2 |
| S-Sulfhydration | 0.838 | 0.712 | 0.884 | 0.794 | PTM-Mamba |
| S-Carboxyethylation | 0.887 | 0.773 | 0.908 | 0.822 | PTM-Mamba |
| ADP-ribosylation | 0.860 | 0.729 | 0.899 | 0.778 | AstraPTM2 |
| Neddylation | 0.821 | 0.674 | 0.884 | 0.778 | AstraPTM2 |
| Succinylation | 0.884 | 0.778 | 0.895 | 0.753 | MTPrompt-PTM |
| Crotonylation | 0.915 | 0.813 | 0.972 | 0.836 | AstraPTM2 |
| Myristoylation | 0.852 | 0.728 | 0.888 | 0.808 | AstraPTM2 |
| Farnesylation | 0.942 | 0.830 | 0.966 | 0.819 | AstraPTM2 |
| Geranylgeranylation | 0.908 | 0.767 | 0.912 | 0.793 | AstraPTM2 |
| S-nitrosylation | 0.923 | 0.791 | 0.951 | 0.826 | PTMGPT2 |
| Glutathionylation | 0.958 | 0.832 | 0.972 | 0.858 | PTMGPT2 |
| Hydroxylation | 0.855 | 0.765 | 0.855 | 0.743 | AstraPTM2 |
| Oxidation | 0.934 | 0.840 | 0.974 | 0.845 | AstraPTM2 |
| Deamidation | 0.849 | 0.731 | 0.880 | 0.785 | AstraPTM2 |
| Sulfation | 0.911 | 0.762 | 0.935 | 0.797 | AstraPTM2 |
| Citrullination | 0.923 | 0.783 | 0.968 | 0.887 | AstraPTM2 |
| Amidation | 0.949 | 0.835 | 0.955 | 0.871 | PTMGPT2 |
| GPI anchor | 0.938 | 0.818 | 0.942 | 0.833 | AstraPTM2 |

| | | | | | |
|---|---|---|---|---|---|
| Malonylation | 0.905 | 0.774 | 0.934 | 0.808 | PTMGPT2 |
| Glutarylation | 0.946 | 0.794 | 0.964 | 0.862 | AstraPTM2 |
| Lactylation | 0.919 | 0.791 | 0.957 | 0.871 | AstraPTM2 |
| Formylation | 0.955 | 0.813 | 0.979 | 0.861 | AstraPTM2 |
| Butyrylation | 0.975 | 0.844 | 0.965 | 0.858 | AstraPTM2 |
| N-palmitoylation | 0.954 | 0.799 | 0.961 | 0.849 | MTPrompt-PTM |
| S-diacylglycerol | 0.762 | 0.611 | 0.825 | 0.733 | AstraPTM2 |
| C-linked glycosylation | 0.837 | 0.692 | 0.872 | 0.788 | AstraPTM2 |
| Gamma-carboxyglutamic acid | 0.739 | 0.590 | 0.787 | 0.693 | AstraPTM2 |
| Sulfoxidation | 0.917 | 0.759 | 0.940 | 0.833 | AstraPTM2 |
| Nitration | 0.834 | 0.687 | 0.877 | 0.781 | AstraPTM2 |
| Pyrrolidone-carboxylic acid | 0.801 | 0.666 | 0.832 | 0.732 | AstraPTM2 |
| Dephosphorylation | 0.759 | 0.662 | 0.801 | 0.683 | AstraPTM2 |

Table S8. Comparative performance of ProtSyntax across the 13 PTM types evaluated against multiple baseline models.

| **Type** | **Model** | **MCC** | **AP** |
|---|---|---|---|
| Phosphorylation | ProtSyntax | 0.893 | 0.937 |
| | PTMGPT2 | 0.770 | 0.848 |
| | AstraPTM2 | 0.751 | 0.805 |
| Acetylation | ProtSyntax | 0.872 | 0.914 |
| | PTM-Mamba | 0.781 | 0.812 |
| | AstraPTM2 | 0.749 | 0.804 |
| Ubiquitination | ProtSyntax | 0.845 | 0.873 |
| | PTMGPT2 | 0.749 | 0.794 |
| | MTPrompt-PTM | 0.744 | 0.799 |
| N-linked glycosylation | ProtSyntax | 0.891 | 0.919 |
| | AstraPTM2 | 0.749 | 0.805 |
| | MTPrompt-PTM | 0.706 | 0.787 |
| O-linked glycosylation | ProtSyntax | 0.904 | 0.933 |
| | MTPrompt-PTM | 0.806 | 0.803 |
| | PTMGPT2 | 0.763 | 0.810 |
| Methylation | ProtSyntax | 0.823 | 0.845 |
| | MTPrompt-PTM | 0.712 | 0.748 |
| | AstraPTM2 | 0.709 | 0.724 |

| | | | |
|---|---|---|---|
| Sumoylation | ProtSyntax | 0.856 | 0.892 |
| | PTM-Mamba | 0.719 | 0.810 |
| | AstraPTM2 | 0.680 | 0.793 |
| S-palmitoylation | ProtSyntax | 0.809 | 0.851 |
| | PTMGPT2 | 0.660 | 0.742 |
| | AstraPTM2 | 0.591 | 0.675 |
| Succinylation | ProtSyntax | 0.884 | 0.895 |
| | MTPrompt-PTM | 0.778 | 0.753 |
| | AstraPTM2 | 0.725 | 0.688 |
| S-nitrosylation | ProtSyntax | 0.923 | 0.951 |
| | PTMGPT2 | 0.791 | 0.826 |
| | AstraPTM2 | 0.755 | 0.803 |
| Glutathionylation | ProtSyntax | 0.958 | 0.972 |
| | PTMGPT2 | 0.832 | 0.858 |
| | AstraPTM2 | 0.694 | 0.721 |
| Amidation | ProtSyntax | 0.949 | 0.955 |
| | PTMGPT2 | 0.835 | 0.871 |
| | AstraPTM2 | 0.788 | 0.826 |
| Malonylation | ProtSyntax | 0.905 | 0.934 |
| | PTMGPT2 | 0.774 | 0.808 |
| | AstraPTM2 | 0.690 | 0.724 |

In Table S9, we report the performance of ProtSyntax and the second-best model, DeepPCT, on the protein post-translational modification crosstalk prediction task. In Table S10, we further provide a comprehensive comparison between ProtSyntax and other baseline models on the same task.

Table S9. Detailed performance comparison for the PTM CrossTalk task. MCC1 represents the MCC achieved by ProtSyntax, whereas MCC2 represents the MCC achieved by the second-best model. Sub_Model indicates the corresponding second-best comparative model. The same notation applies to AUC and AP.

| **Type** | **MCC1** | **MCC2** | **AUC1** | **AUC2** | **AP1** | **AP2** | **Sub_model** |
|---|---|---|---|---|---|---|---|
| PCT | 0.463 | 0.389 | 0.840 | 0.769 | 0.522 | 0.454 | DeepPCT |

Table S10. Performance comparison between ProtSyntax and other models on the PCT task.

| **Model** | **MCC** | **AUC** | **AP** |
|---|---|---|---|
| ProtSyntax | 0.463 | 0.840 | 0.522 |
| DeepPCT | 0.389 | 0.769 | 0.454 |
| ProXTalk | 0.357 | 0.752 | 0.404 |

Tables S10 and S11 provide a detailed comparison between ProtSyntax and existing state-of-the-art models for kinase-specific phosphorylation site prediction. For benchmark selection, we included two recently published leading methods, DCPPS and

LMPhosSite, as representative comparators.

Table S11.Detailed performance comparison for the kinase-specific phosphorylation prediction task. MCC1 represents the MCC achieved by ProtSyntax, whereas MCC2 represents the MCC achieved by the second-best model. Sub_Model indicates the corresponding second-best comparative model. The same notation applies to AUC and AP.

| **Type** | **MCC1** | **MCC2** | **AUC1** | **AUC2** | **AP1** | **AP2** | **Sub_Model** |
|---|---|---|---|---|---|---|---|
| CDK | 0.712 | 0.620 | 0.842 | 0.758 | 0.509 | 0.423 | DCPPS |
| AGC | 0.802 | 0.699 | 0.905 | 0.818 | 0.847 | 0.734 | |
| PKC | 0.773 | 0.697 | 0.869 | 0.778 | 0.815 | 0.716 | |
| MAPL | 0.779 | 0.680 | 0.943 | 0.838 | 0.762 | 0.672 | |

Table S12. Comparative performance of ProtSyntax and existing state-of-the-art models on the kinase-specific phosphorylation site prediction task.

| **Type** | **Model** | **MCC** | **AUC** | **AP** |
|---|---|---|---|---|
| CDK | ProtSyntax | 0.712 | 0.842 | 0.509 |
| | DCPPS | 0.620 | 0.758 | 0.423 |
| | LMPhosSite | 0.569 | 0.714 | 0.385 |
| AGC | ProtSyntax | 0.802 | 0.905 | 0.847 |
| | DCPPS | 0.699 | 0.818 | 0.734 |
| | LMPhosSite | 0.656 | 0.762 | 0.689 |
| PKC | ProtSyntax | 0.773 | 0.869 | 0.815 |
| | DCPPS | 0.697 | 0.778 | 0.716 |
| | LMPhosSite | 0.667 | 0.741 | 0.682 |
| MAPL | ProtSyntax | 0.779 | 0.943 | 0.762 |
| | DCPPS | 0.680 | 0.838 | 0.672 |
| | LMPhosSite | 0.641 | 0.800 | 0.633 |

In Tables S13 and S14, we systematically benchmark ProtSyntax against the state-of-the-art enzyme kinetics prediction models CatPred and DKEP for the prediction of enzyme kinetic parameters.

Table S13. Detailed performance comparison for the enzyme kinetic parameter prediction task. $R^2$1 represents the $R^2$ achieved by ProtSyntax, whereas $R^2$2 represents the $R^2$ achieved by the second-best model. Sub_Model indicates the corresponding second-best comparative model. The same notation applies to MAE.

| **Type** | **$R^2$1** | **$R^2$2** | **MAE1** | **MAE2** | **Sub_Model** |
|---|---|---|---|---|---|
| $K_{cat}$ | 0.670 | 0.596 | 0.593 | 0.697 | CatPred |
| $K_m$ | 0.701 | 0.611 | 0.564 | 0.665 | |

| | | | | |
|---|---|---|---|---|
| $K_i$ | 0.653 | 0.575 | 0.582 | 0.698 |

Table S14. Comparative performance of ProtSyntax in enzyme kinetic parameter prediction.

| **Type** | **Model** | **$R^2$** | **MAE** |
|---|---|---|---|
| $K_{cat}$ | ProtSyntax | 0.670 | 0.593 |
| | CatPred | 0.596 | 0.697 |
| | DKEP | 0.566 | 0.702 |
| $K_m$ | ProtSyntax | 0.701 | 0.564 |
| | CatPred | 0.611 | 0.665 |
| | DKEP | 0.552 | 0.698 |
| $K_i$ | ProtSyntax | 0.653 | 0.582 |
| | CatPred | 0.575 | 0.698 |
| | DKEP | 0.560 | 0.673 |

### 7. Extended Discussion

ProtSyntax establishes a PTM-centered protein large language model for decoding post-translational modification syntax and its functional consequences. Rather than treating PTM prediction as an isolated residue-level annotation problem, ProtSyntax reframes modification biology as a structured protein language problem in which residue chemistry, motif order, long-range sequence context, three-dimensional microenvironment, combinatorial modification state and protein-level function jointly determine regulatory meaning. This formulation is important because PTMs rarely operate as independent biochemical events. A residue may be chemically compatible with a modification but remain unmodified because it is structurally inaccessible, embedded in an unfavorable domain context, or conditionally regulated by other proximal or distal PTMs. Conversely, a local modification can propagate beyond the modified residue to reshape catalytic efficiency, substrate recognition, interaction surfaces, phase behavior or disease-associated regulatory states. By explicitly modeling these hierarchical dependencies, ProtSyntax moves PTM prediction from static site detection toward syntax-aware reasoning over the modified proteome.

A central contribution of ProtSyntax is that it connects architectural design with the biological organization of PTM regulation. Bio-RoPE introduces PTM-aware positional and physicochemical encoding, enabling the model to represent motif order together with residue-specific chemical constraints. Bi-Gated DeltaNet provides efficient bidirectional propagation of long-range sequence evidence, allowing candidate residues to be evaluated in the context of both upstream and downstream regulatory signals rather than through a one-sided or local sequence window alone. Geometric Gated Attention further incorporates residue-frame-based structural constraints directly into the attention mechanism, enabling the model to distinguish residues that are merely sequence-compatible from those located in permissive three-dimensional PTM microenvironments. Finally, PACE-Nash couples residue-level PTM learning with uncertainty-aware enzyme kinetic supervision, encouraging local modification representations to remain aligned with protein-level functional readouts. The resulting model is therefore not simply a larger PTM predictor, but a biologically structured language model in which sequence syntax, structural grammar and functional semantics are jointly optimized. This design yielded consistent improvements across general PTM-site prediction, kinase-specific phosphorylation, PTM crosstalk prediction and enzyme kinetic regression. However, the more important implication is not only higher benchmark performance, but the emergence of capabilities that are difficult to obtain from task-specific classifiers. ProtSyntax recovered masked PTM types and sites in cloze-style evaluations, rejected sequence-matched but structurally invalid decoys, transferred PTM rules to low-resource modification types with few examples, reconstructed cooperative and antagonistic crosstalk relationships, and linked in silico PTM perturbations to quantitative changes in Kcat, Km and Ki. These results suggest that the model learns transferable PTM syntax rather than memorizing modification-specific patterns. In particular, its ability to generalize to rare PTMs and combinatorial modification states supports the view that PTM regulation contains reusable biochemical grammar, including residue compatibility, motif organization, chemical relatedness among modification classes and context-dependent dependency rules.

ProtSyntax also provides a step toward function-aware interpretation of the modified proteome. Most existing PTM predictors answer whether a residue is likely to be modified, but they do not directly address why the modification matters. By integrating enzyme kinetic regression and perturbation-based analysis, ProtSyntax begins to bridge this gap. The model can prioritize PTM events according to their predicted functional effects, distinguish perturbations near catalytic or substrate-binding regions from distal solvent-exposed sites, and estimate the directionality and confidence of functional changes. This capability is particularly relevant for experimental prioritization, protein engineering and disease variant interpretation, where the key question is often not simply whether a modification exists, but whether it alters protein activity, molecular recognition or regulatory state. In this sense, ProtSyntax offers a computational framework for moving from PTM cataloguing toward mechanistic hypothesis generation.

The generalization analyses further indicate that PTM syntax can extend beyond canonical site annotation. In disease-associated variant analysis, ProtSyntax translated local steric or electrostatic disruption of PTM microenvironments into interpretable regulatory consequences, including loss or gain of modification propensity. In condensate-associated proteins, the model captured nonlinear responses of multivalent PTM states in intrinsically disordered regions, suggesting that learned sequence syntax can remain informative even when high-confidence tertiary structure is unavailable. These results broaden the conceptual scope of PTM modeling: PTMs can be viewed not only as residue-specific chemical marks, but also as dynamic regulatory operators that reshape protein interaction landscapes, phase behavior and disease-relevant functional states. ProtSyntax provides an initial computational route for studying these effects in a unified sequence–structure–function framework.

Despite these advances, several limitations remain. First, ProtSyntax is constrained by the incompleteness, imbalance and context bias of current PTM annotations. Many rare, transient, low-stoichiometry or condition-specific modifications remain underrepresented, and negative samples may include unobserved true PTM sites due to incomplete experimental coverage. Second, although Geometric Gated Attention enables structure-aware reasoning, the structural branch still depends on predicted or static conformations. Such structures cannot fully capture conformational ensembles, enzyme–substrate encounter complexes, membrane-associated states, allosteric transitions or condensate-specific molecular environments. Third, the current model links PTM syntax to function primarily through enzyme kinetic supervision and in silico perturbation. It does not yet explicitly model PTM occupancy, temporal signaling dynamics, proteoform distributions, enzyme availability, subcellular localization or cell-type-specific regulatory programs. These factors are essential for determining when and where a predicted PTM event is realized in vivo. Future work should therefore extend ProtSyntax along three directions. The first is data expansion: integrating quantitative and time-resolved proteomics, perturbation screens, kinase and writer–eraser specificity maps, interactome data and cell-state-resolved PTM atlases would allow the model to learn not only modification compatibility, but also modification regulation under biological conditions. The second is dynamic structural modeling: incorporating conformational ensembles, molecular dynamics, protein–protein docking, enzyme–substrate complexes and intrinsically disordered-region representations could improve the modeling of PTMs governed by transient or context-dependent structural states. The third is functional grounding: coupling PTM syntax with phenotypic readouts, disease annotations, pathway perturbations and experimentally measured proteoform effects would help transform PTM prediction into causal regulatory inference.

More broadly, ProtSyntax supports a shift in PTM computational biology from the question “Can this residue be modified?” to a richer set of questions: “Under which molecular context is it modified, how does it interact with other modifications, and what functional consequence does it produce?" By framing PTM biology as a foundation-model problem over protein sequence, structure and function, ProtSyntax provides a scalable and interpretable framework for decoding the regulatory syntax of the modified proteome. This framework can assist in prioritizing functional PTM events, interpreting regulatory variants, designing protein perturbation experiments and generating mechanistic hypotheses for disease biology and protein engineering.

#### Reference


[1] Miller, M. L., & Blom, N. (2009). Kinase-specific prediction of protein phosphorylation sites. *Phospho-Proteomics: Methods and Protocols*, 299-310.

[2] Chen, M., Zhang, W., Gou, Y., Xu, D., Wei, Y., Liu, D., ... & Xue, Y. (2023). GPS 6.0: an updated server for prediction of kinase-specific phosphorylation sites in proteins. *Nucleic acids research*, *51*(W1), W243-W250.

[3] Wang, D., Liu, D., Yuchi, J., He, F., Jiang, Y., Cai, S., ... & Xu, D. (2020). MusiteDeep: a deep-learning based webserver for protein post-translational modification site prediction and visualization. *Nucleic Acids Research*, *48*(W1), W140-W146.

[4] Zhai, J., Wang, Z., Tang, C., Zhong, H., Xu, Z., Liu, Y., ... & Lu, T. (2025). A general language model for peptide function identification. *arXiv preprint arXiv:2502.15610*.

[5] Han, Y., He, F., Shao, Q., Wang, D., & Xu, D. (2025). MTPrompt-PTM: A Multi-Task Method for Post-Translational Modification Prediction Using Prompt Tuning on a Structure-Aware Protein Language Model. *Biomolecules*, *15*(6), 843.

[6] Shrestha, P., Kandel, J., Tayara, H., & Chong, K. T. (2024). Post-translational modification prediction via prompt-based fine-tuning of a GPT-2 model. *Nature Communications*, *15*(1), 6699.

[7] Peng, Z. (2024, October). PTM-Mamba: a PTM-aware protein language model with bidirectional gated Mamba blocks. In *Proceedings of the 33rd ACM international conference on information and knowledge management* (pp. 5475-5478).

[8] Bozkurt, Ç., Vasilyeva, A., & Goteti, A. (2025). AstraPTM2: A Context-Aware Transformer for Broad-Spectrum PTM Prediction. *bioRxiv*, 2025-10.

[9] Sun, S., Wong, T. S., Zhang, X. Q., Pu, J. K., Lee, N. P., Day, P. J., ... & Leung, G. K. (2012). Protein alterations associated with temozolomide resistance in subclones of human glioblastoma cell lines. *Journal of neuro-oncology*, *107*(1), 89-100.

[10] Ou, S., Song, W., Li, J., Gao, S., Ma, Y., & Shi, X. (2025, December). ProXTalk:

A pLLM-Driven Dual-Stream Framework for Inter-Protein PTM Crosstalk Prediction. In *2025 IEEE International Conference on Bioinformatics and Biomedicine (BIBM)* (pp. 339-344). IEEE.

[11] Rives, A., Meier, J., Sercu, T., Goyal, S., Lin, Z., Liu, J., ... & Fergus, R. (2021). Biological structure and function emerge from scaling unsupervised learning to 250 million protein sequences. *Proceedings of the national academy of sciences*, *118*(15), e2016239118.

[12] Su, M. G., & Lee, T. Y. (2013). Incorporating substrate sequence motifs and spatial amino acid composition to identify kinase-specific phosphorylation sites on protein three-dimensional structures. *BMC bioinformatics*, *14*(Suppl 16), S2.

[13] Zhou, Z., Zhang, L., Yu, Y., Wu, B., Li, M., Hong, L., & Tan, P. (2024). Enhancing efficiency of protein language models with minimal wet-lab data through few-shot learning. *Nature Communications*, *15*(1), 5566.

[14] Minguez, P., Letunic, I., Parca, L., Garcia-Alonso, L., Dopazo, J., Huerta-Cepas, J., & Bork, P. (2015). PTMcode v2: a resource for functional associations of post-translational modifications within and between proteins. *Nucleic acids research*, *43*(D1), D494-D502.

[15] Ochoa, D., Jarnuczak, A. F., Viéitez, C., Gehre, M., Soucheray, M., Mateus, A., ... & Beltrao, P. (2020). The functional landscape of the human phosphoproteome. *Nature biotechnology*, *38*(3), 365-373.
